# Synaptic homeostasis transiently leverages Hebbian mechanisms for a multiphasic response to inactivity

**DOI:** 10.1101/2022.06.18.496642

**Authors:** Simón(e) D. Sun, Daniel Levenstein, Boxing Li, Nataniel Mandelberg, Nicolas Chenouard, Benjamin S. Suutari, Sandrine Sanchez, Guoling Tian, John Rinzel, György Buzsáki, Richard W. Tsien

**Affiliations:** Center for Neural Science, New York University, New York, NY 10003, USA.; Department of Neuroscience and Physiology, Neuroscience Institute, NYU Langone Health, New York, NY 10016, USA; Cold Spring Harbor Laboratory, Cold Spring Harbor, NY 11724; Montreal Neurological Institute, Department of Neurology and Neurosurgery, McGill University, 3810 University Street, Montreal, QC, Canada; Neuroscience Program, Guangdong Provincial Key Laboratory of Brain Function and Disease, Zhongshan School of Medicine and The Fifth Affiliated Hospital, Sun Yat-sen University, Guangzhou, 510810, China; CNRS, Interdisciplinary Institute for Neuroscience, IINS, UMR 5297, University of Bordeaux, F- 33000 Bordeaux, France

**Keywords:** homeostatic plasticity, synaptic homeostasis, Hebbian plasticity, L-type calcium channels, CaMKII, calcineurin, LTP, LTD

## Abstract

Neurons use various forms of negative feedback to maintain their synaptic strengths within an operationally useful range. While this homeostatic plasticity is thought to distinctly counteract the destabilizing positive feedback of Hebbian plasticity, there is considerable overlap in the molecular components mediating both forms of plasticity. The varying kinetics of these components spurs additional inquiry into the dynamics of synaptic homeostasis. We discovered that upscaling of synaptic weights in response to prolonged inactivity is nonmonotonic. Surprisingly, this seemingly oscillatory adaptation involved transient appropriation of molecular effectors associated with Hebbian plasticity, namely CaMKII, L-type Ca^2+^ channels, and Ca^2+^-permeable AMPARs, and homeostatic elements such as calcineurin. We created a dynamic model that shows how traditionally “Hebbian” and “homeostatic” mechanisms can cooperate to autoregulate postsynaptic Ca^2+^ levels. We propose that this combination of mechanisms allows excitatory synapses to adapt to prolonged activity changes and safeguard the capability to undergo future strengthening on demand.

## Introduction

Synaptic plasticity is thought to be a key biological substrate of learning, memory, and neural circuit development. It is theorized that two distinct forms of plasticity – (1) Hebbian positive feedback to strengthen active synapses (Long-term potentiation, LTP) and diminish less active ones (Long-term depression, LTD), and (2) Homeostatic negative feedback to maintain functional stability – enable neurons to integrate recent activity changes without veering into extreme hypo- or hyperactivity. However, despite over a quarter-century of experimental and theoretical study of adaptation to inactivity, its temporal dynamics, molecular underpinnings, and functional implications remain unsettled. As a result, how biochemical and cell biological processes merge into a homeostatic feedback system has not been quantitatively tested.

While Hebbian and homeostatic plasticity are typically conceptualized as distinct processes, the requisite molecular players exhibit considerable overlap (Lee et al., 2014). For example, the calcium (Ca^2+^)/calmodulin (CaM)-dependent kinase CaMKII and phosphatase calcineurin (CaN) are primarily associated with LTP/LTD (Barria et al., 1997; Hell, 2014; Lee et al., 1998; Lisman et al., 2012), but are also involved in homeostatic responses (Arendt et al., 2015; Kim and Ziff, 2014; Thiagarajan et al., 2007). L-type voltage-gated calcium channels (LTCCs), which regulate CaN and CaMKII (Hudmon et al., 2005; Murphy et al., 2014), are involved in both Hebbian (Bauer et al., 2002; Li et al., 2016; Wheeler et al., 2008) and homeostatic (Henry et al., 2018; O’Leary et al., 2010; Slutsky et al., 2004) changes. Such divisions are even murkier when linking AMPA (α-Amino-3-hydroxy-5-methyl-4-isoxazolepropionic acid) receptor (AMPAR) subunits to certain plasticity paradigms (Chater and Goda, 2014; Diering and Huganir, 2018). Calcium-permeable AMPARs (CPARs), while often associated with LTP (Hayashi et al., 2000; Park et al., 2018; Sanderson et al., 2016), are also involved in homeostatic plasticity (Kim and Ziff, 2014; Sanderson et al., 2018), and homeostatic feedback can change the expression levels of several AMPAR subunits: GluA1 (Diering et al., 2014; Soares et al., 2013; Thiagarajan et al., 2005; Wierenga et al., 2005) and GluA2/3 (Gainey et al., 2009; Turrigiano, 2012; Wierenga et al., 2005). Curiously, each of these players operate on different timescales, ranging from seconds (Coultrap and Bayer, 2012; Fujii et al., 2013; Lee et al., 2009) to minutes (Li et al., 2016; Makino and Malinow, 2009) to hours (Adesnik et al., 2005; Henry et al., 2018; Ibata et al., 2004; Schaukowitch et al., 2017). Conceptually distinguishing Hebbian and Homeostatic plasticity requires improved understanding of how overlapping players are differentially engaged when these forms of neuronal plasticity are induced.

In systems with multiple feedback interactions at differing timescales, understanding response dynamics to simple perturbations can reveal how the constitutive mechanisms interact and provide valuable insights into the system’s behavior in complex biologically relevant contexts (Izhikevich, 2007). Theoretical studies generally model homeostatic synaptic plasticity as a control system (O’Leary and Wyllie, 2011; Zenke and Gerstner, 2017) wherein differences in the kinetics of positive or negative feedback components determine response dynamics, which can be monotonic, oscillatory, or unstable (Buonomano, 2005; Harnack et al., 2015; Toyoizumi et al., 2014). Despite theoretical emphasis on the temporal properties of homeostatic adaptations, experimental studies often center on one or two recorded time points after a perturbation in firing rate or synaptic transmission (Kim and Ziff, 2014; Li et al., 2020; Thiagarajan et al., 2005; Zenke et al., 2017). No study so far has focused on the dynamics following a simple intervention such as block of spiking. Understanding these dynamics could reveal a functional logic for the confluence of molecular players employed by homeostatic and Hebbian plasticity.

Here, we show that the response to a widely used homeostatic perturbation, activity silencing by tetrodotoxin (TTX), is a nonmonotonic, near-oscillatory fluctuation involving transient increases of CPARs. These hitherto-overlooked dynamics require an ordered sequence of CaN deactivation (Kim and Ziff, 2014) and CaMKII recruitment and activation (Thiagarajan et al., 2002). We also found that LTCCs tune these dynamics, by studying responses in neurons harboring a Ca_v_1.2 mutation associated with Timothy Syndrome, a rare but highly penetrant form of autism (Bauer et al., 2021; Dick et al., 2016; Splawski et al., 2004). Building on these experimental observations, we propose a quantitative model of synaptic homeostasis that incorporates dynamic interactions between positive and negative feedback signaling. Our findings support the premise that synaptic responsiveness is a variable under feedback control and that spine Ca^2+^/CaM acts as the sensor that recruits “Hebbian” and “homeostatic” elements to stabilize synapses and prepare them for future input. Intriguingly, both our experiments and model provide mechanistic links between synaptic Ca^2+^ homeostasis and activity-dependent regulation of spike width, exemplifying how de-centralized synapses may regulate cell-wide properties.

## Results

### Adaptation to inactivity is a slow non-monotonic response

To delineate the time course of inactivity-induced synaptic homeostasis, we recorded miniature excitatory postsynaptic currents (mEPSCs or minis) from neuronal cortical cultures chronically treated with TTX for 0, 3, 6, 12, 24, 48, and 72 hours (±1.5 h) between 13 to 17 days in vitro (DIV) (n’s: 0 h: 19; 3 h: 7; 6 h: 13; 12 h: 11; 24 h: 17; 48 h: 13; 72 h: 13) and characterized changes in mini amplitude, frequency, and kinetics. Based on previous reports, we expected that the average amplitudes of mEPSCs would increase after 3 h of TTX (Ibata et al., 2008), rise monotonically (Turrigiano and Nelson, 1998, 2004) and persist until 72 h (Ancona Esselmann et al., 2017; O’Brien et al., 1998). To the contrary, the increase in mEPSC amplitude was strikingly non-monotonic, with observed peaks at 6 and 48 h of TTX treatment (One-way ANOVA; F_6,86_ = 8.37, *p<0.0001.* Figure 1C, S1C, Table S2). mEPSC instantaneous frequency also displayed a significant non-monotonic, seemingly oscillatory change with TTX treatment (Figures 1D, S1D, One-way ANOVA F_6,86_ =15.72, *p<0.0001.* Table S2). In addition to the predicted amplitude increase, mEPSCs from TTX-treated neurons also displayed a broad decrease in decay constants (decay **τ**) (Figures S1A&B, Table S2). These data show that upscaling of excitatory synapses during prolonged spike blockade is not monotonic but instead appears to oscillate.

**Figure 1:**
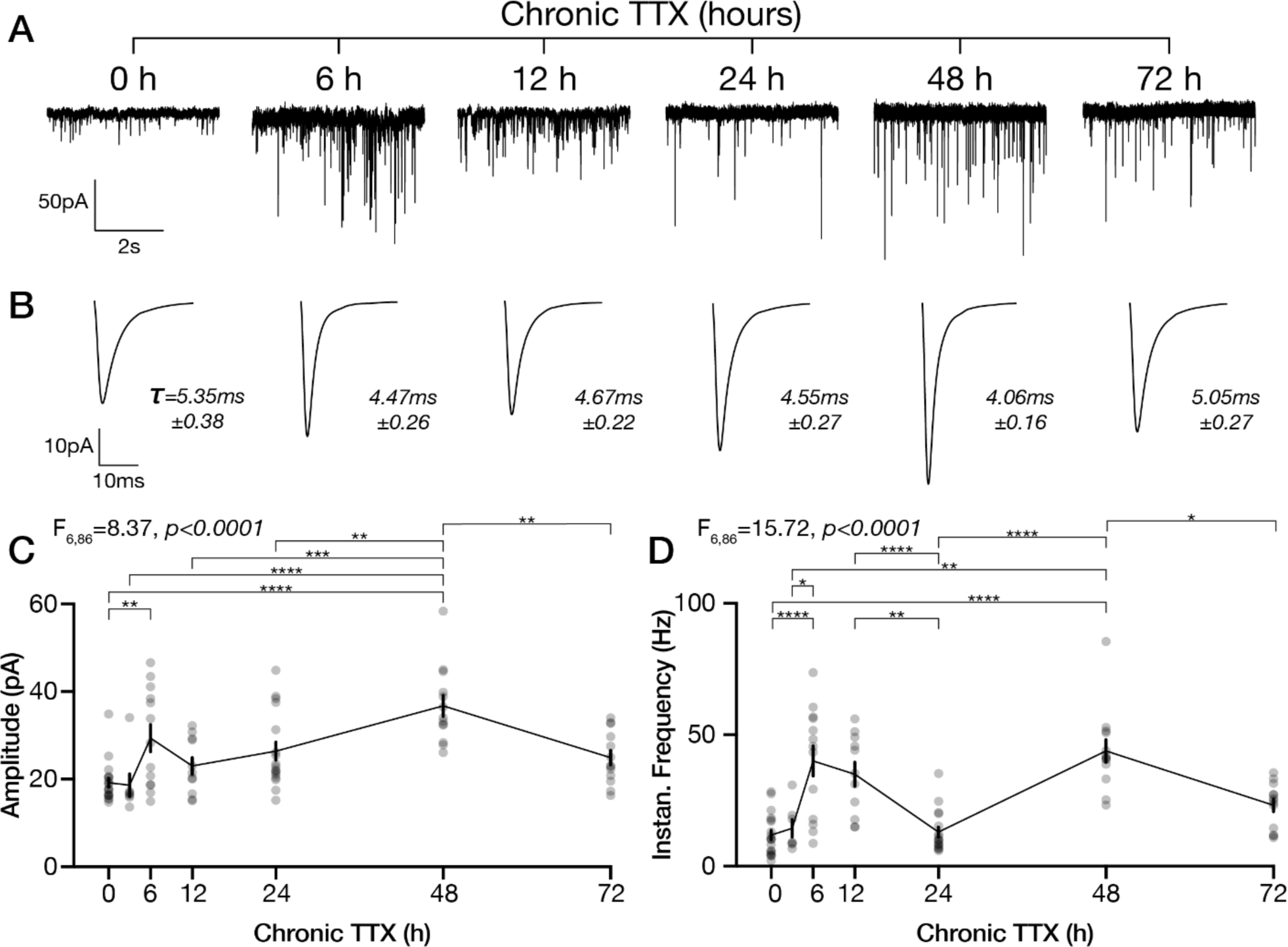
Homeostatic upscaling of synaptic weight exhibits slow non-monotonic changes in mEPSC properties. A. Example voltage clamp recordings of neurons chronically treated with TTX for 0, 6, 12, 24, 48, 72 hours. B. Mean mEPSC waveforms of the mean waveforms and mean decay **τ** of neurons recorded from corresponding TTX timepoints in A. C. Mean±SEM amplitude of mEPSCs, individual cells as gray circles. D. Mean±SEM frequency of mEPSCs. Stars signifying corrected multiple comparison with * *p<0.05*, ** *p<0.01*, *** *p<0.001*, **** *p<0.0001.* See also Figures S1, S2, & S3.

### L-Type Ca^2+^ Channel voltage activation shapes the time course of the homeostatic response

Though voltage-gated calcium channels have been implicated in synaptic strength regulation for over a decade (O’Leary et al., 2010; Thiagarajan et al., 2005), their precise role in inactivity-induced homeostasis remains a mystery. We utilized a mouse model of the Timothy Syndrome (TS2-neo) gain-of-function mutation G406R (Bader et al.. 2011, Bett et al., 2012) to investigate what aspects of L-type channel signaling could be involved in homeostasis and how they might contribute to the dynamic changes we observed. Timothy Syndrome (TiS) is a syndromic form of Autism (Auerbach et al., 2011; Sztainberg and Zoghbi, 2016) that arises from a point mutation in the Ca_v_1.2 calcium channel. This mutation causes a shift in Ca_v_1.2’s voltage-activation curve and faulty inactivation, resulting in channel openings from smaller depolarizations with increased calcium flux when compared to Wild Type (WT) Ca_v_1.2 (Dick et al., 2016; Splawski et al., 2004, 2005). This mutation also leads to exaggerated voltage-dependent conformational signaling (VΔC) and kinase activity during excitation (Li et al., 2016) and plays some role in the homeostatic elevation of spontaneous synaptic Ca^2+^-transients (Li et al., 2020). This led us to hypothesize that the TiS mutation would affect the overall magnitude of synaptic homeostasis.

Surprisingly, mEPSCs from littermatched-TiS cultures exhibited similarly non-monotonic upscaling as seen in their WT counterparts (Figures 2 & S2) but with differing phases in the oscillatory-like changes in amplitudes (Figure 2, TiS in red, WT in gray from Fig. 1; TiS *n’s*: 0 h: 17; 3 h: 7; 6 h: 5; 12 h: 11; 24 h: 14; 48 h: 11; 72 h: 12). Timepoints of peak amplitude in TiS significantly differed from WT at timepoints of 12, 24, and 48 h (Figure 2C, black stars, Two-Way ANOVA, F_6,156_=4.959, *p=0.001*; Table S3). Within TiS cultures, there was a significant change in amplitude at 12, 24, and 72 h of TTX (Figure 2C & S2C red stars, Hours TTX F_6,156_ = 9.610, *p<0.0001*; Table S3A) and in frequency at 6, 12, 24, and 48h of TTX (Figure 2D & S2D red stars, Hours TTX F_6,156_=15.19, *p<0.0001*. Table S3B). We did not observe a phase alteration of mEPSC frequency (F_1, 156_=0.5204, *p*=0.4717) or for mEPSC decay time constants (F_1, 156_=0.5481, *p*=0.4602) (Figure S2A & B) between WT and TiS. These results indicate that the voltage activation of Ca_v_1.2 tunes the dynamics – not the magnitude – of excitatory synaptic homeostasis.

**Figure 2:**
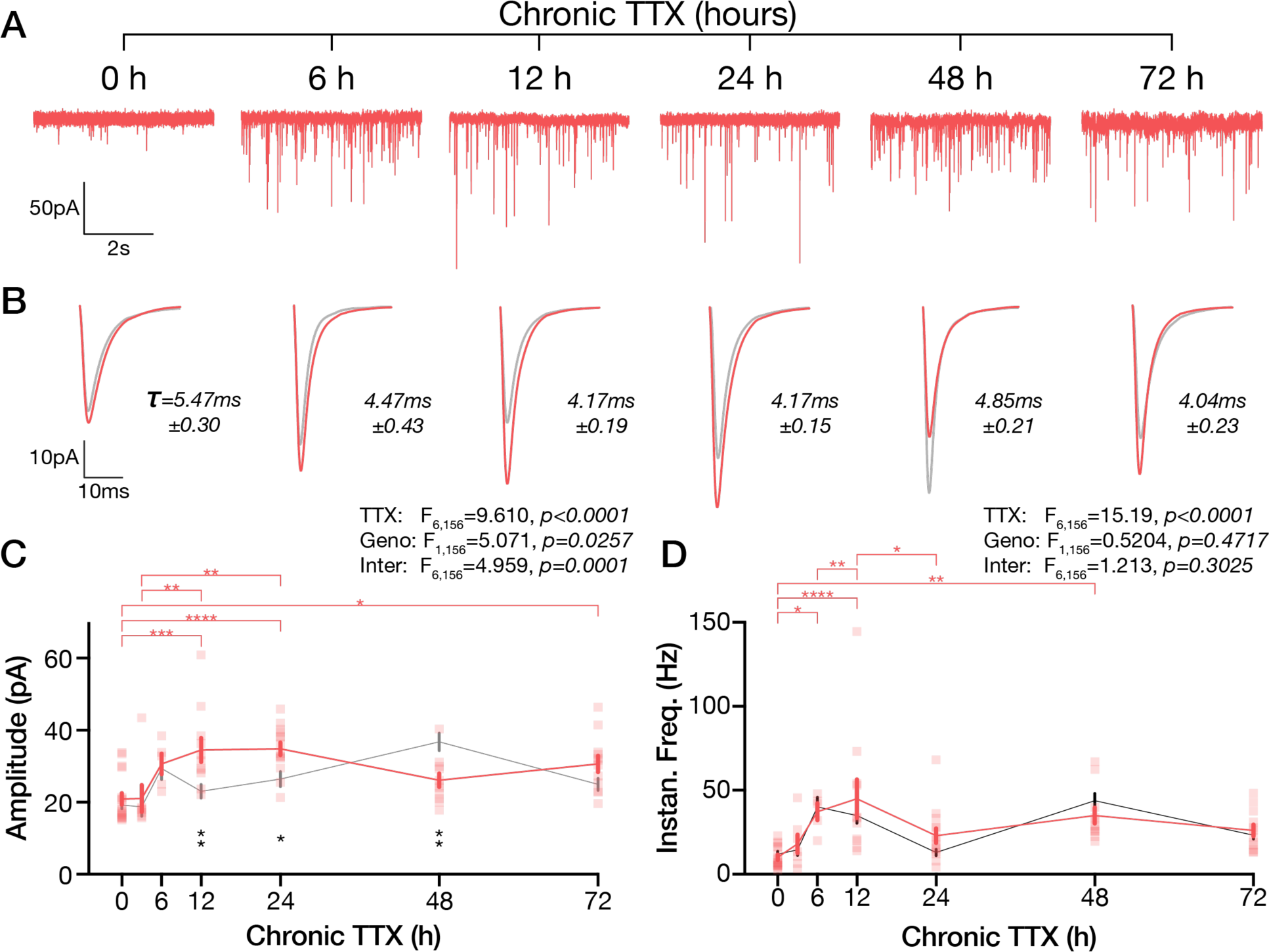
Neurons with the Timothy Syndrome (TiS) Ca_v_1.2 gain-of-function mutation possess an altered homeostatic timecourse. A. Example voltage clamp recordings of TiS neurons chronically treated with TTX for 0, 6, 12, 24, 48, 72 hours. B. Mean mEPSC waveforms of the mean waveforms of neurons recorded in the corresponding TTX timepoint in A. Littermatched WT waveforms from Figure 1 in gray. C. Mean±SEM amplitude of mEPSCs from TiS (red, individual cells as red squares) and littermatched WT (gray from Fig. 1). D. Mean±SEM frequency of mEPSCs from TiS (red) and WT (gray from Fig. 1). Top horizontal red stars indicate corrected significant differences in TiS TTX-treated timepoints. Vertical black stars below indicate corrected significant differences in TTX-treated timepoints across WT and TiS genotype. See also Figure S2, S3.

### mEPSC properties do not developmentally fluctuate

To confirm that our observations were not confounded by developmental or non-homeostatic changes, we recorded mEPSCs from cultures of days *in vitro* (DIV) during which all TTX experiments were conducted (Figure S3, *n’s*: WT: DIV 13: 5, DIV 14: 7, DIV 15: 6, DIV 16: 6, DIV 17: 6; TiS DIV 13: 5, DIV 14: 8, DIV 15: 6, DIV 16: 5, DIV 17: 6). We did not observe significant changes in mEPSC amplitude or frequency 13 to 17 DIV (Figure S3C&E) in either WT or TiS cultures. We observed a small increase in decay kinetics through days *in vitro* (Two Way ANOVA: F_4,52_=5.690, *p=0.0007,* Figure S3G&J). These changes did not fluctuate, in contrast to the changes in mini kinetics seen in homeostatic perturbations, all suggesting that any *in vitro* developmental changes in synaptic and network properties did not confound our measurements and observations of adaptation to activity perturbation.

### GluA1-containing CPARs support synaptic homeostasis at timepoints of peak mEPSC amplitude and fastest decay

We and others have repeatedly observed homeostatic changes in mEPSC decay properties, whereby upscaled mEPSCs decay faster as they increase in amplitude (Kim and Ziff, 2014; Thiagarajan et al., 2005). By comparing the mean mEPSC waveforms at various timepoints from our TTX experiments in both WT and TiS (Figures 1 and 2), we confirmed that mEPSCs with higher amplitudes consistently exhibited faster decays (Figure 3A). Indeed, plotting the mean amplitude (ordinate) against the mean decay **τ** (abscissa) for each timepoint revealed a strong negative correlation (Figure 3B; *r^2^=0.83*). Changes in mEPSC decay time reflects AMPAR composition and the prevalence of GluA1 relative to GluA2: mEPSC events with fast decay times point to a prevalence of calcium-permeable AMPARs (CPARs) (Mosbacher et al., 1994; Thiagarajan et al., 2005). We confirmed that timepoints of highest amplitude and fastest decay to be CPAR dominant by recording from 0, 24, and 48 h TTX-treated cultures while acutely exposing them to philanthotoxin (PhTx), a CPAR specific antagonist. mEPSCs only exhibited PhTx-sensitivity at specific hours of chronic TTX treatment with correlating genotypes. WT cultures treated with TTX for 48 h displayed a PhTx-induced reduction in mEPSC amplitude (Figure 3C bottom right; matched 2-Way ANOVA; n=10; *p=0.0018*. Table S4B) and increase in decay **τ** (Figure S4F; *p=0.0002*. Table S4D). TiS cultures were not PhTx-sensitive at 48 h but were responsive earlier at 24 h instead (Figure 3C middle & S4F. Table S4A & S4C). We noted that mEPSCs from TiS cultures at baseline exhibited slight sensitivity to PhTx, as indicated by an increase in decay **τ** (Figure S4F). While WT neurons exhibited a significant decrease in mEPSC frequency at 24 h TTX + PhTx (Figure S4G), TiS neurons displayed no notable PhTx-induced frequency changes at any timepoint tested (Figure S4E&G).

**Figure 3:**
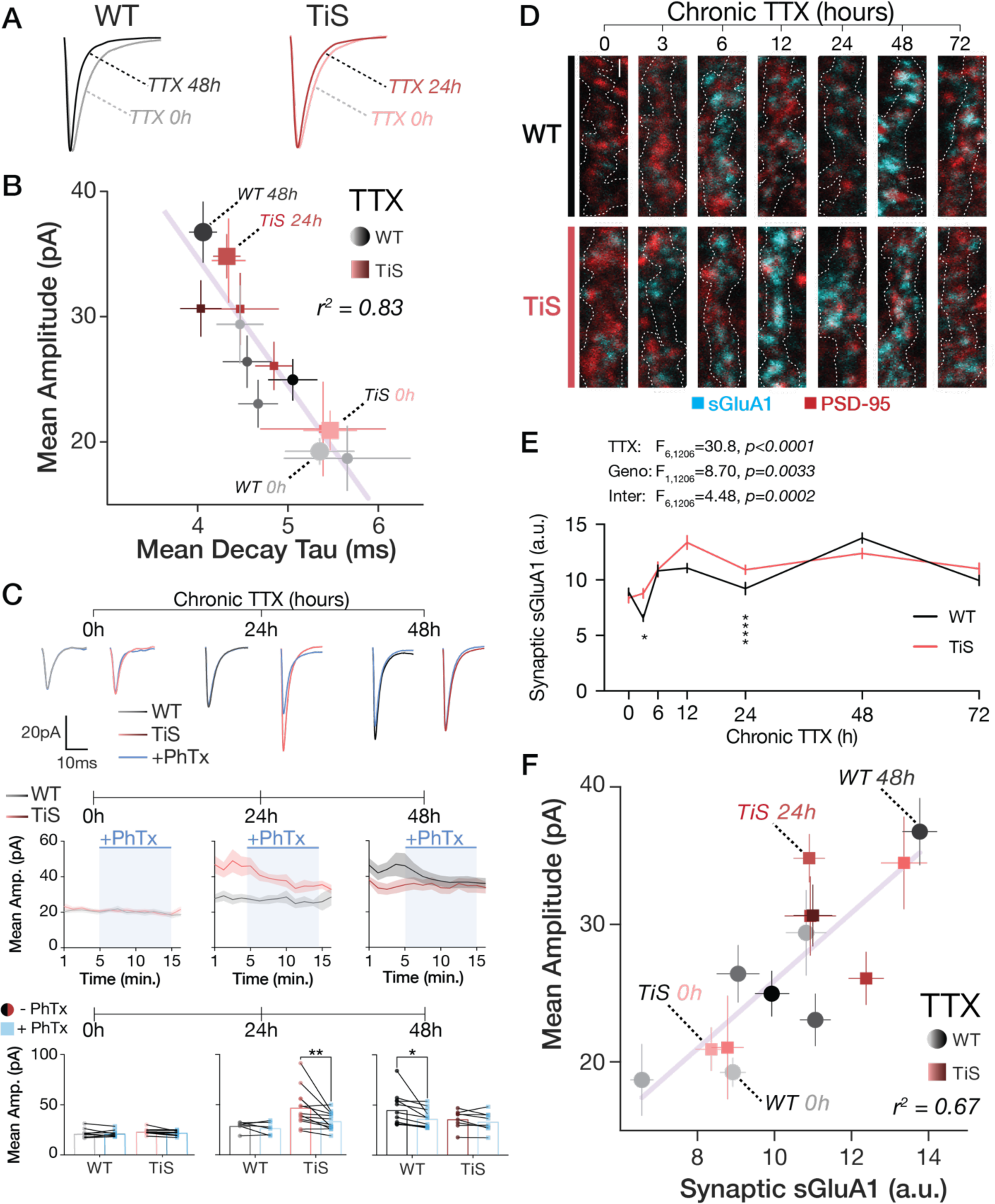
Calcium Permeable GluA1-containing AMPARs transiently contribute to high-amplitude timepoints of synaptic homeostasis. A. Scaled mean mEPSC waveforms from Figs. 1 and 2 comparing the waveform from the highest amplitude timepoints of WT (48 h TTX, black) and TiS (24 h TTX, red) to the corresponding control. B. Mean±SEM amplitude (ordinate) plotted against the Mean±SEM decay (abscissa) of all timepoints (0 – 72 h TTX, light to dark) and genotypes (WT gray to black, TiS pink to maroon). Timepoints selected for PhTx experiment indicated. C. (Top) Mean mEPSC waveforms from voltage clamp recordings with the corresponding TTX timepoint before philanthotoxin (PhTx) wash-on (WT gray to black; TiS pink to maroon) and after PhTx wash-on in blue. (Middle) Timecourse of Mean±SEM mEPSC amplitudes with PhTx wash- on in blue for TTX 0h, 24h, and 48h, for both WT (gray to black) and TiS (pink to maroon). (Bottom) Matched amplitude means from 3 minutes of baseline preceding PhTx wash-on and last 3 minutes of PhTx recording. D. Representative micrographs of surface GluA1 (cyan) and PSD-95 (red) from cortical cultures chronically treated with TTX. Dendritic ROI was determined by MAP2 staining, represented as dotted-white outline. Scale bar 1μm. E. Timecourse of Mean±SEM intensities from (D) for synaptically localized sGluA1colocalization with PSD-95 puncta). F. Mean±SEM amplitude (ordinate) plotted against the Mean±SEM synaptic sGluA1 (abscissa) of all timepoints (0 – 72 h TTX, light to dark) and genotypes (WT gray to black, TiS pink to maroon). Timepoints selected for PhTx experiment indicated. See also Figure S4

We then assessed if there were changes in GluA1 levels during prolonged inactivity by measuring levels of surface GluA1 (sGluA1) with immunofluorescence. We colabeled sGluA1 and PSD-95 in cultures and measured the intensities of the sGluA1 signal colocalized to PSD-95 puncta within dendritic regions, identified by MAP2 immunolabelling (Figure 3D, white outline). We observed a non-monotonic increase in synaptically localized sGluA1 in both WT and TiS cultures (Figure 3E), with significant differences between WT and TiS at 3 and 12 h of chronic TTX (Two-Way ANOVA, TTX: F_6,1206_=30.8, *p<0.0001*, Genotype: F_1,1206_=8.70, *p=0.0033*, Interaction F_6,1206_=4.48, *p=0.0002*. Table S5A). Broadly, TTX treatment increased the intensity of sGluA1 puncta with highest intensity timepoints paralleling the electrophysiological data (Figure 3F, *r^2^=0.67*). We also saw similar changes in shaft sGluA1 (Figure S4J&K. Table S5B), suggesting that synaptic increases were not a consequence of translocation of GluA1. These results indicate that the composition of upscaled synapses changes over the course of homeostatic adaptation by transiently incorporating the GluA1 AMPAR subunit, typically associated with LTP (Diering and Huganir, 2018). Taken together, our results show that dynamic inclusion of GluA1- containing CPARs contributes to increasing synaptic weight in response to prolonged inactivity.

### Chronic Calcineurin blockade mimics early synaptic homeostasis with a monotonic timecourse

The multiphasic homeostatic response suggests that two opposing signaling pathways interact during prolonged inactivity. Based on our findings that Ca_v_1.2 tunes the timecourse of these fluctuations (Figure 2) and that GluA1/CPARs transiently participate (Figure 3), we hypothesized that peaks in mEPSC amplitudes might involve variations in local Ca^2+^-signaling downstream of one or more signaling pathways. A corollary of this hypothesis is that the initial reduction of activity would result in perturbation of a participatory Ca^2+^ signaling enzyme. The Ca^2+^/CaM-dependent phosphatase Calcineurin (CaN) presented as a promising candidate, since it is already implicated in both rapid and prolonged synaptic homeostasis through the action of calcium-permeable AMPARs (CPARs) (Arendt et al., 2015; Kim and Ziff, 2014; Sanderson et al., 2018). We predicted that TTX results in a reduction of CaN activity, via Ca^2+^- signaling through Ca_v_1.2, to mediate early synaptic upscaling. To determine the timecourse of CaN’s involvement, we chronically blocked CaN in WT and TiS cultures with FK506, an inhibitor of CaN activity. In line with our hypothesis, we observed an increase in mEPSC amplitudes by 6 h of chronic FK506 that persisted through 12 h and 24 h in WT and TiS cultures (Figure 4C, S5A&B, Two-way ANOVA: F_4,76_ = 9.68, *p<0.0001*. Table S6A). TiS cultures exhibited a quicker and larger increase in mEPSC amplitudes compared to WT (Figure 4C, S5G&H, F_1,76_=11.38, *p=0.001*. Table S6A). These amplitude increases coincided with drops in decay time constants (FK506: F_4,75_=5.96, *p=0.0003*; Genotype: F_1,75_=9.769, *p=0.0025*. Table S6B) and a rise in frequency (FK506: F_4,75_=3.655, *p=0.0089*; Genotype: F_1,75_=4.27, *p=0.0422*) (Figure 4D&E, S5C-F, S5I-L). Curiously, chronic CaN inhibition in WT cultures did not exhibit a drop of mEPSC amplitudes after 6 h as seen in TTX-treated conditions (Figure 4F), while kinetic properties of the mEPSCs remained comparable (Figure 4G), implying that CaN may be reactivated after an initial response in TTX-induced upscaling.

**Figure 4:**
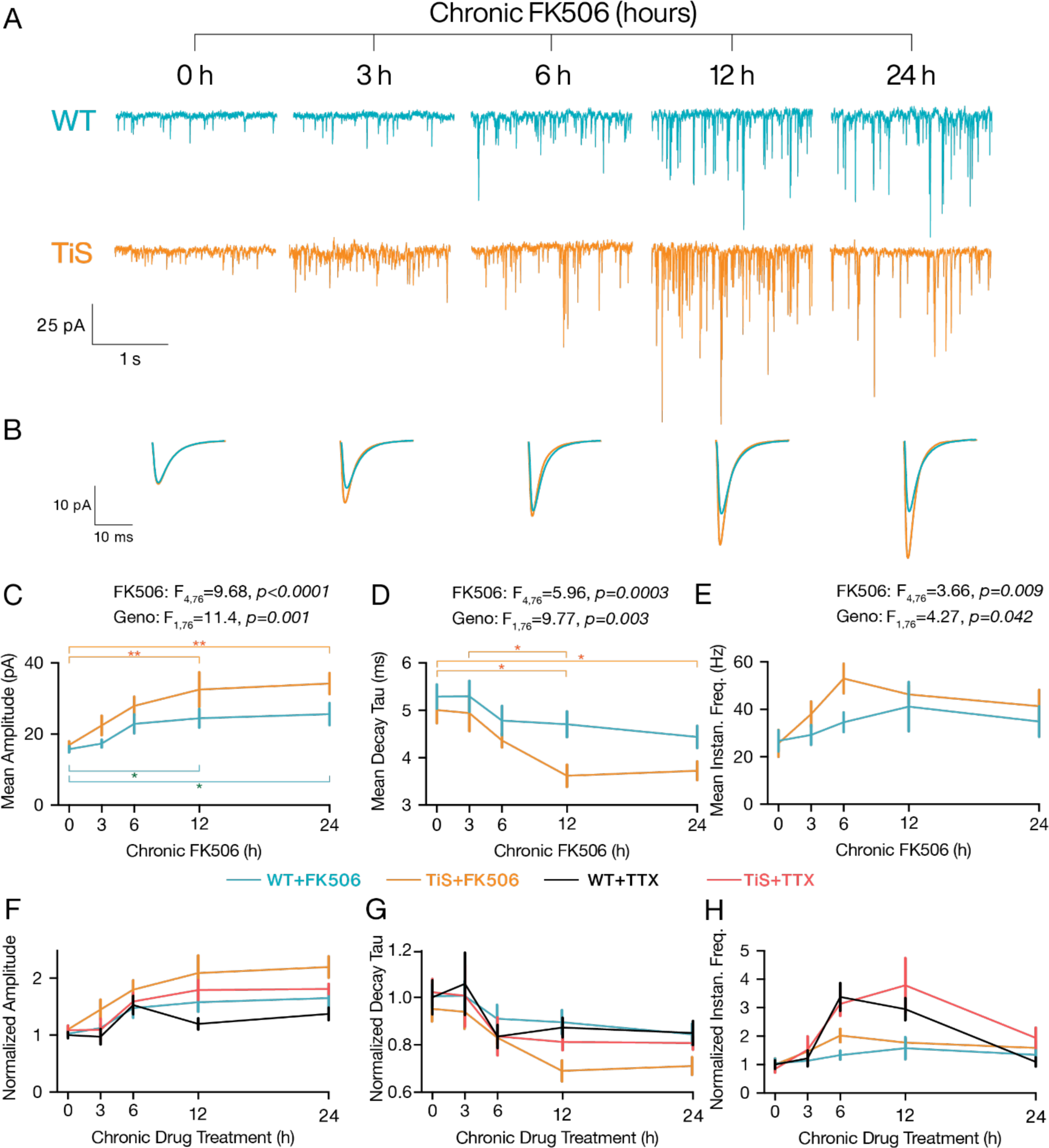
Chronic inactivation of Calcineurin elicits monotonic synaptic upscaling. A. Example voltage clamp recordings of WT (teal) and TiS (orange) neurons chronically treated with CaN inhibitor FK506 for 0, 3, 6, 12, and 24 hours. B. Mean mEPSC waveforms of neurons recorded in the corresponding FK506 timepoint. C. Mean±SEM amplitude of mEPSCs from FK506-treated neurons WT (teal), TiS (orange). D. Mean±SEM decay kinetics of mEPSCs from FK506-treated neurons. E. Mean±SEM instantaneous frequencies of mEPSCs from FK506-treated neurons. F. Normalized Mean±SEM amplitude of mEPSCs from neurons: WT-TTX (black), TiS-TTX (red), WT- FK506 (teal), and TiS-FK506 (orange) over first 24 h of drug treatment. G. Normalized Mean±SEM decay tau of mEPSCs from neurons: WT-TTX (black), TiS-TTX (red), WT- FK506 (teal), and TiS-FK506 (orange) over first 24 h of drug treatment. H. Normalized Mean±SEM instantaneous frequency of mEPSCs from neurons: WT-TTX (black), TiS-TTX (red), WT-FK506 (teal), and TiS-FK506 (orange) over first 24 h of drug treatment. See also Figures S5.

### Transient increase in CaMKII phosphorylation during adaptation to chronic inactivity

Our observations of GluA1 and CPARs increases in both TTX-treated and FK506-treated neurons suggested that upregulation of GluA1 in either condition is driven by another Ca^2+^-dependent player like CaMKII. Furthermore, the monotonic synaptic upscaling from FK506-treatment activity hints that the TTX-induced multiphasic response may result from concerted – but not simultaneous – CaN inactivity and CaMKII activity changes. Since formation of phosphorylated-CaMKII (pCaMKII) is one avenue to CaMKII activation, we colabeled T286/287-phosphorylated CaMKII (pCaMKII) with PSD-95 in TTX-treated cultures in dendritic regions identified by MAP2 immunolabelling. TTX treatment led to dynamic changes in pCaMKII levels in both WT and TiS (Figure 5A&B, S6&7; 2Way ANOVA: hours TTX: F_6,690_ = 7.91, *p<0.0001*; genotype: F_1,690_ = 171.1, *p<0.0001*; interaction: F_6,690_ = 44.86, *p<0.0001*. Table S8A). We discovered a strong transient activation of CaMKII at 6 h of chronic TTX treatment that decreased to an above-baseline level at 12 h of TTX in both WT and TiS cultures. In contrast to other TiS results, the magnitude of pCaMKII at 6 h and 24 h of TTX in TiS neurons was significantly greater than that of WT (Figure 5B, S6A-D & S7A; Sidak-corrected t-tests, 6h and 24h; *p<0.0001.* Table S8A). At other timepoints during when pCaMKII was less active, we still observed smaller but significant differences in its amount of activation over time. These dynamic changes were observed at both putative synapses (PSD-95 colocalized regions, Figure 5B, S5A&C) and putative dendritic shafts (regions not PSD95- colocalized, Figure S7A, S6B&D. Table S8B). These results extend previous observations on synaptic inactivity (Thiagarajan et al., 2002) and provide a dynamic picture of CaMKII regulation in response to spike blockade.

**Figure 5:**
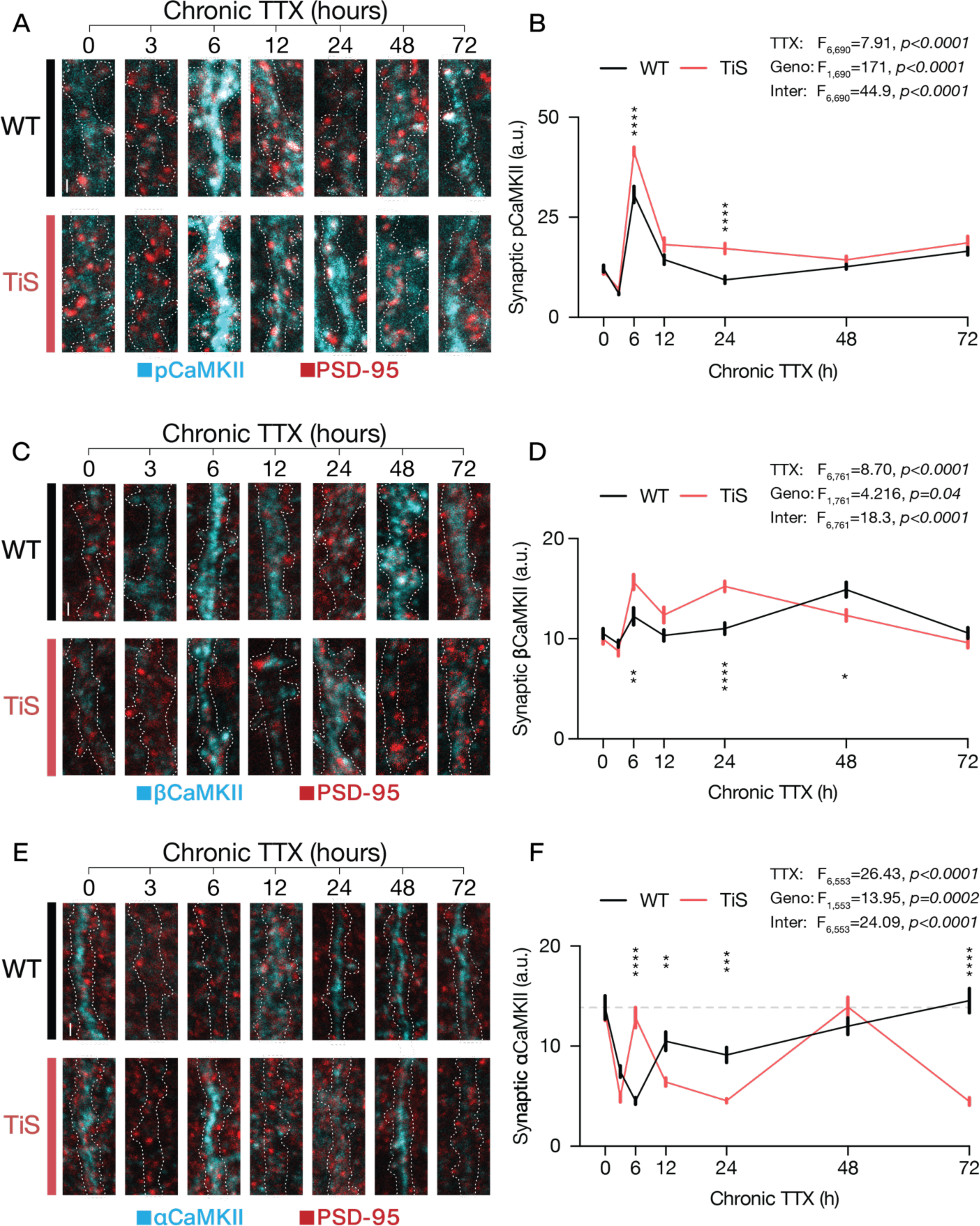
CaMKII activation and isoform expression fluctuates through TTX-induced homeostasis. A. Representative micrographs of pCaMKII (cyan) and PSD-95 (red) from cortical cultures chronically treated with TTX. Dendritic ROI was determined by MAP2 staining, represented as dotted-white outline. B. Timecourse of Mean±SEM intensities from (A) for synaptically localized pCaMKII (colocalization with PSD-95 puncta). C. Representative micrographs of βCaMKII (cyan) and PSD-95 (red) from cortical cultures chronically treated with TTX. Dendritic ROI was determined by MAP2 staining, represented as dotted-white outline. D. Timecourse of Mean±SEM intensities from (C) for synaptically localized βCaMKII (colocalization with PSD-95 puncta). E. Representative micrographs of αCaMKII (cyan) and PSD-95 (red) from cortical cultures chronically treated with TTX. Dendritic ROI was determined by MAP2 staining, represented as dotted-white outline. F. Timecourse of Mean±SEM intensities from (E) for synaptically localized αCaMKII (colocalization with PSD-95 puncta). Scale bars 1μm. See also Figure S6 and S7.

### Synaptic levels of both β and α isoforms of CaMKII fluctuate in response to inactivity

We previously demonstrated that in response to prolonged spike blockade, the balance between CaMKII isoforms shifts from predominantly α to mostly β (Thiagarajan et al., 2002). Because the βCaMKII holoenzyme is ∼8-fold more sensitive to Ca^2+^/calmodulin (Brocke et al., 1999; De Koninck and Schulman, 1998; Miller and Kennedy, 1986), increased β/α could mediate homeostatic regulation of CaMKII calcium sensitivity (Thiagarajan et al., 2002). Thus, we sought to understand how both βCaMKII and αCaMKII changed over time during TTX-induced homeostasis of excitatory synapses. To do so, we co-labeled specific CaMKII isoforms with PSD-95 in separate cultures and measured the intensities of the CaMKII signal colocalized to PSD-95 puncta within dendritic regions, identified by MAP2 immunolabelling.

For synaptic βCaMKII levels, we found a general increase after 6 h of chronic TTX treatment (Figure 5C, Two-way ANOVA hours TTX: F_6,761_=8.70 *p<0.0001*), with similar non-monotonic changes in βCaMKII levels in the dendritic shaft (Figure S7C). These increases differed between WT and TiS (Two-way ANOVA, Genotype: F_1,761_=4.216, *p=0.04*; interaction: F_6,761_=18.3, *p<0.0001*) (Figure 5C&D, S6E-H). βCaMKII levels seemingly oscillate in WT neurons, with elevations of synaptic βCaMKII mirroring that of mEPSC amplitude in showing increases peaking at 6 and 48 h TTX treatment (Figure 5C&D, S6E-H, Table S8C). In TiS neurons, synaptic levels of βCaMKII quickly rose by 6 h of TTX, followed by a slight fluctuation before returning to baseline levels at 72 h. Notably, in WT the mean mEPSC amplitude was linearly correlated with the βCaMKII level at the same timepoint (Figure S7D, *r^2^=0.8975*); the correlation was weaker in TiS cultures (Figure S7E, *r^2^=0.4512*). The elevation of βCaMKII levels and the transient activation of CaMKII suggest that LTP mechanisms are used in low-activity scenarios through an increased availability of the βCaMKII isoform.

For αCaMKII, we observed broad nonmonotonic decreases in synaptic and dendritic shaft levels in response to TTX (Figure 5E&F, S6I-L and S7C, Two-way ANOVA hours TTX: F_6,553_=26.43, *p<0.0001*) and significantly differed between WT and TiS (Two-way ANOVA, genotype: F_1,553_=13.95, *p=0.0002*; interaction: F_6,553_=24.09, *p<0.0001*). αCaMKII levels in WT significantly dropped after 3 h of TTX, reaching a nadir by 6 h before recovering at 72 h (Figure 5F, Table S8E). In contrast, αCaMKII levels in TiS neurons oscillated, with significant troughs at 3, 24, and 72 h of TTX treatment (Table S8E). Between WT and TiS, αCaMKII levels differed at 6, 12, 24, and 72 h (Tukey corrected t-tests), suggesting that the dynamics of αCaMKII isoform expression in response to inactivity were affected by the Ca_v_1.2 mutation. Our data confirm that α- and βCaMKII levels change in opposite directions in response to inactivity (Thiagarajan et al., 2002). Increases in βCaMKII occur with slow (multi-hour) kinetics, consistent with a series of steps such as activation of transcription, translation, and intermediate steps of mRNA/protein transport (Burgin et al., 1990).

### Integrating negative and positive feedback via phosphatase and kinase actions captures key features of homeostatic response

Our results thus far show that TTX-induced inactivity leads to seemingly oscillatory changes in mEPSC properties (Figures 1 & 2) and demonstrate the involvement of CaMKII and CaN activity (Figures 4 & 5). We wished to determine if the interaction of these signaling pathways is suficient to produce a multiphasic homeostatic response, using a reduced model of calcium-dependent phosphorylation of GluA1 (Figure 6, also see model supplement). In the model, postsynaptic Ca^2+^ is determined by the presynaptic quantal rate, *R*, via the equation

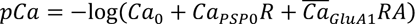

where *pCa* is the negative log concentration of postsynaptic calcium available to be bound to calmodulin; in linear units, *Ca*_0_ is a baseline level of calcium, 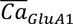 is the maximal calcium per quantal rate through calcium-permeable AMPARs (CPARs), and *Ca_psp0_* is the calcium per quantal rate through CPAR-independent sources. The postsynaptically functional fraction of CPARs, *A*, is determined by a kinetic equation in which *k_f_* and *k_d_* represent the rate of GluA1 phosphorylation and dephosphorylation.

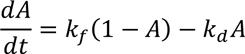

**Figure 6:**
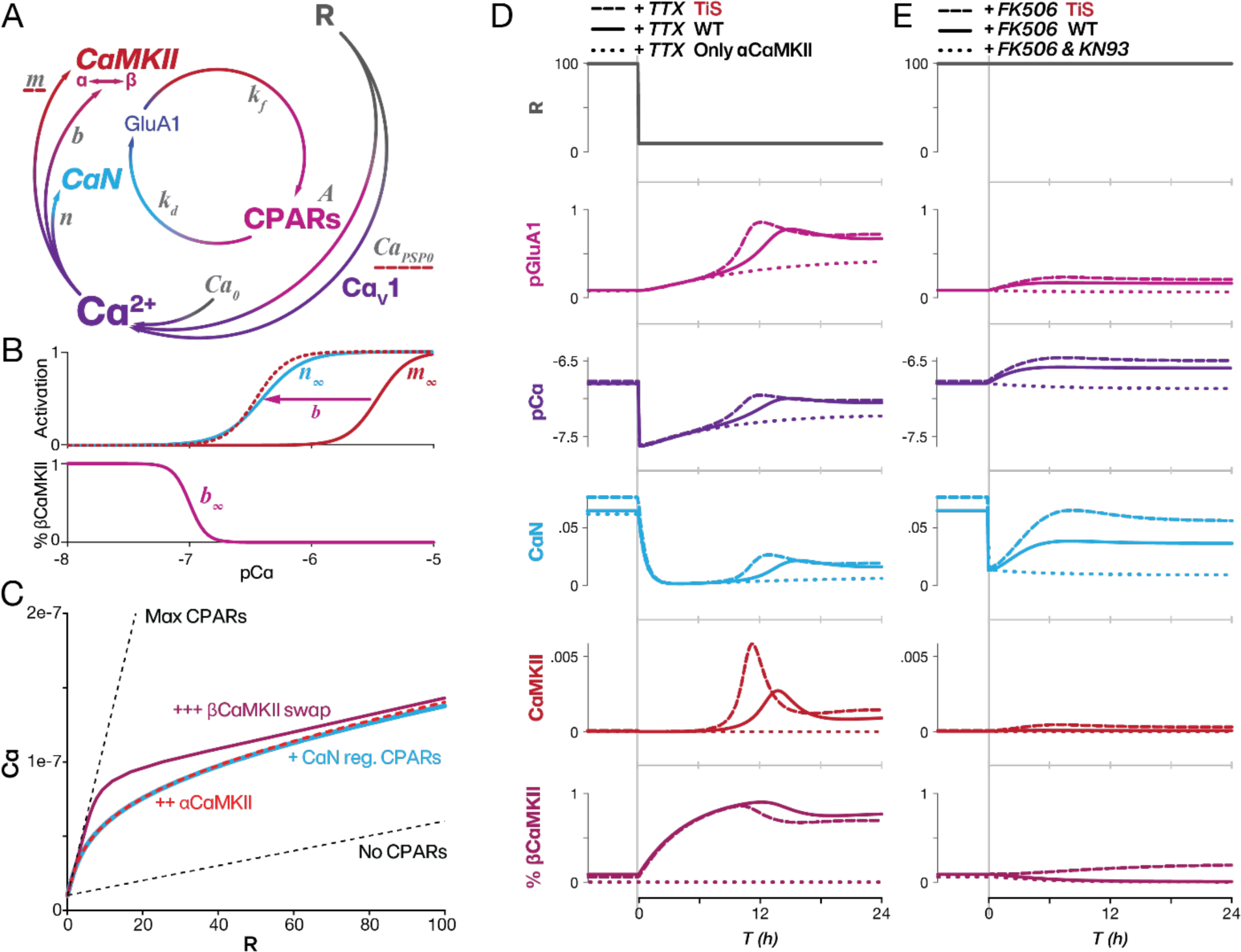
Integrating Ca^2+^-dependent phosphorylation feedback captures key features of homeostatic response. A. Schematic of signaling players (colors) with related mathematical variables (gray). Variables modified to model TiS G406R mutation with red dashed underline. B. Activation curves of 4 (CaN, blue), 3 (100% αCaMKII solid red, 100% βCaMKII dotted red), and in bottom 5, signifying CaMKII ratio *b = β(α + β*), also known as % βCaMKII. Subscript ‘∞’ denotes steady-state value throughout. C. *Ca*(*R*) curve, the steady-state level of calcium as a function of quantal rate (*R*). Full range of steady states exhibited by the model when *A* = 0 and *A* = 1 (black dotted lines). For the cases of CaN-homeostasis only (*k_caMKII_* = 0, light blue), for CaN-homeostasis with αCaMKII only (*Ca*_Δ*b*_ = 0, red dotted), and the full model with switchover from αCaMKII to bCaMKII (maroon). D. Response of the model to the TTX manipulation without presynaptic oscillation by clamping presynaptic quantal release from a baseline level of 100 Hz to “TTX” level of 10 Hz. The response of the model with WT (solid lines), TiS (dashed), and model without α to βCaMKII switch (dotted) for pGluA1 (magenta), pCa (purple), CaN (blue), CaMKII (red), and % βCaMKII (maroon). A. E. Response of the model to CaN inhibition with FK506 with WT parameters (solid lines) and TiS parameters (dashed lines), simulated by reducing CaN eficacy to 20% while maintaining quantal rate (*R* = 100*Hz*); this mimics experimental observation of a monotonic rise in mEPSC amplitudes. Predicted response of the model to FK506 and KN93 (dotted lines) by reducing CaN and CaMKII eficacy while maintaining quantal rate; this results in no change in pGluA1 levels. See also Figure S8 and model supplement.

The phosphorylation rates (*k_f_*, *k_d_*) (Equation S1) depend on activation variables *m, n*, which represent the activation of CaMKII and CaN (Brocke et al., 1999; Stemmer and Klee, 1994) respectively, and 5, which represents the proportion of CaMKII that is βCaMKII and thus determines the calcium sensitivity of CaMKII (Brocke et al., 1999; Thiagarajan et al., 2002). In turn, 3, 4, and 5 are determined by the level of calcium and follow first order kinetics with exponential time constants of 1 min, 40 min, and 300 min, respectively. The model is thus composed of 4 dynamical variables that represent the activity of CaN, CaMKII and CPARs, their relationship with postsynaptic calcium, and the influence of presynaptic rate (Figure 6B; see model supplement and Table S1 for further interpretation of model equations, parameters, and variables).

To understand the relative contribution of the various phosphokinetic components, we first consider their effect on the model’s steady state response as a function of quantal rate (Figure 6C, Methods). Without CPARs (*A* = 0), calcium increases linearly with *R*, reflecting *Ca_PSP0_*. When the proportion of functional CPARs is clamped at *A* = 1, the slope of this relationship increases dramatically. If *A* is allowed to vary according to CaN’s phosphatase activity, the operating curve (light blue) swings between the extremes delineated by fully-functional and non-functional CPARs. This highlights the impact of Ca^2+^/CaM regulation of CaN, which buffers the dependence on rate by adjusting CPAR-mediated Ca^2+^ entry per quantum. Incorporation of the influence of αCaMKII (dashed red line) has little effect on the operating relationship because CaMKII is negligibly recruited at near basal Ca^2+^ levels, while at high quantal rate the activation of αCaMKII results in bistability between a high-[Ca^2+^] CaMKII-active and low- [Ca^2+^] CaMKII-inactive state (Figures S8A & B). In contrast, the calcium-dependent replacement of αCaMKII with βCaMKII has a significant impact on basal Ca^2+^ (solid maroon line), boosting the steady state calcium curve at low rates and flattening it further. In combining the negative feedback effects of CaN deactivation and βCaMKII, this steady-state relationship allows [Ca^2+^] to increase by only ∼40% with a 5-fold variation in quantal rate (e.g. between 20 and 100 Hz).

To understand dynamic changes in [Ca^2+^] and glutamate receptor level following spike blockade, we turned next to modeling time-dependent changes in *pCa* and *A* that follow a sudden change in *R* (Figure 6D). To mimic the application of TTX, we reduced *R* from a “baseline” rate of 100 Hz that incorporates quantal release due to spontaneous action potential firing, to a “TTX” quantal rate of 10 Hz, comparable our data (Figure S8A). The sharp decrease in rate caused a rapid shift in *pCa* from 6.9 to 7.6, and a corresponding decrease in CaN activity. This in turn leads to a slow rise in pGluA1, driven by the basal level of protein kinase activity that brings *pCa* back towards steady-state levels. Simultaneously, the altered *pCa* caused a rise in βCaMKII, which eventually shifted the Ca^2+^-sensitivity of CaMKII enough to trigger a rapid activation of CaMKII and recruitment of CPAR at ∼8 h; this in turn leads *pCa* to return to near its pre-TTX steady-state level. Thus, the interaction of CaN, CaMKII, and the transition between α- to βCaMKII isoforms is sufficient to account for the first peak in mEPSC amplitude (Figure 1) and the corresponding large peak in synaptic pCaMKII (Figure 5A). This simulates the data observed following 24 h of TTX. The pCaMKII peak was exaggerated in a TiS mutation-like modification of the model (dashed lines, Figure 6D), in which the CPAR-independent calcium permeability and effective calcium sensitivity of CaMKII were increased, mimicking the effect of prolonged LTCC currents (Dick et al., 2016; Splawski et al., 2005) and heightened relocation of CaMKII to dendritic spines (Li et al., 2016). On the other hand, these dynamic features were abrogated if *b* was held fixed at *0* (Figure 6D, dotted lines), highlighting the contribution of the α- to βCaMKII conversion.

In addition to the TTX experiment, we were able to simulate the synaptic response to FK506 application by inhibiting CaN-mediated dephosphorylation. To simulate this with the model, we clamped *R* at its baseline rate and at *t = 0*, reduced CaN efficacy to a small fraction of its basal value. In response, the basal level of protein kinase activity *k_fo_* drove *A* to a new, higher steady state with first order kinetics (Figure 6E, solid line), in line with the monotonic change in quantal amplitude upon exposure to FK506 (Figure 5). In contrast, when we simultaneously reduced CaN activity and CaMKII efficacy as well, the slow rise in *A* was not seen (Figure 6E, dotted line). Additionally, with a TiS mutation-like modification of the model, *A* rose more quickly to a higher steady state in response to FK506 (Figure 6E, dashed line).

We noted that the model did not produce the secondary peak in GluA1 at ∼ 48 h of TTX (Figure 1C). We hypothesized that the longer-timescale phase in mEPSC amplitude and CPARs changes may be a result of fluctuations in presynaptic spontaneous release, dependent on a distinct synaptic signaling pathway (Henry et al., 2018), and indicated by the oscillations in mEPSC frequency in our data (Figure 1D). To test for this possibility, we imposed a damped oscillation in *R* that matched the observed mEPSC frequency changes (Figure 1D, S8C top, see Methods). This robustly produced a slow late peak in pGluA1 in the model (Figure S8C, gray shading), with a large CaMKII peak in the first cycle (Figure S8C, red shading) that occurred earlier, closer to the 6 h peak seen in our data (Figure 5A&B). Taken together with the simulation without a presynaptic oscillation (Figure 6D), this model simulation highlighted three distinguishable stages of the response to prolonged reduction in activity. In the first stage, a rapid drop in calcium results in the inactivation of CaN, which allows GluA1 phosphorylation to rise due to the basal activity of protein kinases, including CaMKII (Figure S8B, blue shading). Second, if the rate remains sufficiently low for a long period, a large and transient peak in CaMKII phosphorylation activity occurs at around 6 to 12 h, mediated by a slow increase in β/α ratio (Figure S8C, red shading). The third stage (Figure S8C, gray shading) is dominated by the slow presynaptic oscillation in *R*. Notably, inclusion of the presynaptic oscillation also brought postsynaptic calcium levels closer to pre-TTX condition than the model response without any presynaptic oscillation. Taken together, these simulations of Ca^2+^-sensitive phosphatase and kinase activities, respectively providing negative and positive feedback, were able to mimic the dynamics of inactivity-dependent synaptic plasticity.

### Blockade of CaMKII prevents FK506-induced upscaling

Our model suggests that the cooperation between CaN and CaMKII might be suficient to upscale excitatory synapses early in response to prolonged activity reduction (Figure 6D&E dotted lines, Figure S8B blue and red shading). To experimentally confirm if indeed CaMKII activity was necessary for this early response, we cotreated WT and TiS cultures with FK506 and KN93 to pharmacologically block CaMKII activation and measured mEPSCs at 0, 3, 6, 12, and 24 h. When the eficacies of CaN and CaMKII were simultaneously reduced we saw no change in mEPSCs, as predicted by our model. Mini amplitudes in both WT and TiS did not increase over a 24 h period (Figure 7A-C, S9A-D, WT purple, TiS magenta) (Two-way ANOVA hours treatment: F_4,70_=0.8523, *p=0.4970*; genotype: F_1,70_=0.0052, *p=0.9427*; n’s 6 to 10 per group). mEPSC decay kinetics did not significantly change (Figure 7D, S9E-H) (Two-way ANOVA hours treatment: F_4,70_=1.601, *p=0.1838*; genotype: F_1,70_=2.524, *p=0.1166*) nor did instantaneous frequencies of mEPSC (Figure 7E, S9I-L) (Two-way ANOVA hours treatment: F_4,70_=1.565, *p=0.1933*; genotype: F_1,70_=0.0110, *p=0.9169*). Thus, our experiments fulfilled the theoretical prediction that synaptic upscaling due to deactivation of CaN requires activity of CaMKII.

**Figure 7:**
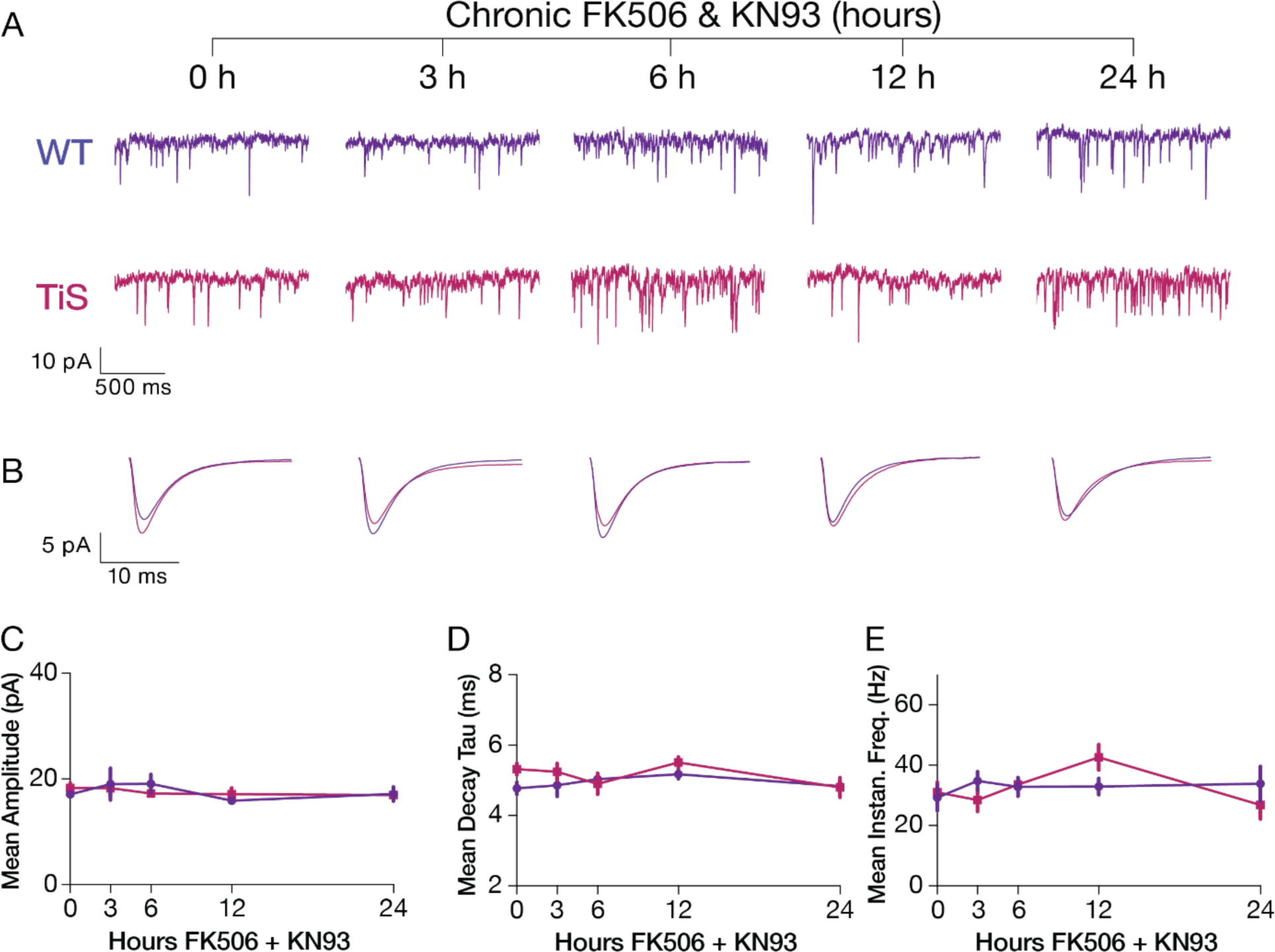
CaMKII blockade prevents FK506-induced upscaling. A. Mean mEPSC waveforms from neurons of corresponding FK506+KN93 timepoint. B. Mean mEPSC waveforms of the mean waveforms of neurons recorded in the corresponding FK506+KN93 timepoint, WT (purple), TiS (magenta). C. Mean±SEM amplitude kinetics of mEPSCs from FK506+KN93 treated neurons, WT (purple), TiS (magenta). D. Mean±SEM instantaneous frequencies of mEPSCs from FK506+KN93 treated neurons. E. Mean±SEM decay kinetics of mEPSCs from FK506+KN93 treated neurons. See also Figure S9.

## Discussion

Synaptic homeostasis is thought to be a slow process that enables gradual readjustment of synaptic transmission while integrating Hebbian changes from prior stimuli. Much is known about both forms of plasticity, but it remains enigmatic how they interact, given their differing dynamics, to enable synaptic flexibility without loss of functional stability (O’Leary and Wyllie, 2011; Toyoizumi et al., 2014; Turrigiano and Nelson, 1998; Zenke and Gerstner, 2017). We unexpectedly discovered that cortical pyramidal neurons respond to prolonged spike blockade with a non-monotonic increase in mEPSC amplitudes (Figures 1 & 2) accompanied by fluctuations in mEPSC decay rate, PhTx sensitivity, GluA1 expression levels (Figure 3) and phosphorylated CaMKII (Figure 5). We determined that this homeostatic fluctuation is a coordinated multiphasic response featuring a transient appropriation of molecular players typically associated with Hebbian plasticity.

Our results indicate that the multiphasic response depends on the sequential change in Ca^2+^-dependent phosphatase and kinase activities (Figure 4, 5, & 6): First, a rapid drop in calcium leads to inactivation of CaN (Figure 4), resulting in increases in phosphorylated GluA1 due to the baseline kinase activity, paralleled by a switchover to βCaMKII expression (Figure 6D, S8C top blue section, Figure 5C-D). Second, a large and transient peak in βCaMKII-mediated CaMKII kinase activity occurs at around 6 to 12 h (Figure 6D, S8C, top red section, Figure 5A-B). Finally, in the model, a slow (∼24 h period) oscillation continues, driven by fluctuation in mEPSC frequency, as calcium settles to a new steady-state (Figure S8C, top gray section). Our experiments verified the necessity of CaMKII for generating the fluctuating response (Figure 5), and the coordinated roles of CaN and CaMKII in generating even the very earliest postsynaptic strengthening (Figure 7).

### Excitatory Synapses engage Hebbian mechanisms to homeostatically regulate Ca^2+^ and preserve synaptic responsiveness

While homeostatic feedback is often invoked as a separate process that counteracts the destabilizing dynamics of Hebbian synaptic strengthening, we found that elements of LTP (CaMKII) are transiently leveraged during homeostatic plasticity, albeit more slowly than in LTP, and interact with more traditional homeostatic elements (Kim and Ziff, 2014; Maghsoodi et al., 2008; Sanderson et al., 2012, 2018; Wang et al., 2011). CaMKII is a long-studied necessary actor in synapse strengthening by AMPAR phosphorylation (Lisman et al., 2012; Malinow and Malenka, 2002). Its activation by Ca^2+^/CaM links CaMKII to synaptic and neuronal activity, with increases in intracellular Ca^2+^ concentration as indicators of elevated activity, activating CaMKII (Herring and Nicoll, 2016). Thus, it appears paradoxical that CaMKII is activated by prolonged *inactivity*, because a reduction in intracellular Ca^2+^ would be expected from the elimination of action potentials (Ibata et al., 2008; Lee and Chung, 2014; Wang et al., 2011). This CaMKII activation coincided with expression of βCaMKII (Figure 5B&D) and depended on it according to isotype-specific knockdown experiments (Groth et al., 2011). Grounded in this data, the model shows how CaMKII activity can be harnessed for homeostatic purposes in addition to its classic Hebbian role.

Why might synapses co-opt classically “Hebbian” mechanisms for homeostasis? First, we propose that this strategy may enable synapses to upregulate postsynaptic calcium more quickly and effectively in response to a decrease in activity. Indeed, when in our model “Hebbian” aspects of CaMKII mobilization were prevented by blocking switchover to the β isoform, the readjustment of calcium was kinetically slower and weaker than with “Hebbian” support (Figure 6D, dotted line). Use of the machinery of Hebbian plasticity provides a transient boost to the dynamics of homeostatic adaptation.

Second, involving CaN and CaMKII (both “Hebbian” elements, and both Ca^2+^/CaM sensitive enzymes) provides a powerful arrangement for spatial and temporal control of intracellular calcium [Ca^2+^]i, thus homeostatically buffering [Ca^2+^]i to prevent cellular dysfunction by excessive Ca^2+^ (Carafoli and Krebs, 2016). In the model, this Ca^2+^ autoregulation is achieved by counterbalancing AMPAR phosphorylation and dephosphorylation and thus Ca^2+^ fluxes via CPARs. Ca^2+^/CaM occupies a central position, downstream of CPAR and LTCC function, yet upstream of phosphatase and kinase activity. There is functional logic to assigning postsynaptic Ca^2+^ the role of the biological variable under homeostatic regulation, as postulated in the classic Marder-Abbott feedback loop (LeMasson et al., 1993; Marder et al., 1996; Siegel et al., 1994). This is a non-trivial choice, insofar as previous schemes considered candidates other than postsynaptic [Ca^2+^]i (Yeung et al., 2004), including AMPAR weight (Turrigiano and Nelson, 1998), firing rate (Hengen et al., 2013, 2016; Torrado Pacheco et al., 2021; Trojanowski et al., 2021), nuclear CaMKIV (Ibata et al., 2008; Joseph and Turrigiano, 2017; Trojanowski and Turrigiano, 2021) and L-type channel activation (Mullins et al., 2016; Thiagarajan et al., 2005).

Third, the autoregulation of the “Hebbian” elements themselves confers the additional benefit of safeguarding the availability of plasticity mechanisms to support synaptic strengthening over seconds and minutes. This rapid strengthening recruits slower remodeling of the postsynaptic machinery involving gene expression and altered enzyme abundance, not just rapid posttranslational modifications. Our model thus integrates earlier work identifying various components of LTP and LTD in homeostatic plasticity (Groth et al., 2011; Lindskog et al., 2010; Purkey and Dell’Acqua, 2020; Thiagarajan et al., 2002, 2005, 2007). The Ca^2+^-dependent expression of the βCaMKII isoform is responsible for calibrating the degree of rate increase necessary to evoke an all-or-none response. This adaptation allows synapses to maintain their responsiveness, as if synaptic responsiveness is under homeostatic regulation and not synaptic weight *per se.* This was probed in the model by clamping the %βCaMKII (*b*) at various levels. The results (FigureS8B) show a change in the properties of the bistability when CaMKII sensitivity adapts to changes in basal [Ca^2+^]i. The shift in Ca^2+^/CaM-sensitivity of CaMKII (Thiagarajan et al., 2002) differs from “metaplasticity”, a sudden and decisive change in the rules governing synaptic plasticity (Abraham and Bear, 1996; Li et al., 2019), but has a “meta” quality in safeguarding responsiveness to an LTP-inducing stimulus. The switchover from αCaMKII to βCaMKII could be framed as a self-regulation of the sensitivity of CaMKII to preserve the inducibility of LTP. We propose that miniature EPSCs co-opt “Hebbian” or “homeostatic” effector mechanisms for local synaptic Ca^2+^ autoregulation at varying activity levels, monitoring synaptic readiness from moment-to-moment and thus continually keeping synapses responsive.

Fourth, the leveraging of “Hebbian” machinery also extends to CaN, which plays a dominant role in long-term synaptic depression (LTD). CaN-driven synaptic homeostasis could be thought of as “de-depression” or removal of a tonic LTD, extending the pattern of using “Hebbian” mechanisms for homeostasis. In both our model and experiments, CaN activity plays a key role in regulating synaptic weight, responding early after a TTX-induced decrease in network activity as seen in the results of Kim & Ziff, 2014. Our experiments provide the additional demonstration that CaN inhibition can elicit a monotonic time course of homeostatic synaptic scaling whose extent is enlarged by a genetic manipulation of LTCC (Figures 2 & 6). These results are in line with previous work elucidating CaN’s role in synaptic homeostasis whether by GluA1 phosphorylation (Kim and Ziff, 2014; Sanderson et al., 2018) or via retinoic acid synthesis (Arendt et al., 2015). The prevention of mEPSC changes with simultaneous CaMKII and CaN blockade in both WT and TiS neurons (Figure 7) further demonstrates joint operation of both limbs of CaV1-related signaling as captured by the model (Figure 6E).

### Ca_v_1.2 channels act as homeostatic effectors: insights from an autism-associated mutation

Our experiments and modeling of the TiS G406R mutation (Bauer et al., 2021; Dick et al., 2016; Splawski et al., 2004, 2005) suggest that Ca_v_1.2 LTCCs act not as sensors of activity changes, but as effectors recruited for homeostatic adaptations. LTCCs are known to interact with multiple postsynaptic elements such as AKAP67/150 (Dittmer et al., 2014; Oliveria et al., 2007), PKA (Diering et al., 2014; Sanderson et al., 2018), CaN (Sanderson et al., 2012; Xu et al., 2010) and CaMKII (Li et al., 2016; Oliveria et al., 2007) and play central roles in signaling synaptic activity to the nucleus (Deisseroth et al., 1996; Greenberg et al., 1986; Li et al., 2016; Ma et al., 2014; Mandelberg, 2020; Morgan and Curran, 1986; Murphy et al., 2014; Wheeler et al., 2008). Their prominent involvement in these processes would allow them to serve as voltage-dependent sensors of activity perturbations (Mullins et al., 2016); indeed, spine LTCCs can be activated by single quantal depolarizations (Li et al., 2020). Thus, we predicted that in the G406R TiS Ca_v_1.2, altered voltage activation, impaired inactivation, increased Ca^2+^ flux (Dick et al., 2016; Splawski et al., 2004, 2005) or exaggerated VΔC (Li et al., 2016) would cause a greater inactivity-induced adaptation relative to WT Ca_v_1.2. We did observe dynamic shifts in CPAR-mediated mEPSCs (Figure 2 & 3) and larger TTX-induced peaks of pCaMKII activation and βCaMKII in spines (Figure 5), indicating that accentuated Ca_v_1.2 VΔC favors local CaMKII accumulation and activation (Li et al., 2016). However, TiS neurons did not exhibit exaggerated mEPSC amplitudes after TTX-exposure, nor were there notable differences observed at baseline (Figure 2). Our experiments showing larger FK506-induced changes in mEPSCs in TiS neurons (Figure 4) and modeling (Figure 6D) suggest that increased pCaMKII activity is counterbalanced by higher basal CaN activity. We interpret these results to indicate that Ca_v_1.2 functions as an effector mechanism engaged by CPAR-mediated depolarizations. When engaged, Ca_v_1.2 channels recruit CaMKII and CaN to restore synaptic Ca^2+^ homeostasis. Future work could determine if other more-rapid forms of plasticity that involve LTCCs, such as LTP/LTD (Bauer et al., 2002; Grover and Teyler, 1990; Oliet et al., 1997; Weisskopf et al., 1999), STDP (Bi and Poo, 1998), or BTSP (Bittner et al., 2017), are affected by alteration of a homeostatic effector.

### Retrograde regulation of presynaptic function extends the non-monotonic response to homeostatic perturbation

Throughout our TTX-induced perturbations, we observed significant fluctuations in mEPSC frequency (Figures 1 & 2) suggesting that presynaptic homeostatic responses are also non-monotonic. CPARs and βCaMKII are known to influence presynaptic adaptations to inactivity via retrograde signaling (Groth et al., 2011; Lindskog et al., 2010; Thiagarajan et al., 2005). There is considerable precedent for retrograde signaling to link postsynaptic interventions to presynaptic modifications of vesicle release, at neuromuscular junctions and central synapses (Davis and Muller, 2015) both in invertebrates (Frank, 2014) and vertebrates (Murthy et al., 2001; Wang et al., 2010). This kind of communication is evident even during prolonged activity perturbations, with complete action potential blockade (Kavalali, 2015).

Thus, it may not be surprising to find evidence for a TTX-induced prolonged oscillation in presynaptic eficacy as monitored by mEPSC frequency, influencing quantal rate, and thus interacting with purely postsynaptic aspects. Indeed, when we remove the presynaptic oscillation in our model, the late stage of the homeostatic response does not occur (Figure 6). LTCC blockade can also engage presynaptic homeostatic plasticity (Thiagarajan et al., 2005) through postsynaptic mTORC1-mediated BDNF signaling (Henry et al., 2012, 2018). This provides a mechanistic scenario for how postsynaptic translational control through mTORC1 (Wang et al., 2021) and retinoic acid (Arendt et al., 2015) might act through Ca^2+^-mediated BDNF release (Groth et al., 2011; Lindskog et al., 2010) to signal back to presynaptic function. Another possible scheme invokes other retrograde messengers such as nitric oxide (Chenouard et al., 2020) produced by nitric oxide synthase, which is also regulated by CaMKII (Araki et al., 2020; Komeima et al., 2000; Watanabe et al., 2003) and CaN (Komeima and Watanabe, 2001; Rameau et al., 2004). Multiple mechanisms could support post- and presynaptic coordination in homeostatic synaptic responses (Tokuoka and Goda, 2008) and impact the response dynamics overall.

### Synaptic Autoregulation Drives Homeostasis of Intrinsic Properties

The scaling of synaptic weights is only one facet of neuronal responses to inactivity. By clarifying the involvement of “Hebbian” components and mechanisms in homeostatic synaptic plasticity - for example, LTCCs, CaMKII, CaN, or CPAR - our findings also illuminate the possible relationship between plasticity of postsynaptic responses and homeostatic adjustment of intrinsic properties such as membrane ionic currents and action potential firing. Our findings align with the suggestion that adaptation of synaptic properties informs activity dependent adaptation of spiking rather than the other way around (Fong et al., 2015). This direction of causation is exemplified by inactivity-driven broadening of action potential duration, which we found to be induced in the absence of spiking, initiated by CPARs and intercepted by blocking CPARs with PhTx (Li et al., 2020; O’Leary et al., 2010). The mechanistic link between changes in postsynaptic properties and neuron-wide changes in intrinsic properties is provided by synapto-nuclear communication, initiated by the very signaling mechanisms - CPAR, LTCCs, CaMKII, or CaN - responsible for the local synaptic homeostasis itself. Ca_v_1.2 also regulates NFAT-mediated transcription via PKA and CaN signaling (Dittmer et al., 2014; Murphy et al., 2014; Oliveria et al., 2007; Sanderson et al., 2018) and CREB transcription factor phosphorylation signals conveyed by γCaMKII (Ma et al., 2014). The common origin of these signals at CaV1 channels supports the idea that CaV1 channels are playing two key roles: reestablishing the local Ca^2+^-dependent kinase/phosphatase balance at the synapse and signaling that activity-state to the nucleus. Together, activity dependent Ca^2+^- autoregulation of many distributed synapses leading to changes in intrinsic properties, may provide substrates for memory encoding (Andersen et al., 2017; Wu et al., 2021), in line with suggestions that memory storage relies on both Hebbian and homeostatic mechanisms (Mullins et al., 2016; Poo et al., 2016)

## Supporting information

Sun et al., Supplemental Materials

## Author Contributions

**Simón(e) Sun:** conceptualization, data curation, formal analysis, funding acquisition, investigation, methodology, validation, visualization, writing – original draft, review, editing. **Daniel Levenstein:** conceptualization, data curation, formal analysis, methodology, software, writing – original draft, review, editing. **Boxing Li:** conceptualization, resources, writing – review, editing. **Nataniel Mandelberg:** formal analysis, writing – review. **Nicolas Chenouard:** data curation, methodology, resources, software, writing –review. **Benjamin Suutari:** data curation, investigation, methodology. **Sandrine Sanchez:** resources, project administration. **Guoling Tian:** resources, project administration. **John Rinzel:** supervision, writing – review. **György Buzsáki:** supervision, writing – review. **Richard Tsien:** conceptualization, funding acquisition, methodology, supervision, validation, visualization, writing – original draft, review, editing.

The authors declare no competing interests.

**Project Funding:** Funding was provided by National Institutes of Health: National Institute of Mental Health through grants **R01MH071739** and **F31MH115611**.

## Acknowledgements

We would like to acknowledge the NYU Langone Animal Research Facility, NYU Langone Microscopy Resources; Samantha Larsen, Simon Chamberland, and other members of the Tsien Lab for stimulating discussion and feedback on the project and manuscript.

## Bibliography

1. Abraham, W.C., and Bear, M.F. (1996). Metaplasticity: the plasticity of synaptic plasticity. Trends Neurosci. 19, 126–130. https://doi.org/10.1016/s0166-2236(96)80018-x.

2. Adesnik, H., Nicoll, R.A., and England, P.M. (2005). Photoinactivation of native AMPA receptors reveals their real-time trafficking. Neuron 48, 977–985. https://doi.org/10.1016/j.neuron.2005.11.030.

3. Ancona Esselmann, S.G., Díaz-Alonso, J., Levy, J.M., Bemben, M.A., and Nicoll, R.A. (2017). Synaptic homeostasis requires the membrane-proximal carboxy tail of GluA2. Proc. Natl. Acad. Sci. U. S. A. 114, 13266–13271.

4. Andersen, N., Krauth, N., and Nabavi, S. (2017). Hebbian plasticity in vivo: relevance and induction. Curr. Opin. Neurobiol. 45, 188–192. https://doi.org/10.1016/j.conb.2017.06.001.

5. Araki, S., Osuka, K., Takata, T., Tsuchiya, Y., and Watanabe, Y. (2020). Coordination between Calcium/Calmodulin-Dependent Protein Kinase II and Neuronal Nitric Oxide Synthase in Neurons. Int. J. Mol. Sci. 21, 7997. https://doi.org/10.3390/ijms21217997.

6. Arendt, K.L., Zhang, Z., Ganesan, S., Hintze, M., Shin, M.M., Tang, Y., Cho, A., Graef, I.A., and Chen, L. (2015). Calcineurin mediates homeostatic synaptic plasticity by regulating retinoic acid synthesis. Proc. Natl. Acad. Sci. U. S. A. 112, E5744–52. https://doi.org/10.1073/pnas.1510239112.

7. Auerbach, B.D., Osterweil, E.K., and Bear, M.F. (2011). Mutations causing syndromic autism define an axis of synaptic pathophysiology. Nature 480, 63. https://doi.org/10.1038/nature10658.

8. Bader, P.L., Faizi, M., Kim, L.H., Owen, S.F., Tadross, M.R., Alfa, R.W., Bett, G.C.L., Tsien, R.W., Rasmusson, R.L., and Shamloo, M. (2011). Mouse model of Timothy syndrome recapitulates triad of autistic traits. Proc. Natl. Acad. Sci. U. S. A. 108, 15432–15437.

9. Barria, A., Muller, D., Derkach, V., Griffith, L.C., and Soderling, T.R. (1997). Regulatory Phosphorylation of AMPA-Type Glutamate Receptors by CaM-KII During Long-Term Potentiation. Science 276, 2042–2045.

10. Bauer, E.P., Schafe, G.E., and LeDoux, J.E. (2002). NMDA receptors and L-type voltage-gated calcium channels contribute to long-term potentiation and different components of fear memory formation in the lateral amygdala. J. Neurosci. 22, 5239–5249. https://doi.org/10.1523/jneurosci.22-12-05239.2002.

11. Bauer, R., Timothy, K.W., and Golden, A. (2021). Update on the Molecular Genetics of Timothy Syndrome. Front Pediatr 9, 668546. https://doi.org/10.3389/fped.2021.668546.

12. Berridge, M.J., Lipp, P., and Bootman, M.D. (2000). The versatility and universality of calcium signalling. Nat. Rev. Mol. Cell Biol. 1, 11–21. https://doi.org/10.1038/35036035.

13. Bett, G.C., Lis, A., Wersinger, S.R., Baizer, J.S., Duffey, M.E., and Rasmusson, R.L. (2012). A Mouse Model of Timothy Syndrome: a Complex Autistic Disorder Resulting from a Point Mutation in Ca_v_1.2. N. Am. J. Med. Sci. 5, 135–140. https://doi.org/10.7156/najms.2012.053135.

14. Bi, G.Q., and Poo, M.M. (1998). Synaptic modifications in cultured hippocampal neurons: dependence on spike timing, synaptic strength, and postsynaptic cell type. J. Neurosci. 18, 10464–10472. https://doi.org/10.1523/jneurosci.18-24-10464.1998.

15. Bittner, K.C., Milstein, A.D., Grienberger, C., Romani, S., and Magee, J.C. (2017). Behavioral time scale synaptic plasticity underlies CA1 place fields. Science 357, 1033–1036. https://doi.org/10.1126/science.aan3846.

16. Brocke, L., Chiang, L.W., Wagner, P.D., and Schulman, H. (1999). Functional implications of the subunit composition of neuronal CaM kinase II. J. Biol. Chem. 274, 22713–22722. https://doi.org/10.1074/jbc.274.32.22713.

17. Buonomano, D.V. (2005). A learning rule for the emergence of stable dynamics and timing in recurrent networks. J. Neurophysiol. 94, 2275–2283. https://doi.org/10.1152/jn.01250.2004.

18. Burgin, K.E., Waxham, M.N., Rickling, S., Westgate, S.A., Mobley, W.C., and Kelly, P.T. (1990). In situ hybridization histochemistry of Ca2+/calmodulin-dependent protein kinase in developing rat brain. J. Neurosci. 10, 1788–1798.

19. Carafoli, E., and Krebs, J. (2016). Why Calcium? How Calcium Became the Best Communicator. J. Biol. Chem. 291, 20849–20857. https://doi.org/10.1074/jbc.R116.735894.

20. Chater, T.E., and Goda, Y. (2014). The role of AMPA receptors in postsynaptic mechanisms of synaptic plasticity. Front. Cell. Neurosci. 8, 401. https://doi.org/10.3389/fncel.2014.0040.

21. Chenouard, N., Xuan, F., and Tsien, R.W. (2020). Synaptic vesicle traffic is supported by transient actin filaments and regulated by PKA and NO. Nat. Commun. 11, 5318. https://doi.org/10.1038/s41467-020-19120-1.

22. Coultrap, S.J., and Bayer, K.U. (2012). CaMKII regulation in information processing and storage. Trends Neurosci. 35, 607–618. https://doi.org/10.1016/j.tins.2012.05.003.

23. Davis, G.W., and Muller, M. (2015). Homeostatic control of presynaptic neurotransmitter release. Annu. Rev. Physiol. 77, 251–270. https://doi.org/10.1146/annurev-physiol-021014-071740.

24. De Koninck, P., and Schulman, H. (1998). Sensitivity of CaM kinase II to the frequency of Ca2+ oscillations. Science 279, 227–230. https://doi.org/10.1126/science.279.5348.227.

25. Deisseroth, K., Bito, H., and Tsien, R.W. (1996). Signaling from synapse to nucleus: postsynaptic CREB phosphorylation during multiple forms of hippocampal synaptic plasticity. Neuron 16, 89–101. https://doi.org/10.1016/s0896-6273(00)80026-4.

26. Dick, I.E., Joshi-Mukherjee, R., Yang, W., and Yue, D.T. (2016). Arrhythmogenesis in Timothy Syndrome is associated with defects in Ca(2+)-dependent inactivation. Nat. Commun. 7, 10370. https://doi.org/10.1038/ncomms10370.

27. Diering, G.H., and Huganir, R.L. (2018). The AMPA Receptor Code of Synaptic Plasticity. Neuron 100, 314–329. https://doi.org/10.1016/j.neuron.2018.10.018.

28. Diering, G.H., Gustina, A.S., and Huganir, R.L. (2014). PKA-GluA1 coupling via AKAP5 controls AMPA receptor phosphorylation and cell-surface targeting during bidirectional homeostatic plasticity. Neuron 84, 790–805. https://doi.org/10.1016/j.neuron.2014.09.024.

29. Dittmer, P.J., Dell’Acqua, M.L., and Sather, W.A. (2014). Ca2+/calcineurin-dependent inactivation of neuronal L-type Ca2+ channels requires priming by AKAP-anchored protein kinase A. Cell Rep. 7, 1410–1416. https://doi.org/10.1016/j.celrep.2014.04.039.

30. Ehlers, M.D. (2003). Activity level controls postsynaptic composition and signaling via the ubiquitin-proteasome system. Nat. Neurosci. 6, 231–242. https://doi.org/10.1038/nn1013.

31. Fong, M.-F., Newman, J.P., Potter, S.M., and Wenner, P. (2015). Upward synaptic scaling is dependent on neurotransmission rather than spiking. Nat. Commun. 6, 6339. https://doi.org/10.1038/ncomms7339.

32. Frank, C.A. (2014). Homeostatic plasticity at the Drosophila neuromuscular junction. Neuropharmacology 78, 63–74. https://doi.org/10.1016/j.neuropharm.2013.06.015.

33. Fujii, H., Inoue, M., Okuno, H., Sano, Y., Takemoto-Kimura, S., Kitamura, K., Kano, M., and Bito, H. (2013). Nonlinear decoding and asymmetric representation of neuronal input information by CaMKIIα and calcineurin. Cell Rep. 3, 978–987.

34. Gainey, M.A., Hurvitz-Wolff, J.R., Lambo, M.E., and Turrigiano, G.G. (2009). Synaptic scaling requires the GluR2 subunit of the AMPA receptor. J. Neurosci. 29, 6479–6489. https://doi.org/10.1523/JNEUROSCI.3753-08.2009.

35. Greenberg, M.E., Ziff, E.B., and Greene, L.A. (1986). Stimulation of neuronal acetylcholine receptors induces rapid gene transcription. Science 234, 80–83.

36. Grienberger, C., and Konnerth, A. (2012). Imaging calcium in neurons. Neuron 73, 862–885. https://doi.org/10.1016/j.neuron.2012.02.011.

37. Groth, R.D., Lindskog, M., Thiagarajan, T.C., Li, L., and Tsien, R.W. (2011). Beta Ca2+/CaM-dependent kinase type II triggers upregulation of GluA1 to coordinate adaptation to synaptic inactivity in hippocampal neurons. Proc. Natl. Acad. Sci. U. S. A. 108, 828–833. https://doi.org/10.1073/pnas.1018022108.

38. Grover, L.M., and Teyler, T.J. (1990). Two components of long-term potentiation induced by different patterns of afferent activation. Nature 347, 477–479. https://doi.org/10.1038/347477a0.

39. Hanson, P.I., Meyer, T., Stryer, L., and Schulman, H. (1994). Dual role of calmodulin in autophosphorylation of multifunctional cam kinase may underlie decoding of calcium signals. Neuron 12, 943–956. https://doi.org/10.1016/0896-6273(94)90306-9.

40. Harnack, D., Pelko, M., Chaillet, A., Chitour, Y., and van Rossum, M.C.W. (2015). Stability of Neuronal Networks with Homeostatic Regulation. PLoS Comput. Biol. 11, e1004357. https://doi.org/10.1371/journal.pcbi.1004357.

41. Hayashi, Y., Shi, S.H., Esteban, J.A., Piccini, A., Poncer, J.C., and Malinow, R. (2000). Driving AMPA receptors into synapses by LTP and CaMKII: requirement for GluR1 and PDZ domain interaction. Science 287, 2262– 2267. https://doi.org/10.1126/science.287.5461.2262.

42. Hell, J.W. (2014). CaMKII: claiming center stage in postsynaptic function and organization. Neuron 81, 249–265. https://doi.org/10.1016/j.neuron.2013.12.024.

43. Hengen, K.B., Lambo, M.E., Van Hooser, S.D., Katz, D.B., and Turrigiano, G.G. (2013). Firing Rate Homeostasis in Visual Cortex of Freely Behaving Rodents. Neuron 80, 335–342. https://doi.org/10.1016/j.neuron.2013.08.038.

44. Hengen, K.B., Torrado Pacheco, A., McGregor, J.N., Van Hooser, S.D., and Turrigiano, G.G. (2016). Neuronal Firing Rate Homeostasis Is Inhibited by Sleep and Promoted by Wake. Cell 165, 180–191. https://doi.org/10.1016/j.cell.2016.01.046.

45. Henry, F.E., McCartney, A.J., Neely, R., Perez, A.S., Carruthers, C.J.L., Stuenkel, E.L., Inoki, K., and Sutton, M.A. (2012). Retrograde changes in presynaptic function driven by dendritic mTORC1. J. Neurosci. 32, 17128–17142 https://doi.org/10.1523/JNEUROSCI.2149-12.2012.

46. Henry, F.E., Wang, X., Serrano, D., Perez, A.S., Carruthers, C.J.L., Stuenkel, E.L., and Sutton, M.A. (2018). A Unique Homeostatic Signaling Pathway Links Synaptic Inactivity to Postsynaptic mTORC1. Journal of Neuroscience 38, 2207–2225. https://doi.org/10.1523/jneurosci.1843-17.2017.

47. Herring, B.E., and Nicoll, R.A. (2016). Long-Term Potentiation: From CaMKII to AMPA Receptor Trafficking. Annu. Rev. Physiol. 78, 351–365. https://doi.org/10.1146/annurev-physiol-021014-071753.

48. Hubbard, M.J., and Klee, C.B. (1987). Calmodulin binding by calcineurin. Ligand-induced renaturation of protein immobilized on nitrocellulose. J. Biol. Chem. 262, 15062–15070.

49. Hudmon, A., Schulman, H., Kim, J., Maltez, J.M., Tsien, R.W., and Pitt, G.S. (2005). CaMKII tethers to L-type Ca2+ channels, establishing a local and dedicated integrator of Ca2+ signals for facilitation. J. Cell Biol. 171, 537–547. https://doi.org/10.1083/jcb.200505155.

50. Ibata, K., Sun, Q., and Turrigiano, G.G. (2008). Rapid synaptic scaling induced by changes in postsynaptic firing. Neuron 57, 819–826. https://doi.org/10.1016/j.neuron.2008.02.031.

51. Izhikevich, E.M. (2007). Dynamical Systems in Neuroscience (MIT Press).

52. Joseph, A., and Turrigiano, G.G. (2017). All for One But Not One for All: Excitatory Synaptic Scaling and Intrinsic Excitability Are Coregulated by CaMKIV, Whereas Inhibitory Synaptic Scaling Is Under Independent Control. J. Neurosci. 37, 6778–6785. https://doi.org/10.1523/JNEUROSCI.0618-17.2017.

53. Kavalali, E.T. (2015). The mechanisms and functions of spontaneous neurotransmitter release. Nat. Rev. Neurosci. 16, 5–16. https://doi.org/10.1038/nrn3875.

54. Kim, S., and Ziff, E.B. (2014). Calcineurin mediates synaptic scaling via synaptic trafficking of Ca2+-permeable AMPA receptors. PLoS Biol. 12, e1001900. https://doi.org/10.1371/journal.pbio.1001900.

55. Komeima, K., and Watanabe, Y. (2001). Dephosphorylation of nNOS at Ser(847) by protein phosphatase 2A. FEBS Lett. 497, 65–66. https://doi.org/10.1016/s0014-5793(01)02389-4.

56. Komeima, K., Hayashi, Y., Naito, Y., and Watanabe, Y. (2000). Inhibition of Neuronal Nitric-oxide Synthase by Calcium/ Calmodulin-dependent Protein Kinase IIα through Ser847 Phosphorylation in NG108-15 Neuronal Cells. J. Biol. Chem. 275, 28139–28143. https://doi.org/10.1074/jbc.m003198200.

57. Lee, K.F.H., Soares, C., and Béïque, J.-C. (2014). Tuning into diversity of homeostatic synaptic plasticity. Neuropharmacology 78, 31–37.

58. Lee, K.Y., and Chung, H.J. (2014). NMDA receptors and L-type voltage-gated Ca^2+^ channels mediate the expression of bidirectional homeostatic intrinsic plasticity in cultured hippocampal neurons. Neuroscience 277, 610–623. https://doi.org/10.1016/j.neuroscience.2014.07.038.

59. Lee, H.K., Kameyama, K., Huganir, R.L., and Bear, M.F. (1998). NMDA induces long-term synaptic depression and dephosphorylation of the GluR1 subunit of AMPA receptors in hippocampus. Neuron 21, 1151–1162. https://doi.org/10.1016/s0896-6273(00)80632-7.

60. Lee, S.-J.R., Escobedo-Lozoya, Y., Szatmari, E.M., and Yasuda, R. (2009). Activation of CaMKII in single dendritic spines during long-term potentiation. Nature 458, 299–304.

61. LeMasson, G., Marder, E., and Abbott, L.F. (1993). Activity-dependent regulation of conductances in model neurons. Science 259, 1915–1917. https://doi.org/10.1126/science.8456317.

62. Li, B., Tadross, M.R., and Tsien, R.W. (2016). Sequential ionic and conformational signaling by calcium channels drives neuronal gene expression. Science 351, 863–867. https://doi.org/10.1126/science.aad3647.

63. Li, B., Suutari, B.S., Sun, S.D., Luo, Z., Wei, C., Chenouard, N., Mandelberg, N.J., Zhang, G., Wamsley, B., Tian, G., et al. (2020). Neuronal Inactivity Co-opts LTP Machinery to Drive Potassium Channel Splicing and Homeostatic Spike Widening. Cell 181, 1547–1565.e15. https://doi.org/10.1016/j.cell.2020.05.013.

64. Li, J., Park, E., Zhong, L.R., and Chen, L. (2019). Homeostatic synaptic plasticity as a metaplasticity mechanism - a molecular and cellular perspective. Curr. Opin. Neurobiol. 54, 44–53. https://doi.org/10.1016/j.conb.2018.08.010.

65. Lindskog, M., Li, L., Groth, R.D., Poburko, D., Thiagarajan, T.C., Han, X., and Tsien, R.W. (2010). Postsynaptic GluA1 enables acute retrograde enhancement of presynaptic function to coordinate adaptation to synaptic inactivity. Proc. Natl. Acad. Sci. U. S. A. 107, 21806–21811. https://doi.org/10.1073/pnas.1016399107.

66. Lisman, J., Yasuda, R., and Raghavachari, S. (2012). Mechanisms of CaMKII action in long-term potentiation. Nat. Rev. Neurosci. 13, 169–182. https://doi.org/10.1038/nrn3192.

67. Ma, H., Groth, R.D., Cohen, S.M., Emery, J.F., Li, B., Hoedt, E., Zhang, G., Neubert, T.A., and Tsien, R.W. (2014). “aMKII shuttles Ca^2+^/CaM to the nucleus to trigger CREB phosphorylation and gene expression. Cell 159, 281–294. https://doi.org/10.1016/j.cell.2014.09.019.

68. Maghsoodi, B., Poon, M.M., Nam, C.I., Aoto, J., Ting, P., and Chen, L. (2008). Retinoic acid regulates RARalpha-mediated control of translation in dendritic RNA granules during homeostatic synaptic plasticity. Proc. Natl. Acad. Sci. U. S. A. 105, 16015–16020. https://doi.org/10.1073/pnas.0804801105.

69. Makino, H., and Malinow, R. (2009). AMPA receptor incorporation into synapses during LTP: the role of lateral movement and exocytosis. Neuron 64, 381–390. https://doi.org/10.1016/j.neuron.2009.08.035.

70. Malinow, R., and Malenka, R.C. (2002). AMPA receptor trafficking and synaptic plasticity. Annu. Rev. Neurosci. 25, 103–126. https://doi.org/10.1146/annurev.neuro.25.112701.142758.

71. Mandelberg, N.J. (2020). L-Type Calcium Channels Cooperate with NMDA Receptors to Signal to the Nucleus from Dendritic Spines.

72. Marder, E., Abbott, L.F., Turrigiano, G.G., Liu, Z., and Golowasch, J. (1996). Memory from the dynamics of intrinsic membrane currents. Proc. Natl. Acad. Sci. U. S. A. 93, 13481–13486. https://doi.org/10.1073/pnas.93.24.13481.

73. Miller, S.G., and Kennedy, M.B. (1986). Regulation of brain type II Ca2+/calmodulin-dependent protein kinase by autophosphorylation: a Ca2+-triggered molecular switch. Cell 44, 861–870. https://doi.org/10.1016/0092-8674(86)90008-5.

74. Morgan, J.I., and Curran, T. (1986). Role of ion flux in the control of c-fos expression. Nature 322, 552–555.

75. Mosbacher, J., Schoepfer, R., Monyer, H., Burnashev, N., Seeburg, P.H., and Ruppersberg, J.P. (1994). A molecular determinant for submillisecond desensitization in glutamate receptors. Science 266, 1059–1062. https://doi.org/10.1126/science.7973663.

76. Mullins, C., Fishell, G., and Tsien, R.W. (2016). Unifying Views of Autism Spectrum Disorders: A Consideration of Autoregulatory Feedback Loops. Neuron 89, 1131–1156. https://doi.org/10.1016/j.neuron.2016.02.017.

77. Murphy, J.G., Sanderson, J.L., Gorski, J.A., Scott, J.D., Catterall, W.A., Sather, W.A., and Dell’Acqua, M.L. (2014). AKAP-anchored PKA maintains neuronal L-type calcium channel activity and NFAT transcriptional signaling. Cell Rep. 7, 1577–1588. https://doi.org/10.1016/j.celrep.2014.04.027.

78. Murthy, V.N., Schikorski, T., Stevens, C.F., and Zhu, Y. (2001). Inactivity produces increases in neurotransmitter release and synapse size. Neuron 32, 673–682. https://doi.org/10.1016/s0896-6273(01)00500-1.

79. Obermair, G.J., Szabo, Z., Bourinet, E., and Flucher, B.E. (2004). Differential targeting of the L-type Ca2+ channel alpha 1C (Ca_v_1.2) to synaptic and extrasynaptic compartments in hippocampal neurons. Eur. J. Neurosci. 19, 2109–2122. https://doi.org/10.1111/j.0953-816X.2004.03272.x.

80. O’Brien, R.J., Kamboj, S., Ehlers, M.D., Rosen, K.R., Fischbach, G.D., and Huganir, R.L. (1998). Activity-dependent modulation of synaptic AMPA receptor accumulation. Neuron 21, 1067–1078. https://doi.org/10.1016/s0896-6273(00)80624-8.

81. O’Leary, T., and Wyllie, D.J.A. (2011). Neuronal homeostasis: time for a change? J. Physiol. 589, 4811–4826. https://doi.org/10.1113/jphysiol.2011.210179.

82. O’Leary, T., van Rossum, M.C.W., and Wyllie, D.J.A. (2010). Homeostasis of intrinsic excitability in hippocampal neurones: dynamics and mechanism of the response to chronic depolarization. J. Physiol. 588, 157–170. https://doi.org/10.1113/jphysiol.2009.181024.

83. Oliet, S.H., Malenka, R.C., and Nicoll, R.A. (1997). Two distinct forms of long-term depression coexist in CA1 hippocampal pyramidal cells. Neuron 18, 969–982. https://doi.org/10.1016/s0896-6273(00)80336-0.

84. Oliveria, S.F., Dell’Acqua, M.L., and Sather, W.A. (2007). AKAP79/150 anchoring of calcineurin controls neuronal L-type Ca2+ channel activity and nuclear signaling. Neuron 55, 261–275. https://doi.org/10.1016/j.neuron.2007.06.032.

85. Park, P., Kang, H., Sanderson, T.M., Bortolotto, Z.A., Georgiou, J., Zhuo, M., Kaang, B.-K., and Collingridge, G.L. (2018). The Role of Calcium-Permeable AMPARs in Long-Term Potentiation at Principal Neurons in the Rodent Hippocampus. Front. Synaptic Neurosci. 10, 42. https://doi.org/10.3389/fnsyn.2018.00042.

86. Poo, M.-M., Pignatelli, M., Ryan, T.J., Tonegawa, S., Bonhoeffer, T., Martin, K.C., Rudenko, A., Tsai, L.-H., Tsien, R.W., Fishell, G., et al. (2016). What is memory? The present state of the engram. BMC Biol. 14, 40. https://doi.org/10.1186/s12915-016-0261-6.

87. Purkey, A.M., and Dell’Acqua, M.L. (2020). Phosphorylation-Dependent Regulation of Ca2+-Permeable AMPA Receptors During Hippocampal Synaptic Plasticity. Front. Synaptic Neurosci. 12, 8. https://doi.org/10.3389/fnsyn.2020.00008.

88. Rameau, G.A., Chiu, L.-Y., and Ziff, E.B. (2004). Bidirectional regulation of neuronal nitric-oxide synthase phosphorylation at serine 847 by the N-methyl-D-aspartate receptor. J. Biol. Chem. 279, 14307–14314. https://doi.org/10.1074/jbc.M311103200.

89. Sanderson, J.L., Gorski, J.A., Gibson, E.S., Lam, P., Freund, R.K., Chick, W.S., and Dell’Acqua, M.L. (2012). AKAP150-Anchored Calcineurin Regulates Synaptic Plasticity by Limiting Synaptic Incorporation of Ca2+- Permeable AMPA Receptors. J. Neurosci. 32, 15036–15052. https://doi.org/10.1523/JNEUROSCI.3326-12.2012.

90. Sanderson, J.L., Gorski, J.A., and Dell’Acqua, M.L. (2016). NMDA Receptor-Dependent LTD Requires Transient Synaptic Incorporation of Ca2+-Permeable AMPARs Mediated by AKAP150-Anchored PKA and Calcineurin. Neuron 89, 1000–1015. https://doi.org/10.1016/j.neuron.2016.01.043.

91. Sanderson, J.L., Scott, J.D., and Dell’Acqua, M.L. (2018). Control of Homeostatic Synaptic Plasticity by AKAP-Anchored Kinase and Phosphatase Regulation of Ca2+-Permeable AMPA Receptors. J. Neurosci. 38, 2863– 2876. https://doi.org/10.1523/JNEUROSCI.2362-17.2018.

92. Schaukowitch, K., Reese, A.L., Kim, S.-K., Kilaru, G., Joo, J.-Y., Kavalali, E.T., and Kim, T.-K. (2017). An Intrinsic Transcriptional Program Underlying Synaptic Scaling during Activity Suppression. Cell Rep. 18, 1512–1526. https://doi.org/10.1016/j.celrep.2017.01.033.

93. Siegel, M., Marder, E., and Abbott, L.F. (1994). Activity-dependent current distributions in model neurons. Proc. Natl. Acad. Sci. U. S. A. 91, 11308–11312. https://doi.org/10.1073/pnas.91.24.11308.

94. Slutsky, I., Sadeghpour, S., Li, B., and Liu, G. (2004). Enhancement of synaptic plasticity through chronically reduced Ca2+ flux during uncorrelated activity. Neuron 44, 835–849.

95. Soares, C., Lee, K.F.H., Nassrallah, W., and Béïque, J.-C. (2013). Differential subcellular targeting of glutamate receptor subtypes during homeostatic synaptic plasticity. J. Neurosci. 33, 13547–13559.

96. Splawski, I., Timothy, K.W., Sharpe, L.M., Decher, N., Kumar, P., Bloise, R., Napolitano, C., Schwartz, P.J., Joseph, R.M., Condouris, K., et al. (2004). Ca(V)1.2 calcium channel dysfunction causes a multisystem disorder including arrhythmia and autism. Cell 119, 19–31. https://doi.org/10.1016/j.cell.2004.09.011.

97. Splawski, I., Timothy, K.W., Decher, N., Kumar, P., Sachse, F.B., Beggs, A.H., Sanguinetti, M.C., and Keating, M.T. (2005). Severe arrhythmia disorder caused by cardiac L-type calcium channel mutations. Proc. Natl. Acad. Sci. U. S. A. 102, 8089–8096; discussion 8086-8. https://doi.org/10.1073/pnas.0502506102.

98. Stemmer, P.M., and Klee, C.B. (1994). Dual calcium ion regulation of calcineurin by calmodulin and calcineurin B. Biochemistry 33, 6859–6866. https://doi.org/10.1021/bi00188a015.

99. Sztainberg, Y., and Zoghbi, H.Y. (2016). Lessons learned from studying syndromic autism spectrum disorders. Nat. Neurosci. 19, 1408–1417. https://doi.org/10.1038/nn.4420.

100. Thiagarajan, T.C., Piedras-Renteria, E.S., and Tsien, R.W. (2002). alpha- and betaCaMKII. Inverse regulation by neuronal activity and opposing effects on synaptic strength. Neuron 36, 1103–1114. https://doi.org/10.1016/s0896-6273(02)01049-8.

101. Thiagarajan, T.C., Lindskog, M., and Tsien, R.W. (2005). Adaptation to synaptic inactivity in hippocampal neurons. Neuron 47, 725–737. https://doi.org/10.1016/j.neuron.2005.06.037.

102. Thiagarajan, T.C., Lindskog, M., Malgaroli, A., and Tsien, R.W. (2007). LTP and adaptation to inactivity: Overlapping mechanisms and implications for metaplasticity. Neuropharmacology 52, 156–175. https://doi.org/10.1016/j.neuropharm.2006.07.030.

103. Tokuoka, H., and Goda, Y. (2008). Activity-dependent coordination of presynaptic release probability and postsynaptic GluR2 abundance at single synapses. Proc. Natl. Acad. Sci. U. S. A. 105, 14656–14661.

104. Torrado Pacheco, A., Bottorff, J., Gao, Y., and Turrigiano, G.G. (2021). Sleep Promotes Downward Firing Rate Homeostasis. Neuron 109, 530–544.e6. https://doi.org/10.1016/j.neuron.2020.11.001.

105. Toyoizumi, T., Kaneko, M., Stryker, M.P., and Miller, K.D. (2014). Modeling the dynamic interaction of Hebbian and homeostatic plasticity. Neuron 84, 497–510. https://doi.org/10.1016/j.neuron.2014.09.036.

106. Trojanowski, N.F., and Turrigiano, G.G. (2021). CaMKIV signaling is not essential for the maintenance of intrinsic or synaptic properties in mouse visual cortex. ENeuro ENEURO.0135–21.2021. https://doi.org/10.1523/eneuro.0135-21.2021.

107. Trojanowski, N.F., Bottorff, J., and Turrigiano, G.G. (2021). Activity labeling in vivo using CaMPARI2 reveals intrinsic and synaptic differences between neurons with high and low firing rate set points. Neuron 109, 663–676.e5. https://doi.org/10.1016/j.neuron.2020.11.027.

108. Turrigiano, G. (2012). Homeostatic synaptic plasticity: local and global mechanisms for stabilizing neuronal function. Cold Spring Harb. Perspect. Biol. 4, a005736. https://doi.org/10.1101/cshperspect.a005736.

109. Turrigiano, G.G., and Nelson, S.B. (1998). Thinking globally, acting locally: AMPA receptor turnover and synaptic strength. Neuron 21, 933–935. https://doi.org/10.1016/s0896-6273(00)80607-8.

110. Turrigiano, G.G., and Nelson, S.B. (2004). Homeostatic plasticity in the developing nervous system. Nat. Rev. Neurosci. 5, 97–107. https://doi.org/10.1038/nrn1327.

111. Turrigiano, G.G., Leslie, K.R., Desai, N.S., Rutherford, L.C., and Nelson, S.B. (1998). Activity-dependent scaling of quantal amplitude in neocortical neurons. Nature 391, 892–896. https://doi.org/10.1038/36103.

112. Wang, C.S., Kavalali, E.T., and Monteggia, L.M. (2021). BDNF signaling in context: From synaptic regulation to psychiatric disorders. Cell 0. https://doi.org/10.1016/j.cell.2021.12.003.

113. Wang, H.-L., Zhang, Z., Hintze, M., and Chen, L. (2011). Decrease in calcium concentration triggers neuronal retinoic acid synthesis during homeostatic synaptic plasticity. J. Neurosci. 31, 17764–17771. https://doi.org/10.1523/JNEUROSCI.3964-11.2011.

114. Wang, X., Wang, Q., Engisch, K.L., and Rich, M.M. (2010). Activity-dependent regulation of the binomial parameters p and n at the mouse neuromuscular junction in vivo. J. Neurophysiol. 104, 2352–2358. https://doi.org/10.1152/jn.00460.2010.

115. Watanabe, Y., Song, T., Sugimoto, K., Horii, M., Araki, N., Tokumitsu, H., Tezuka, T., Yamamoto, T., and Tokuda, M. (2003). Post-synaptic density-95 promotes calcium/calmodulin-dependent protein kinase II-mediated Ser847 phosphorylation of neuronal nitric oxide synthase. Biochem. J 372, 465–471. https://doi.org/10.1042/bj20030380.

116. Weisskopf, M.G., Bauer, E.P., and LeDoux, J.E. (1999). L-type voltage-gated calcium channels mediate NMDA-independent associative long-term potentiation at thalamic input synapses to the amygdala. J. Neurosci. 19, 10512–10519. https://doi.org/10.1523/jneurosci.19-23-10512.1999.

117. Wheeler, D.G., Barrett, C.F., Groth, R.D., Safa, P., and Tsien, R.W. (2008). CaMKII locally encodes L-type channel activity to signal to nuclear CREB in excitation–transcription coupling. J. Cell Biol. 183, 849–863. https://doi.org/10.1083/jcb.200805048.

118. Wierenga, C.J., Ibata, K., and Turrigiano, G.G. (2005). Postsynaptic Expression of Homeostatic Plasticity at Neocortical Synapses. J. Neurosci. 25, 2895–2905. https://doi.org/10.1523/jneurosci.5217-04.2005.

119. Wu, C.-H., Ramos, R., Katz, D.B., and Turrigiano, G.G. (2021). Homeostatic synaptic scaling establishes the specificity of an associative memory. Curr. Biol. 31, 2274–2285.e5. https://doi.org/10.1016/j.cub.2021.03.024.

120. Xu, H., Ginsburg, K.S., Hall, D.D., Zimmermann, M., Stein, I.S., Zhang, M., Tandan, S., Hill, J.A., Horne, M.C., Bers, D., et al. (2010). Targeting of protein phosphatases PP2A and PP2B to the C-terminus of the L-type calcium channel Ca v1.2. Biochemistry 49, 10298–10307. https://doi.org/10.1021/bi101018c.

121. Yeung, L.C., Shouval, H.Z., Blais, B.S., and Cooper, L.N. (2004). Synaptic homeostasis and input selectivity follow from a calcium-dependent plasticity model. Proc. Natl. Acad. Sci. U. S. A. 101, 14943–14948. https://doi.org/10.1073/pnas.0405555101.

122. Zenke, F., and Gerstner, W. (2017). Hebbian plasticity requires compensatory processes on multiple timescales. Philos. Trans. R. Soc. Lond. B Biol. Sci. 372, 20160259. https://doi.org/10.1098/rstb.2016.0259.

123. Zenke, F., Gerstner, W., and Ganguli, S. (2017). The temporal paradox of Hebbian learning and homeostatic plasticity. Curr. Opin. Neurobiol. 43, 166–176. https://doi.org/10.1016/j.conb.2017.03.015.

