## Supplementary material for "Synaptic homeostasis transiently leverages Hebbian mechanisms for a multiphasic response to inactivity": Sun et al., Supplemental Materials

**METHODS**

**Experimental Model and Subject Details**

**Animal Lines**

TS2-neo mice were originally generated by In-Genious Targeting Laboratory, Stony Brook, NY. Following the original nomenclature by Splawski et al., TS2-neo mice express the Gly-to-Arg mutation at position 406 in exon 8 of the *CACNA1C* gene. All cultures were made from dissociated cortices on P0 from TS2-neo heterozygotes crossed to WT, CB57BL/6 WT giving rise to litters with both WT and TS2-neo genotypes. All animals used were genotyped at P0.

**Data & Script Availability**

All data, scripts, and model are available upon reasonable request through corresponding authors.

**Experimental Methods**

**Primary Cortical Culturing**

Cultures were prepared from cortices dissected from P0-P1 mice taken from CB57BL/6 WT and heterozygous TS2-neo crosses. TS2-neo mice are available and as described in (Bader et al., 2011; Bett et al., 2012) and detailed in Animal Lines section. Litters were sex-mixed to minimize potential sex differences and genotyped for the G406R mutation day of dissection and separated between WT and TS2-neo cortices. Cortices were cultured using the methods described previously (Li et al., 2016), with slight modifications. Briefly, WT and TS2 cortices were separately washed twice in ice-cold modified HBSS (4.2 mM $NaHCO_3$ and 1mM HEPES, pH 7.35, 300 mOsm) containing 20% fetal bovine serum (FBS). Samples were washed and digested for 8 min in a papain solution (2.5 mL HBSS + 145 U papain) at 37°C. 8 µL of DNase I (0.2 M), 0.5mM $CaCl_2$ and 1mM MgCl2 was added after for an additional 3 min. Digestion was ceased by adding 5 mL of modified HBSS containing 20% FBS. After addition washing, tissue was triturated using fire polished pasteur pipettes of decreasing diameter with 1 mL of dissociation solution (HBSS + 8 µL of DNase I (0.2 M)). The cell suspension was pelleted twice by centrifugation (10 min at 1000 RPM) and 4°C, first with 500 µL of 4% BSA at the bottom of the tube, with an additional 1 ml dissociation solution trituration with a 1000µL pipette trituration in between. The pellet was then filtered with a 70µm cell strainer, and plated on 12 mm diameter coverslips coated with poly-L-lysine. Cells were counted such that culture density was approximately 90,000 cells per coverslip. Cultures were maintained in NbActiv4 (BrainBits) at 37°C and 5% $CO_2$. Half of the media was changed at 7 DIV and once a week thereafter. Experiments were performed between 13-17 DIV. Media was not changed during treatment.

**Electrophysiological recordings and analysis**

All chemicals were purchased from Sigma-Aldrich unless otherwise noted. Whole-cell voltage-clamp recording of miniature excitatory postsynaptic currents (EPSCs) were conducted from 13-17 days in vitro (DIV). Recordings were performed at 33°C in 4K Tyrode's solution: NaCl 150mM, KCl 4mM, HEPES 10mM, Glucose 10mM, $MgCl_2$ 2mM, $CaCl_2$ 2mM, at 7.40 pH, with 1μM tetrodotoxin (TTX, Alomone Labs) and neurons clamped at -65mV. Internal voltage clamp solutions: 135 $CsMeSO_4$ 135 mM, KCl 5mM, $MgCl_2$ 4mM with ATP Buffer: HEPES 10mM, EGTA 0.3mM, Tris-Phosphocreatine 10mM, Mg-ATP 4 mM, Na-GTP 0.3mM, pH to 7.35 with KOH. Recordings were not corrected for liquid junction potential. Recorded neurons were rejected if they did not meet the following criteria: $V_{rest}$ < -50mV, $R_{access}$ <20 MΩ with <33% change throughout the recording, and > 900μs membrane decay constant. Acute profusion of 10μM philanthotoxin (Cayman Chemical Company) was conducted for up to 10 min after 5 min of baseline recording. Analysis of electrophysiology recordings were conducted with molecular devices ABF files imported into MatLab. Openly available import functions can be found: abfload.m (Forrest Collman, 2009), detectPSP.m (Phil Larimer, 2007). mEPSCs were detected after lowpass filtering and with an event threshold minimum of 5pA. For each event, amplitude was measured by taking the peak of an event, and instantaneous frequency by inverting the interevent interval (IEI) of the previous event. Decay **τ** constants were calculated by fitting each individual event from peak to baseline with a decaying exponential function. Plotted averages were taken from the mean of the mean of each recorded cell. Empirical cumulative distributions were calculated for each condition, normalized to the number of events for each cell. All scripts and data are available at request.

**Imaging**

Cells were fixed in ice-cold 4% paraformaldehyde in phosphate buffer with 20 mM EGTA and 4% sucrose; permeabilized with 0.1% Triton X-100; blocked with 10% normal donkey serum or 10% bovine serum albumin; and incubated overnight at 4°C in primary antibodies. For surface staining of GluA1, coverslips were fixed and blocked in 10% normal donkey serum or 10% bovine serum albumin for 30 minutes in the absence of Triton X-100. Surface staining with primary antibodies was then performed for 1 hour at room temperature. Cells were then permeabilized in 0.1% Triton X-100, and stained with anti-PSD-95 and anti-MAP2 overnight at 4°C. The next day, cells were washed with PBS, incubated at RT for 40 min with Alexa secondary antibodies (1:1000, Molecular Probes), washed again and mounted with ProLong Gold + DAPI (Invitrogen).

Fixed immunostained cells were imaged with a 63X oil objective on a Zeiss LSM 800 confocal microscope at 2048 x 2048 resolution. Z-stacks were taken such the layer of dendrites was within the full Z-range (approximately 3 – 5 μm). At least two biological replicates were taken for each experimental timepoint and genotype. Maximum intensity z-projections where created with the open-source bioimaging program suite ICY with the Zeiss microscopy software package were then created for image analysis. All intensity quantification was performed using ICY. ROIs were drawn using the following criteria: MAP2 staining was present with PSD-95 puncta that were juxtaposed with channel to be measured (ex. GluA1, pCaMKII, etc) blinded. Approximately 30 to 50 ROIs were taken from each maximum intensity projection. A region of interest lacking cells or neurites was selected in each maximum intensity projection as background, and the mean intensity was then subtracted from all intensity measurements from that field of view.

Within each ROI, the signal of interest (surface GluA1, pCaMKII, αCaMKII, βCaMKII) was colocalized to PSD-95 puncta identified using ICY's spot detector tool with the following settings: UDWTWaveletDetector, bright spot over dark background, scale 2, sensitivity 100, ~3 pixels, without filtering. The region without PSD-95 spots within an ROI was designated as dendritic shaft. The mean intensity was measured, giving two mean intensity measurements for each ROI: synaptic (colocalized to PSD-95) and shaft (not colocalized to PSD-95). ROI measurements from each replicate were combined for each timepoint. Mean intensities were then imported into MATLAB and Prism 9 to conduct statistical analyses and data visualization.

**Statistics**

*Statistical Tests for Group Effects*

Statistical analyses and tests were performed using Graphpad Prism (versions 8 and 9) and Matlab. Ordinary one-way ANOVAs were used to compare WT experiment, Figure 1 with two-sided comparisons and corrections. Ordinary two-way ANOVAs were used to compare group means with regard to genotype (WT, TiS) and hours of chronic drug treatment (TTX, FK506, KN93) and interactions. Two-sided comparisons were always conducted with multiple comparison corrections (Tukey correction for comparisons between TTX timepoints within genotype, and Sidak correction for TTX timepoints across genotypes). For PhTx experiments, mEPSCs were aggregated in minute bins, and aligned to time of drug application. Times before drug application were labelled as baseline. For PhTx to baseline comparisons, the first four minutes of recording were binned as "baseline", the last 7 minutes were binned as "+PhTx". F values and degrees of freedom are reported in main text, figures, and supplemental tables. Statistical significances are indicated with *$p<0.05$, **$p<0.01$, etc. and are detailed in each figure legend and in the supplementary tables. Data is represented as mean±SEM. Number of samples, test results, and corresponding $p$ values can be found in figures and main text, with full reports in supplemental figures and tables.

*Linear regression*

Linear regression fits by least-squares were conducted using Matlab's mldivide function or '\' operator to identify the regression coefficient. Datasets were structured such that a given condition and timepoint, that one mean measurement (i.e. mean Deacy Tau or mean synaptic GluA1 in Figure 3) as the independent variable and the other (i.e. mean mEPSC amplitude) as the dependent variable. $R^2$ values were calculated by: $R^2 = 1 - \frac{\sum_{i=1}^{n}(y_i - \hat{y}_i)^2}{\sum_{i=1}^{n}(y_i - \bar{y}_i)^2}$ .

**Model Supplement**

*Model definition*

We modeled the average concentration of postsynaptic calcium, $Ca$, represented on a negative log scale as $pCa$, treating it as the combination of a baseline level of calcium $Ca_0$, and quantal rate-mediated increments in calcium through synaptic GluA1-independent and GluA1-dependent sources,

$$pCa = -\log(Ca_0 + Ca_{PSP0}R + \overline{Ca}_{GluA1}RA) \quad [1]$$

where $R$ is the total rate of quantal delivery in Hz (including both spontaneous PSCs (minis), and evoked EPSCs), $Ca_{PSP0}$ is the calcium from PSCs at 1 Hz, $A$ is the proportion of GluA1 that has been phosphorylated and trafficked to the synapse, and $\overline{Ca}_{GluA1}$is the maximal calcium provided by GluA1 from PSCs at 1 Hz (i.e. when *A*=1). $Ca_{PSP0}$ and $\overline{Ca}_{GluA1}$were chosen to give a plausible dynamic range of calcium levels at physiological quantal rates. This simple summation of contributions from various calcium sources is a first approximation, ignoring possible non-linear dependence of restorative processes on $Ca$. This expression is also repeated as the first equation in main text.

GluA1 activation by a combination of membrane insertion and C-terminal phosphorylation (Diering and Huganir, 2018) was modeled as a kinetic equation

$$\dot{A} = k_f(1-A) - k_dA \quad [2]$$

in which $k_f$ is the rate of phosphorylation and $k_d$ is the rate of dephosphorylation (Equation S1).

$$k_f = k_{f_0} + \bar{k}_{CaMK}m$$
$$k_d = k_{d_0} + \bar{k}_{CaN}n$$

where $k_f$ and $k_d$ were elevated by the calcium-dependent activation of CaMKII and CaN, respectively, over and above a calcium-independent baseline level, where $k_{f_0}$ and $k_{d_0}$ are the baseline levels of phosphorylation and dephosphorylation, respectively. $m$ and $n$ are dynamic variables representing the activation level of CaMKII and CaN, and $\bar{k}_{CaMK}$ and $\bar{k}_{CaN}$ are the corresponding maximal dynamic ranges.

The dynamics of the $m$ and $n$ gates were each modeled by a first order differential equation,

$$\tau_x \frac{dx}{dt} = -x + x_\infty(pCa)$$

$$x_\infty(pCa) = \left(1 + e^{S_x(pCa - pCa_x)}\right)^{-1} \qquad x \in \{m, n, b\}$$

with time constant tau, and sigmoid activation functions that define steady state for each gate as a function of $pCa$.

The CaMKII activation function was further modulated by a variable, $b$, that represents the proportion of the β isoform, such that with 100% βCamKII, the $m$ activation was more sensitive to calcium by a shift in $pCa$-dependence, based on results from Brocke et al., 1999 and Thiagarajan et al., 2002.

$$pCa_m = pCa_\alpha - bCa_{\Delta b}$$

The parameters for αCamKII and CaN activation functions were chosen to match experimental results of Brocke et al., 1999 and Stemmer and Klee, 1994 respectively. The ratio of $k_{f_0}$and $\bar{k}_{CaN}$ determined the steady-state value of Calcium, $Ia$ ($R$) (Figure 6B), and was chosen so that the steady-state curve was relatively flat at around -7, i.e. such that for a wide range of quantal rate, physiological calcium was maintained by the balance of CaN and kinase activity. All parameter values used can be found in **Table S1**.

The effects of Timothy syndrome were modeled by increasing $Ca_{PSP0}$ by a factor of 1.2 and increasing $pCa_m$ by 0.1.

*Steady state response*

The steady state response of the model to presynaptic rate (Figures 6C, S8A, S8B) was solved by rearranging Equation 1 to separate $R$ from terms containing $pCa$, and fixing the value of $A$ under various conditions.

$$\frac{10^{-pCa} - Ca_0}{Ca_{PSP0} + \overline{Ca}_{GluA1}A} = R$$

Namely, we compared the case with $A = 1$ (CPARs maximally active and conductive), $A = 0$ (no CPARs), and $A$ set to a steady state value determined by the calcium dependent phosphokinetic gates

$$A = \frac{k_f}{k_f + k_d} = \frac{k_{f_0} + \bar{k}_{CaMK}m}{k_{f_0} + \bar{k}_{CaMK}m + k_{d_0} + \bar{k}_{CaN}n}$$

with each model component successively added as outlined below.

1. CaN-only: $n = n_\infty(pCa)$ $m = 0$
2. CaN + αCaMKII: $n = n_\infty(pCa)$ $m = m_\infty(pCa)$
3. CaN + α/βCaMKII $n = n_\infty(pCa)$ $m = m_\infty\big(pCa + Ca_{\Delta b}b_\infty(pCa)\big)$

*Presynaptic oscillation*

The presynaptic oscillation was modeled using an exponentially decaying oscillation,

$$R(t) = f(x) = \begin{cases} R_{eq} - R_0 e^{\frac{-t}{\tau}} \cos\left(\frac{2\pi(t+\varphi)}{f}\right), & x \geq 0 \\ R_{spont}, & x < 0 \end{cases}$$

with parameters selected to approximate the experimentally observed time course ($R_{eq} = 25$; $R_0 = 20$; $\tau = 3000$; $f = 2100$; $\varphi = 300$).

| Parameter | Value | Interpretation | Related Citation |
|---|---|---|---|
| $Ca_0$ | $10^{-8}$ M | Quantal-independent, steady-state free calcium level | (Berridge et al., 2000; Grienberger and Konnerth, 2012) |
| $Ca_{PSP0}$ | 0.05 x $10^{-8}$ M $Hz^{-1}$ | CPAR-independent sources of quantal-mediated calcium | |
| $\overline{Ca}_{GluA1}$ | 1 x $10^{-8}$ M $Hz^{-1}$ | Maximal PSP-mediated calcium via CPARs | |
| $k_{f0}$ | 0.5 x $10^{-3}$ $min^{-1}$ | Rate of CaMKII-independent phosphorylation | |
| $k_{d0}$ | 0 $min^{-1}$ | CaN-independent dephosphorylation | |
| $\bar{k}_{CaMKII}$ | 3 $min^{-1}$ | Maximal CaMKII-mediated phosphorylation rate | |
| $\bar{k}_{CaN}$ | 0.1 $min^{-1}$ | Maximal CaN-mediated dephosphorylation rate | |
| $pCa_m$ | 5.45 | Midpoint of the m-gate activation curve | (Brocke et al., 1999) |
| $pCa_n$ | 6.4 | Midpoint of the n-gate activation curve | (Stemmer and Klee, 1994) |
| $pCa_b$ | 7.0 | Midpoint of the b-gate activation curve | |
| $S_m$ | 8 | Steepness of the m-gate activation curve | (Brocke et al., 1999) |
| $S_n$ | 6 | Steepness of the n-gate activation curve | (Stemmer and Klee, 1994) |
| $S_b$ | -15 | Steepness of the b-gate activation curve | |
| $\tau_m$ | 1 min | Time constant of m-gate activation | (Hanson et al., 1994) |

| | | | |
|---|---|---|---|
| $\tau_n$ | 40 min | Time constant of n-gate activation | (Hubbard and Klee, 1987) |
| $\tau_b$ | 300 min | Time constant of b-gate activation | |
| $Ca_{\Delta b}$ | 1.0 | Maximal leftward shift of the m-gate activation curve, in pCa units, by full α- to βCaMKII conversion | (Brocke et al., 1999) |

**Table S1:** Parameter values for simulations presented. Parameters derived from previous reports are indicated with citation.

#### Supplemental Figures

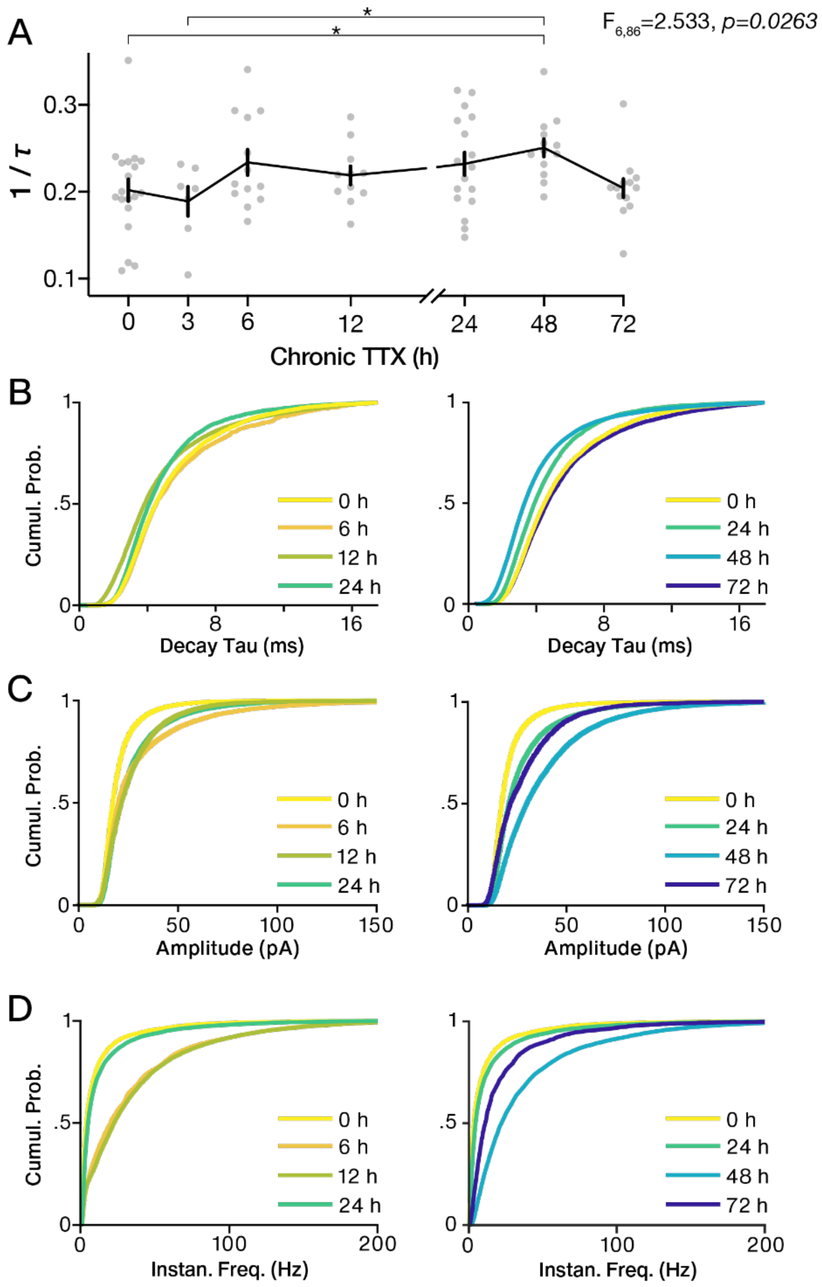

**Figure S1: Extended Data for Figure 1**

A. Mean±SEM 1/τ of mEPSCs from WT neurons recorded.

B. Cumulative distributions of all mEPSCs decay taus by hour of TTX treatment.

C. Cumulative distributions of all mEPSCs amplitudes by hour of TTX treatment.

D. Cumulative distributions of all mEPSCs instantaneous frequencies by hour of TTX treatment.

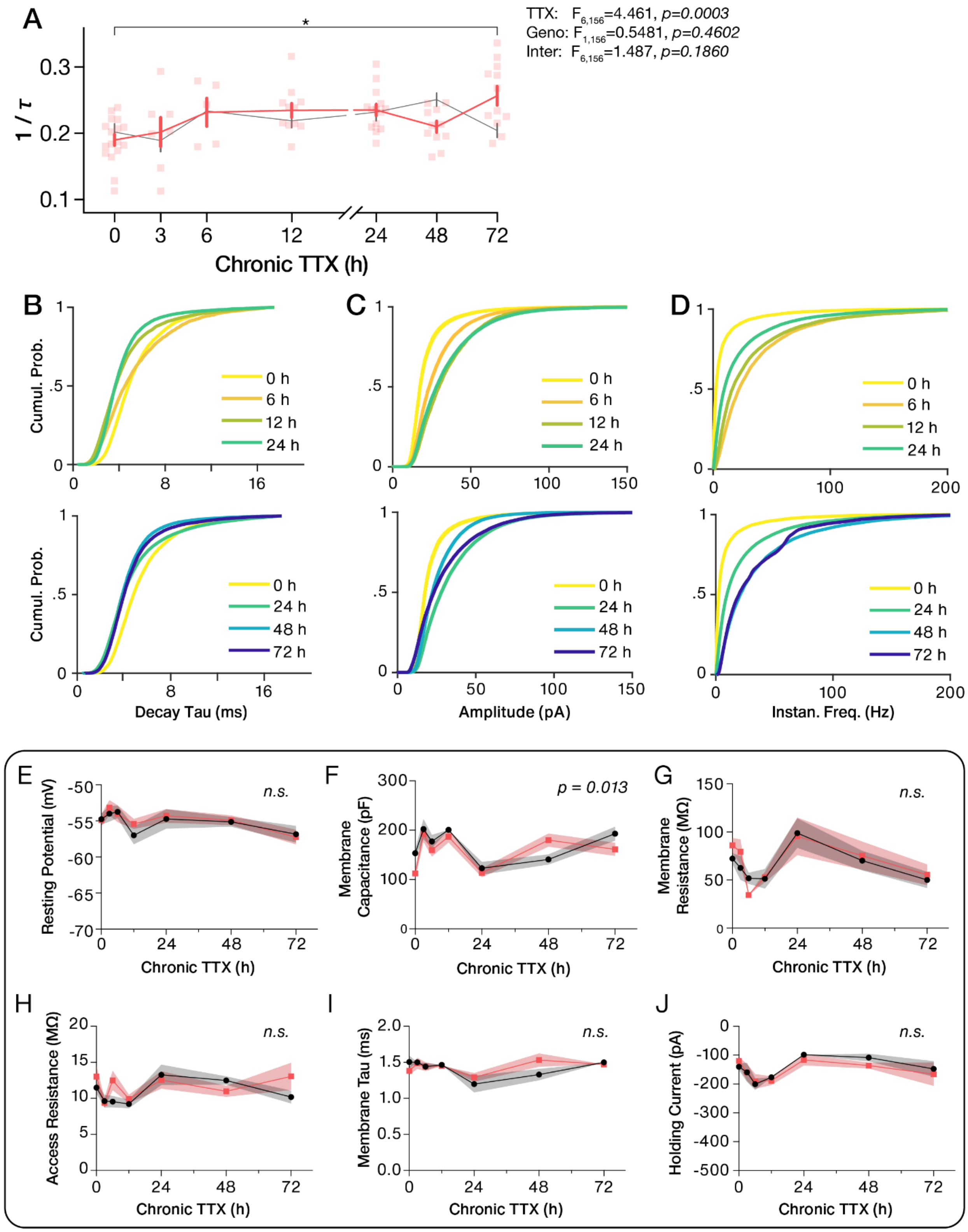

**Figure S2: Extended Data for Figure 2 and 1**

A. Mean±SEM 1/τ of mEPSCs from TiS neurons recorded.

B. Cumulative distributions of all TiS mEPSCs decay taus by hour of TTX treatment.

C. Cumulative distributions of all TiS mEPSCs amplitudes by hour of TTX treatment.

D. Cumulative distributions of all TiS mEPSCs instantaneous frequencies by hour of TTX treatment.

E. Mean±SEM resting membrane potential from recorded neurons from WT (black) and TiS (red) neurons.

F. Mean±SEM membrane capacitance from recorded neurons from WT (black) and TiS (red) neurons.

G. Mean±SEM membrane resistance from recorded neurons from WT (black) and TiS (red) neurons.

H. Mean±SEM access resistance from recorded neurons from WT (black) and TiS (red) neurons.

I. Mean±SEM membrane tau from recorded neurons from WT (black) and TiS (red) neurons.

J. Mean±SEM holding current from recorded neurons from WT (black) and TiS (red) neurons.

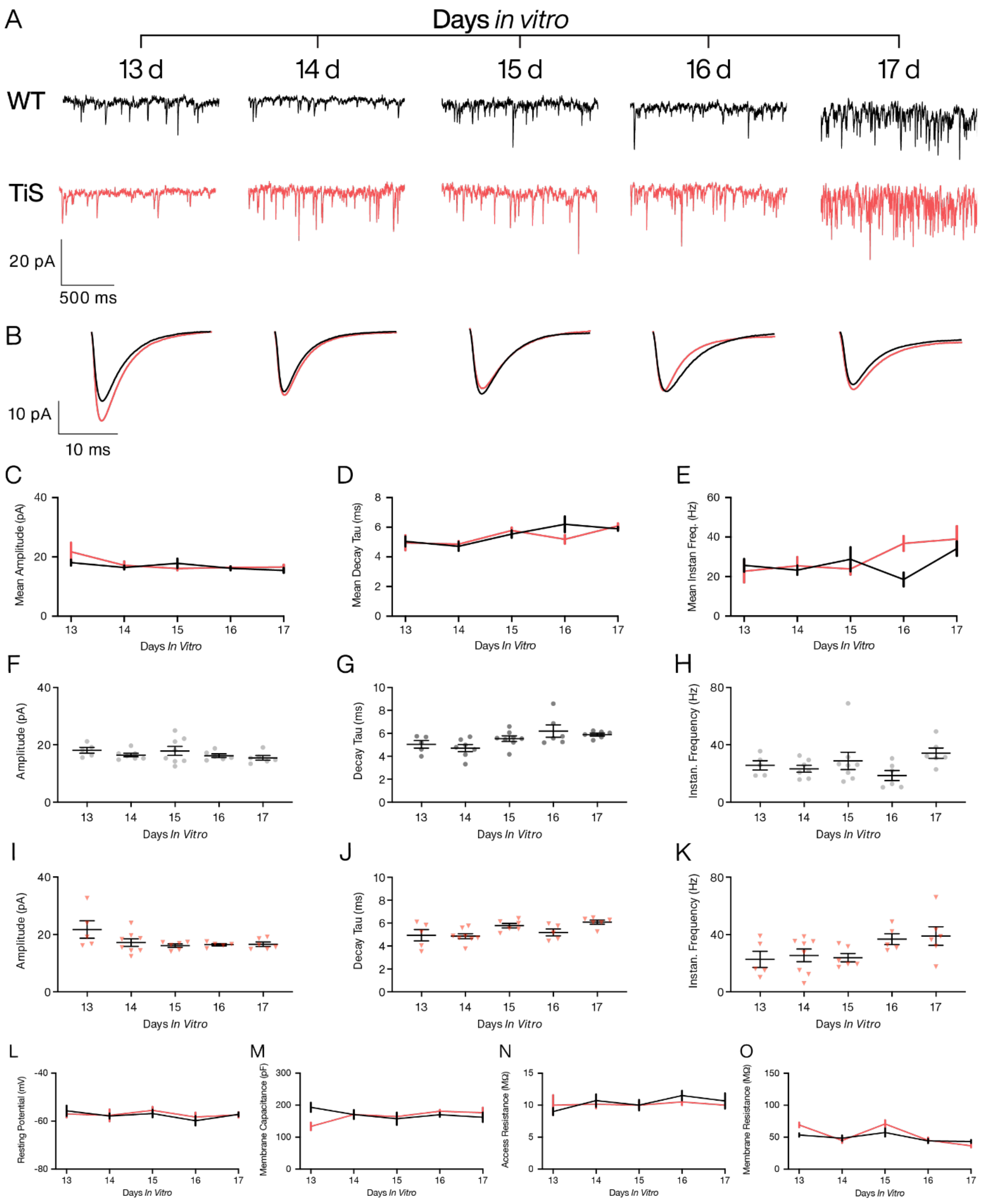

**Figure S3: mEPSC properties from primary cortical cultures do not fluctuate through days *in vitro*.**

A. Example voltage clamp recordings of WT and TiS neurons across days *in vitro* 13, 14, 15, 16, and 17, timepoints all used in TTX experiments.

B. Mean mEPSCs waveforms of the mean waveforms of neurons recorded in the corresponding DIV. Littermatched WT waveforms in black, TiS in red.

C. Mean±SEM amplitude of mEPSCs from WT (black) and TiS (red) across DIV. Two-Way ANOVA: DIV: $F_{4,52} = 2.520$, $p = 0.0521$; Genotype: $F_{1,52} = 0.9586$, $p = 0.3321$; Interaction: $F_{4,52} = 0.085$, $p = 0.3738$.

D. Mean±SEM decay time (plotted as mean $\tau$) of mEPSCs from WT (black) and TiS (red) across DIV. Two-Way ANOVA: DIV: $F_{4,52} = 5.690$, $p = 0.0007$; Genotype: $F_{1,52} = 0.2822$, $p = 0.5975$; Interaction: $F_{4,52} = 1.346$, $p = 0.2654$.

E. Mean±SEM frequency of mEPSCs from WT (black) and TiS (red) across DIV. Two-Way ANOVA: DIV: $F_{4,52} = 2.401$, $p = 0.0616$; Genotype: $F_{1,52} = 1.391$, $p = 0.2436$; Interaction: $F_{4,52} = 1.805$, $p = 0.1420$.

F. Mean±SEM amplitudes from WT recorded cells across DIV.

G. Mean±SEM decay constants from WT recorded cells across DIV.

H. Mean±SEM instantaneous frequencies from WT recorded cells across DIV.

I. Mean±SEM amplitudes from TiS recorded cells across DIV.

J. Mean±SEM decay constants from TiS recorded cells across DIV.

K. Mean±SEM instantaneous frequencies from TiS recorded cells across DIV.

L. Mean±SEM resting membrane potential $V_m$ from WT (black) and TiS (red) recorded cells across DIV.

M. Mean±SEM resting membrane capacitance $C_m$ from WT (black) and TiS (red) recorded cells across DIV.

N. Mean±SEM access resistance from WT (black) and TiS (red) recorded cells across DIV.

O. Mean±SEM membrane resistance from WT (black) and TiS (red) recorded cells across DIV.

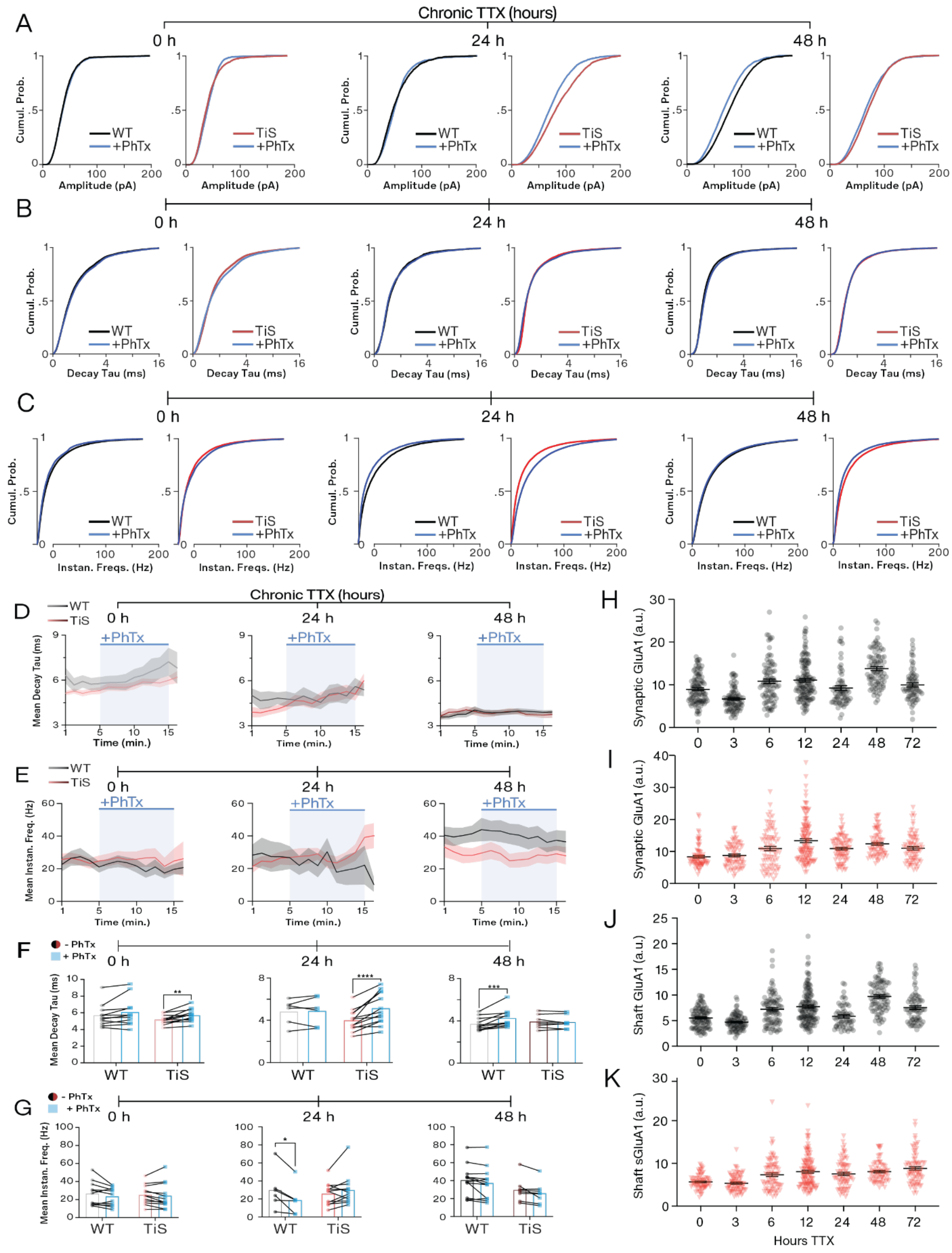

**Figure S4: Extended Data for Figure 3**

A. Cumulative distributions of mEPSC amplitudes from baseline WT (black), baseline TiS (red), and the last 3 minutes of PhTx wash-on (blue) for 0, 24, and 48 h of TTX treatment.

B. Cumulative distributions of mEPSC Decay Tau from baseline WT (black), baseline TiS (red), and the last 3 minutes of PhTx wash-on (blue) for 0, 24, and 48 h of TTX treatment.

C. Cumulative distributions of mEPSC instantaneous frequencies from baseline WT (black), baseline TiS (red), and the last 3 minutes of PhTx wash-on (blue) for 0, 24, and 48 h of TTX treatment.

D. Timecourse of mean±SEM mEPSC decay taus with PhTx wash-on in blue for TTX 0h, 24h, and 48h, for both WT (gray to black) and TiS (pink to maroon).

E. Timecourse of mean±SEM mEPSC instantaneous frequencies with PhTx wash-on in blue for TTX 0h, 24h, and 48h, for both WT (gray to black) and TiS (pink to maroon).

F. Matched decay tau means from 3 minutes of baseline preceding PhTx wash-on and last 3 minutes of PhTx recording.

G. Matched instantaneous frequency means from 3 minutes of baseline preceding PhTx wash-on and last 3 minutes of PhTx recording.

H. Mean±SEM intensities of synaptic sGluA1 staining from WT neurons treated with TTX, individual dendritic ROIs as gray circles.

I. Mean±SEM intensities of synaptic sGluA1 staining from Tis neurons treated with TTX, individual dendritic ROIs as red triangles.

J. Mean±SEM intensities of shaft sGluA1 staining from WT neurons treated with TTX, individual dendritic ROIs as gray circles.

K. Mean±SEM intensities of shaft sGluA1 staining from Tis neurons treated with TTX, individual dendritic ROIs as red triangles.

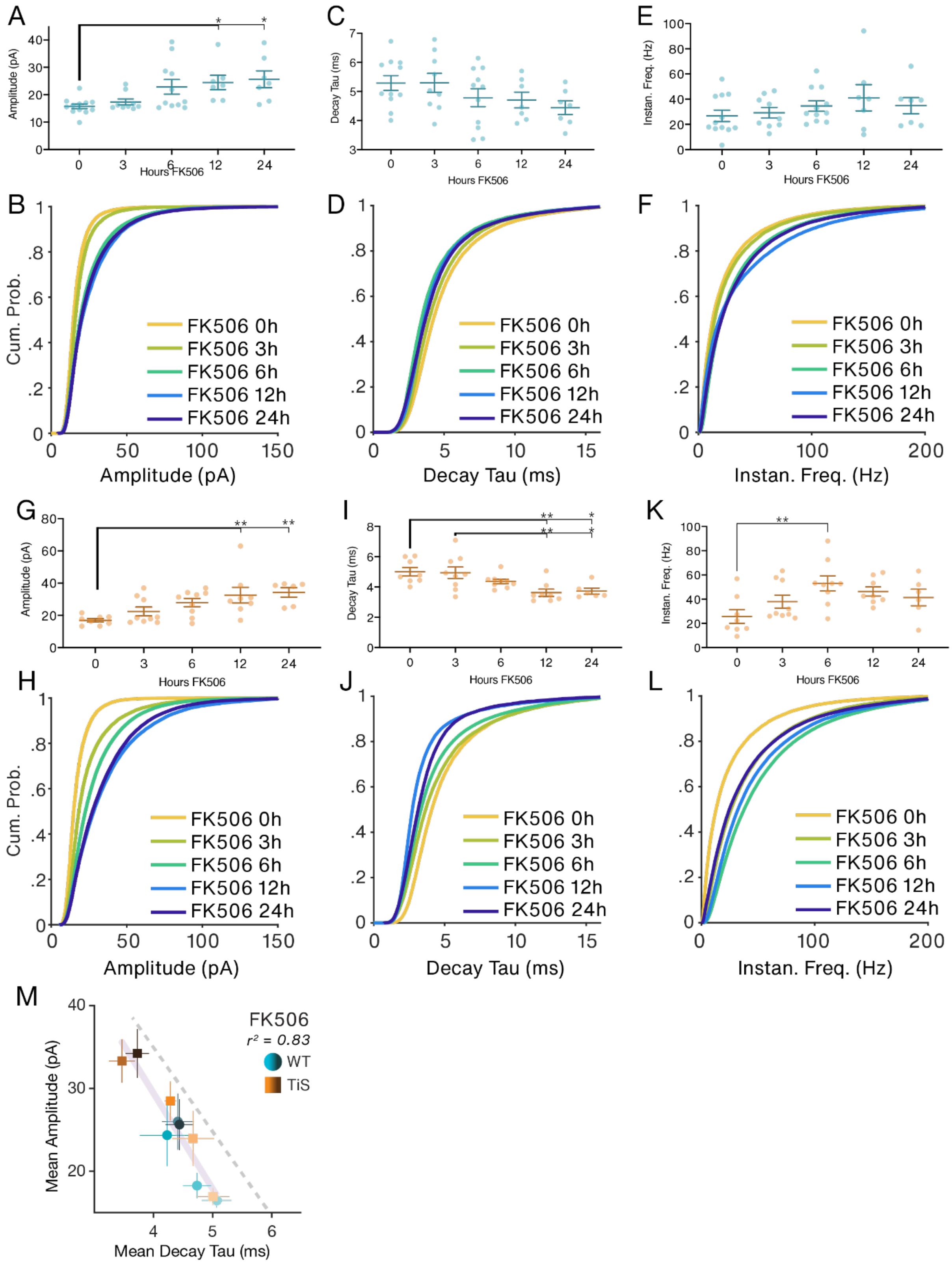

**Figure S5: Extended Data for Figure 4**

A. Mean amplitudes from WT recorded cells across hours FK506. Error bars SEM.

B. Cumulative distributions of all WT mEPSCs amplitudes by hour of TTX treatment.

C. Mean decay taus from WT recorded cells across hours FK506. Error bars SEM.

D. Cumulative distributions of all WT mEPSCs decay taus by hour of TTX treatment.

E. Mean instantaneous frequencies from WT recorded cells across hours FK506. Error bars SEM.

F. Cumulative distributions of all WT mEPSCs instantaneous frequencies by hour of TTX treatment.

G. Mean amplitudes from TiS recorded cells across hours FK506. Error bars SEM.

H. Cumulative distributions of all TiS mEPSCs amplitudes by hour of TTX treatment.

I. Mean decay taus from TiS recorded cells across hours FK506. Error bars SEM.

J. Cumulative distributions of all TiS mEPSCs decay taus by hour of TTX treatment.

K. Mean instantaneous frequencies from TiS recorded cells across hours FK506. Error bars SEM.

L. Cumulative distributions of all TiS mEPSCs instantaneous frequencies by hour of TTX treatment.

M. Mean±SEM amplitude (ordinate) plotted against the mean±SEM decay tau (abscissa) of all timepoints (0 – 24 h FK506, light to dark) and genotypes (WT blue, TiS orange).

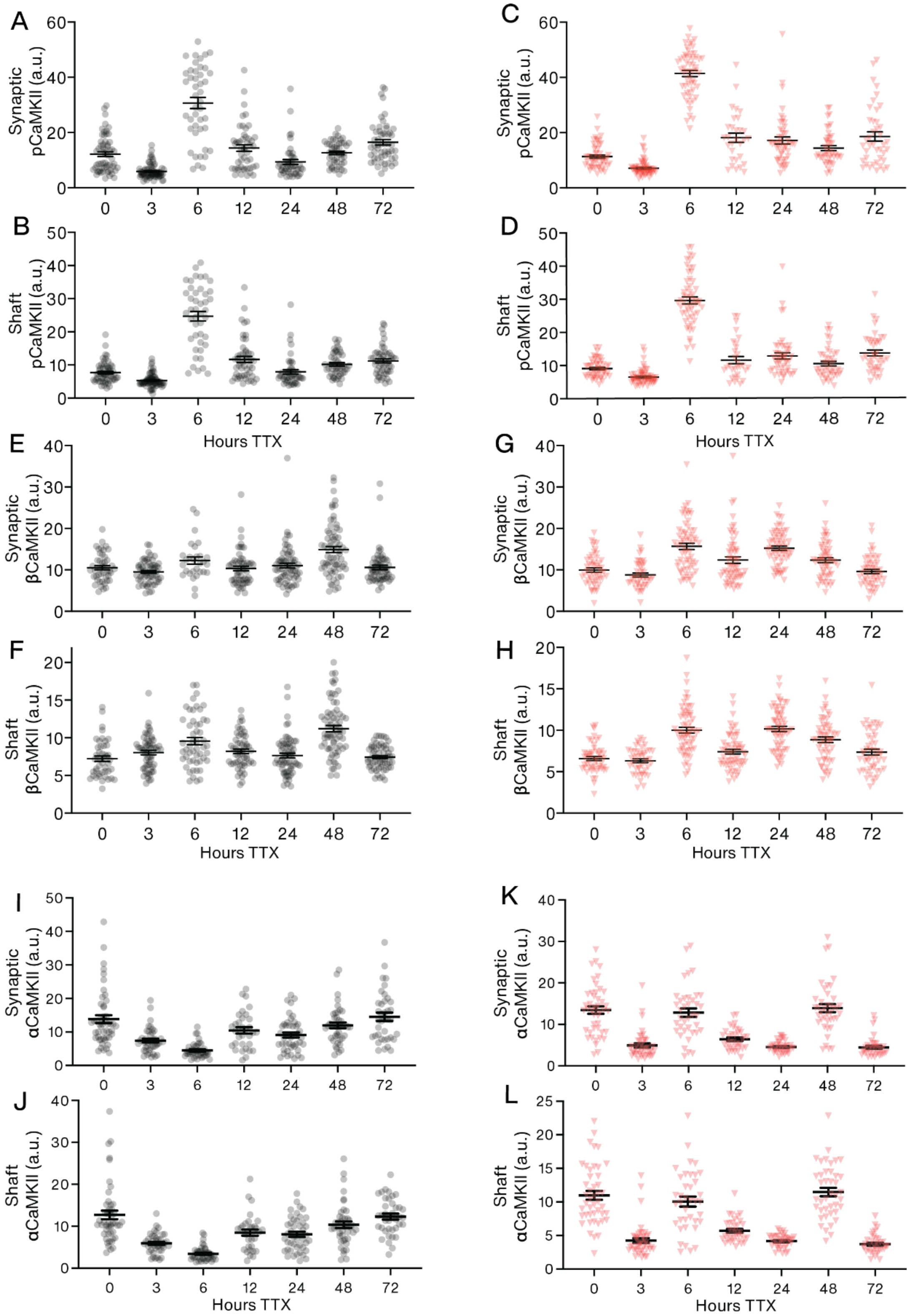

**Figure S6: Extended Data for Figure 5**

A. Mean±SEM intensities of synaptic αCaMKII staining from WT neurons treated with TTX, individual dendritic ROIs as gray circles.

B. Mean±SEM intensities of shaft αCaMKII staining from WT neurons treated with TTX, individual dendritic ROIs as gray circles.

C. Mean±SEM intensities of synaptic αCaMKII staining from TiS neurons treated with TTX, individual dendritic ROIs as red triangles.

D. Mean±SEM intensities of shaft αCaMKII staining from TiS neurons treated with TTX, individual dendritic ROIs as red triangles.

E. Mean±SEM intensities of synaptic βCaMKII staining from WT neurons treated with TTX, individual dendritic ROIs as gray circles.

F. Mean±SEM intensities of shaft βCaMKII staining from WT neurons treated with TTX, individual dendritic ROIs as gray circles.

G. Mean±SEM intensities of synaptic βCaMKII staining from TiS neurons treated with TTX, individual dendritic ROIs as red triangles.

H. Mean±SEM intensities of shaft βCaMKII staining from TiS neurons treated with TTX, individual dendritic ROIs as red triangles.

I. Mean±SEM intensities of synaptic pCaMKII staining from WT neurons treated with TTX, individual dendritic ROIs as gray circles.

J. Mean±SEM intensities of shaft pCaMKII staining from WT neurons treated with TTX, individual dendritic ROIs as gray circles.

K. Mean±SEM intensities of synaptic pCaMKII staining from TiS neurons treated with TTX, individual dendritic ROIs as red triangles.

L. Mean±SEM intensities of shaft pCaMKII staining from TiS neurons treated with TTX, individual dendritic ROIs as red triangles.

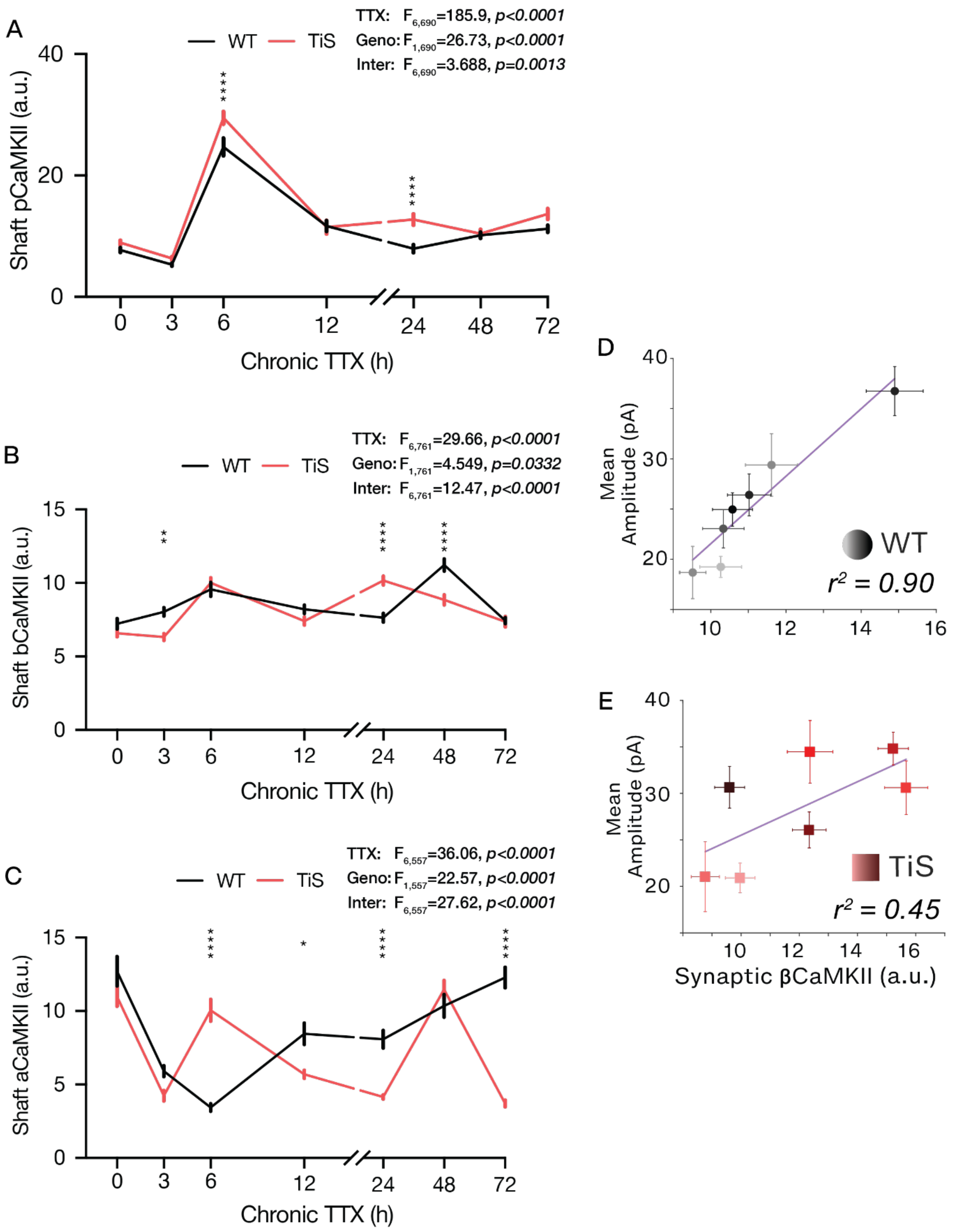

**Figure S7: Extended Data for Figure 5 continued**

A. Mean±SEM intensities of shaft αCaMKII staining from WT (black) and TiS (red) neurons treated with TTX.

B. Mean±SEM intensities of shaft βCaMKII staining from WT (black) and TiS (red) neurons treated with TTX.

C. Mean±SEM intensities of shaft pCaMKII staining from WT (black) and TiS (red) neurons treated with TTX.

D. Mean±SEM amplitude (ordinate) plotted against the mean±SEM synaptic βCaMKII (abscissa) of all timepoints from WT neurons (0 – 72h TTX, light gray to black).

E. Mean±SEM amplitude (ordinate) plotted against the mean±SEM synaptic βCaMKII (abscissa) of all timepoints from TiS neurons (0 – 72h TTX, light red to maroon).

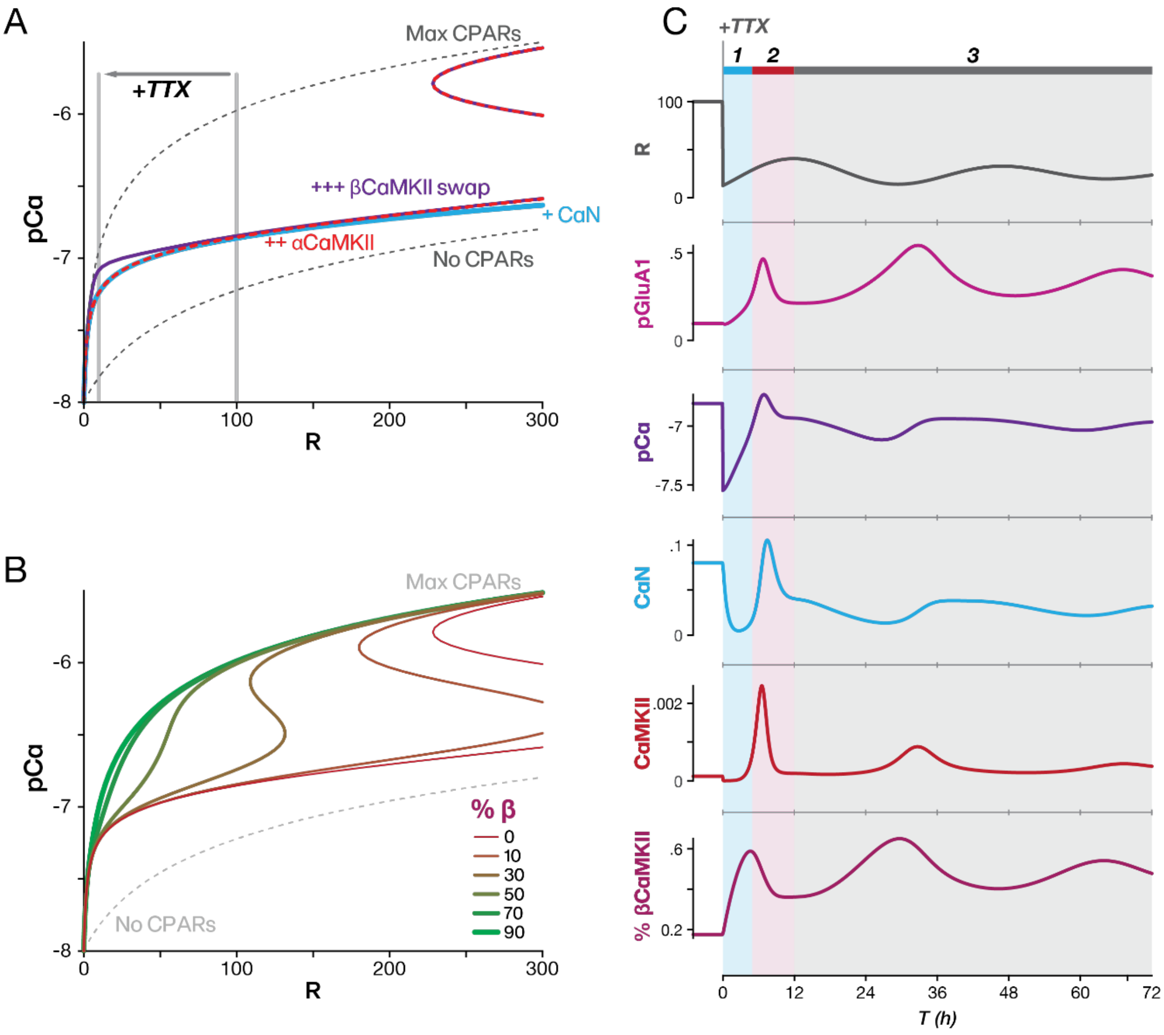

**Figure S8: Extended Data for Figure 6**

A. $pCa(R)$ curve (log scale), the steady-state level of calcium as a function of quantal rate (*R*), for the cases of CaN-homeostasis only ($k_{CaMKII} = 0$, light blue), for CaN-homeostasis with αCaMKII only ($Ca_{\Delta b} = 0$, red dotted), and the full model with switchover from αCaMKII to βCaMKII (purple). The effect of TTX is modeled by dropping the quantal rate from 100 Hz to 10 Hz (transition between gray lines), and then adding the experimentally-observed fluctuations in mini frequency.

B. $pCa(R)$ curve (log scale), the steady-state level of calcium as a function of quantal rate (*R*), for cases of clamped %βCaMKII, with αCaMKII only (thin red) and increments up to 90% βCaMKII (thick green).

C. Response of the full model with an imposed oscillation of presynaptic quantal release, R(t). Stages of the response are indicated by background color with early CaN mediated response (blue), subsequent CaMKII activation (red), and the slow oscillation in pGluA1 (pink).

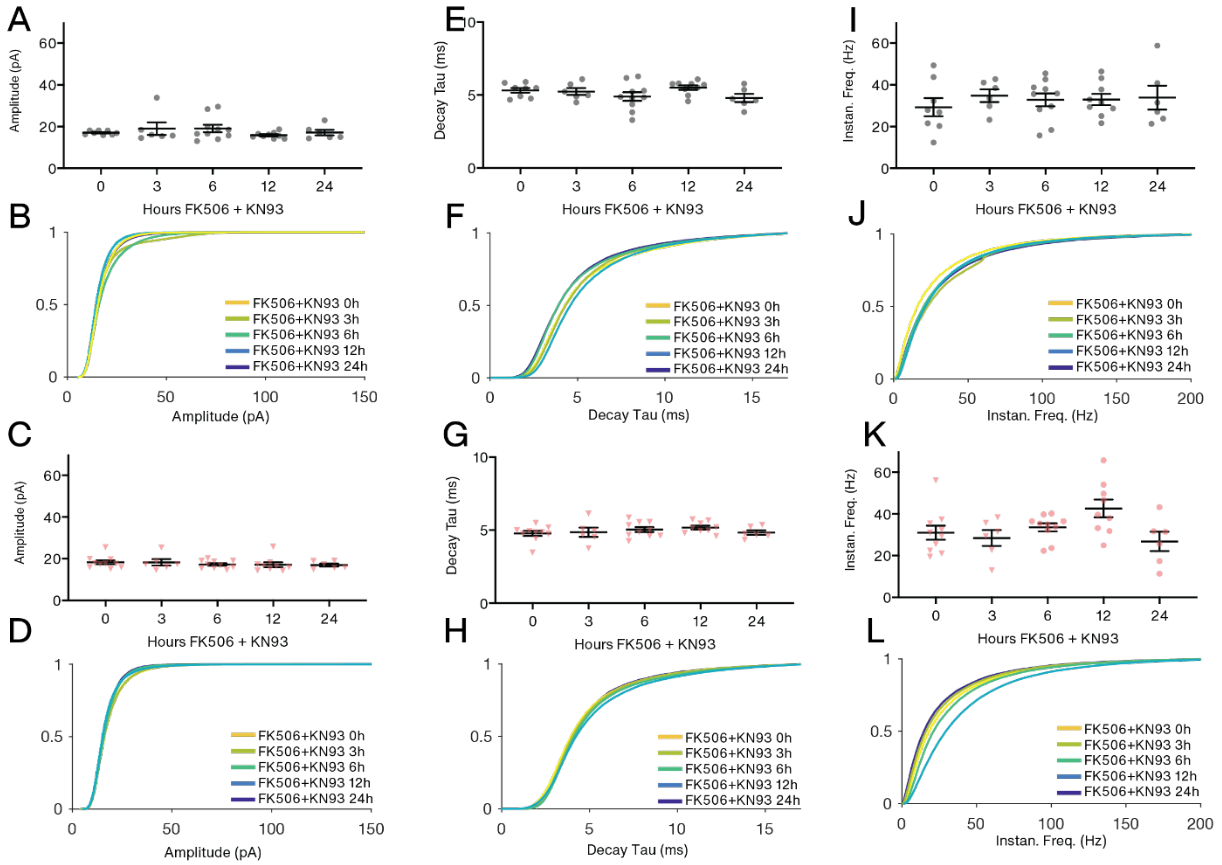

**Figure S9: Extended data for Figure 7**

A. Mean amplitudes from WT recorded cells across hours FK506+KN93.

B. Cumulative distributions of all WT mEPSCs amplitudes by hour of FK506+KN93 treatment.

C. Mean amplitudes from TiS recorded cells across hours FK506+KN93. Error bars SEM.

D. Cumulative distributions of all TiS mEPSCs amplitudes by hour of FK506+KN93 treatment.

E. Mean decay constants from WT recorded cells across hours FK506+KN93. Error bars SEM.

F. Cumulative distributions of all WT mEPSCs decay taus by hour of FK506+KN93 treatment.

G. Mean decay taus from TiS recorded cells across hours FK506+KN93. Error bars SEM.

H. Cumulative distributions of all TiS mEPSCs decay taus by hour of FK506+KN93 treatment.

I. Mean instantaneous frequencies from WT recorded cells across hours FK506+KN93. Error bars SEM.

J. Cumulative distributions of all WT mEPSCs instantaneous frequencies by hour of FK506+KN93treatment.

K. Mean instantaneous frequencies from TiS recorded cells across hours FK506+KN93. Error bars SEM.

L. Cumulative distributions of all TiS mEPSCs instantaneous frequencies by hour of FK506+KN93 treatment.

### Amplitude

| Table Analyzed | WT Amplitude Only | | | | |
|---|---|---|---|---|---|
| Data sets analyzed | A-G | | | | |
| **ANOVA summary** | | | | | |
| F | 8.369 | | | | |
| P value | <0.0001 | | | | |
| P value summary | **** | | | | |
| Significant diff. among means (P < 0.05)? | Yes | | | | |
| R squared | 0.3686 | | | | |
| **Brown-Forsythe test** | | | | | |
| F (DFn, DFd) | 2.742 (6, 86) | | | | |
| P value | 0.0174 | | | | |
| P value summary | * | | | | |
| Are SDs significantly different (P < 0.05)? | Yes | | | | |
| **Bartlett's test** | | | | | |
| Bartlett's statistic (corrected) | 13.94 | | | | |
| P value | 0.0304 | | | | |
| P value summary | * | | | | |
| Are SDs significantly different (P < 0.05)? | Yes | | | | |
| **ANOVA table** | **SS** | **DF** | **MS** | **F (DFn, DFd)** | **P value** |
| Treatment (between columns) | 2992 | 6 | 498.7 | F (6, 86) = 8.369 | P<0.0001 |
| Residual (within columns) | 5125 | 86 | 59.60 | | |
| Total | 8118 | 92 | | | |
| **Data summary** | | | | | |
| Number of treatments (columns) | 7 | | | | |
| Number of values (total) | 93 | | | | |

| Tukey's multiple comparisons test | Mean Diff. | 95.00% CI of diff. | Below threshold? | Summary | Adjusted P Value | |
|---|---|---|---|---|---|---|
| 0 vs. 3 | 0.5597 | -9.746 to 10.87 | No | ns | >0.9999 | A-B |
| 0 vs. 6 | -10.14 | -18.53 to -1.755 | Yes | ** | 0.0079 | A-C |
| 0 vs. 12 | -3.804 | -12.63 to 5.027 | No | ns | 0.8499 | A-D |
| 0 vs. 24 | -7.163 | -14.94 to 0.6180 | No | ns | 0.0917 | A-E |
| 0 vs. 48 | -17.50 | -25.89 to -9.114 | Yes | **** | <0.0001 | A-F |
| 0 vs. 72 | -5.715 | -14.10 to 2.674 | No | ns | 0.3876 | A-G |
| 3 vs. 6 | -10.70 | -21.63 to 0.2230 | No | ns | 0.0588 | B-C |
| 3 vs. 12 | -4.364 | -15.63 to 6.906 | No | ns | 0.9038 | B-D |
| 3 vs. 24 | -7.723 | -18.19 to 2.744 | No | ns | 0.2919 | B-E |
| 3 vs. 48 | -18.06 | -28.99 to -7.136 | Yes | **** | <0.0001 | B-F |
| 3 vs. 72 | -6.275 | -17.20 to 4.652 | No | ns | 0.5956 | B-G |
| 6 vs. 12 | 6.340 | -3.208 to 15.89 | No | ns | 0.4193 | C-D |
| 6 vs. 24 | 2.981 | -5.607 to 11.57 | No | ns | 0.9411 | C-E |
| 6 vs. 48 | -7.359 | -16.50 to 1.783 | No | ns | 0.1989 | C-F |
| 6 vs. 72 | 4.429 | -4.713 to 13.57 | No | ns | 0.7657 | C-G |
| 12 vs. 24 | -3.360 | -12.38 to 5.660 | No | ns | 0.9188 | D-E |
| 12 vs. 48 | -13.70 | -23.25 to -4.151 | Yes | *** | 0.0008 | D-F |
| 12 vs. 72 | -1.911 | -11.46 to 7.637 | No | ns | 0.9965 | D-G |
| 24 vs. 48 | -10.34 | -18.93 to -1.752 | Yes | ** | 0.0083 | E-F |
| 24 vs. 72 | 1.448 | -7.139 to 10.04 | No | ns | 0.9987 | E-G |
| 48 vs. 72 | 11.79 | 2.646 to 20.93 | Yes | ** | 0.0036 | F-G |

| Test details | Mean 1 | Mean 2 | Mean Diff. | SE of diff. | n1 | n2 | q | DF |
|---|---|---|---|---|---|---|---|---|
| 0 vs. 3 | 19.24 | 18.68 | 0.5597 | 3.413 | 19 | 7 | 0.2319 | 86 |
| 0 vs. 6 | 19.24 | 29.39 | -10.14 | 2.779 | 19 | 13 | 5.163 | 86 |
| 0 vs. 12 | 19.24 | 23.05 | -3.804 | 2.925 | 19 | 11 | 1.839 | 86 |
| 0 vs. 24 | 19.24 | 26.41 | -7.163 | 2.577 | 19 | 17 | 3.931 | 86 |
| 0 vs. 48 | 19.24 | 36.75 | -17.50 | 2.779 | 19 | 13 | 8.908 | 86 |
| 0 vs. 72 | 19.24 | 24.96 | -5.715 | 2.779 | 19 | 13 | 2.909 | 86 |
| 3 vs. 6 | 18.68 | 29.39 | -10.70 | 3.619 | 7 | 13 | 4.183 | 86 |
| 3 vs. 12 | 18.68 | 23.05 | -4.364 | 3.732 | 7 | 11 | 1.653 | 86 |
| 3 vs. 24 | 18.68 | 26.41 | -7.723 | 3.467 | 7 | 17 | 3.150 | 86 |
| 3 vs. 48 | 18.68 | 36.75 | -18.06 | 3.619 | 7 | 13 | 7.058 | 86 |
| 3 vs. 72 | 18.68 | 24.96 | -6.275 | 3.619 | 7 | 13 | 2.452 | 86 |
| 6 vs. 12 | 29.39 | 23.05 | 6.340 | 3.163 | 13 | 11 | 2.835 | 86 |
| 6 vs. 24 | 29.39 | 26.41 | 2.981 | 2.844 | 13 | 17 | 1.482 | 86 |
| 6 vs. 48 | 29.39 | 36.75 | -7.359 | 3.028 | 13 | 13 | 3.437 | 86 |
| 6 vs. 72 | 29.39 | 24.96 | 4.429 | 3.028 | 13 | 13 | 2.069 | 86 |
| 12 vs. 24 | 23.05 | 26.41 | -3.360 | 2.987 | 11 | 17 | 1.590 | 86 |
| 12 vs. 48 | 23.05 | 36.75 | -13.70 | 3.163 | 11 | 13 | 6.126 | 86 |
| 12 vs. 72 | 23.05 | 24.96 | -1.911 | 3.163 | 11 | 13 | 0.8546 | 86 |
| 24 vs. 48 | 26.41 | 36.75 | -10.34 | 2.844 | 17 | 13 | 5.141 | 86 |
| 24 vs. 72 | 26.41 | 24.96 | 1.448 | 2.844 | 17 | 13 | 0.7202 | 86 |
| 48 vs. 72 | 36.75 | 24.96 | 11.79 | 3.028 | 13 | 13 | 5.506 | 86 |

### Frequency

| Table Analyzed | WT Frequency | | | | |
|---|---|---|---|---|---|
| Data sets analyzed | A-G | | | | |
| **ANOVA summary** | | | | | |
| F | 15.72 | | | | |
| P value | <0.0001 | | | | |
| P value summary | **** | | | | |
| Significant diff. among means (P < 0.05)? | Yes | | | | |
| R squared | 0.5231 | | | | |
| **Brown-Forsythe test** | | | | | |
| F (DFn, DFd) | 3.560 (6, 86) | | | | |
| P value | 0.0034 | | | | |
| P value summary | ** | | | | |
| Are SDs significantly different (P < 0.05)? | Yes | | | | |
| **Bartlett's test** | | | | | |
| Bartlett's statistic (corrected) | 25.14 | | | | |
| P value | 0.0003 | | | | |
| P value summary | *** | | | | |
| Are SDs significantly different (P < 0.05)? | Yes | | | | |
| **ANOVA table** | **SS** | **DF** | **MS** | **F (DFn, DFd)** | **P value** |
| Treatment (between columns) | 15219 | 6 | 2537 | F (6, 86) = 15.72 | P<0.0001 |
| Residual (within columns) | 13873 | 86 | 161.3 | | |
| Total | 29093 | 92 | | | |
| **Data summary** | | | | | |
| Number of treatments (columns) | 7 | | | | |
| Number of values (total) | 93 | | | | |

| Tukey's multiple comparisons test | Mean Diff | 95.00% CI of diff. | Below threshold? | Summary | Adjusted P Value | |
|---|---|---|---|---|---|---|
| 0 vs. 3 | -2.597 | -19.55 to 14.36 | No | ns | 0.9992 | A-B |
| 0 vs. 6 | -28.22 | -42.02 to -14.42 | Yes | **** | <0.0001 | A-C |
| 0 vs. 12 | -23.12 | -37.65 to -8.592 | Yes | *** | 0.0001 | A-D |
| 0 vs. 24 | -1.112 | -13.91 to 11.69 | No | ns | >0.9999 | A-E |
| 0 vs. 48 | -31.96 | -45.77 to -18.16 | Yes | **** | <0.0001 | A-F |
| 0 vs. 72 | -11.34 | -25.14 to 2.464 | No | ns | 0.1796 | A-G |
| 3 vs. 6 | -25.62 | -43.60 to -7.646 | Yes | *** | 0.0008 | B-C |
| 3 vs. 12 | -20.52 | -39.07 to -1.983 | Yes | * | 0.0203 | B-D |
| 3 vs. 24 | 1.485 | -15.74 to 18.71 | No | ns | >0.9999 | B-E |
| 3 vs. 48 | -29.37 | -47.35 to -11.39 | Yes | **** | <0.0001 | B-F |
| 3 vs. 72 | -8.742 | -26.72 to 9.236 | No | ns | 0.7626 | B-G |
| 6 vs. 12 | 5.099 | -10.61 to 20.81 | No | ns | 0.9570 | C-D |
| 6 vs. 24 | 27.11 | 12.98 to 41.24 | Yes | **** | <0.0001 | C-E |
| 6 vs. 48 | -3.744 | -18.79 to 11.30 | No | ns | 0.9887 | C-F |
| 6 vs. 72 | 16.88 | 1.840 to 31.92 | Yes | * | 0.0177 | C-G |
| 12 vs. 24 | 22.01 | 7.170 to 36.85 | Yes | *** | 0.0004 | D-E |
| 12 vs. 48 | -8.843 | -24.55 to 6.867 | No | ns | 0.6182 | D-F |
| 12 vs. 72 | 11.78 | -3.928 to 27.49 | No | ns | 0.2733 | D-G |
| 24 vs. 48 | -30.85 | -44.98 to -16.72 | Yes | **** | <0.0001 | E-F |
| 24 vs. 72 | -10.23 | -24.36 to 3.902 | No | ns | 0.3142 | E-G |
| 48 vs. 72 | 20.63 | 5.584 to 35.67 | Yes | ** | 0.0015 | F-G |

| Test details | Mean 1 | Mean 2 | Mean Diff. | SE of diff. | n1 | n2 | q | DF |
|---|---|---|---|---|---|---|---|---|
| 0 vs. 3 | 11.87 | 14.46 | -2.597 | 5.616 | 19 | 7 | 0.6539 | 86 |
| 0 vs. 6 | 11.87 | 40.09 | -28.22 | 4.572 | 19 | 13 | 8.730 | 86 |
| 0 vs. 12 | 11.87 | 34.99 | -23.12 | 4.812 | 19 | 11 | 6.795 | 86 |
| 0 vs. 24 | 11.87 | 12.98 | -1.112 | 4.240 | 19 | 17 | 0.3708 | 86 |
| 0 vs. 48 | 11.87 | 43.83 | -31.96 | 4.572 | 19 | 13 | 9.888 | 86 |
| 0 vs. 72 | 11.87 | 23.21 | -11.34 | 4.572 | 19 | 13 | 3.508 | 86 |
| 3 vs. 6 | 14.46 | 40.09 | -25.62 | 5.954 | 7 | 13 | 6.086 | 86 |
| 3 vs. 12 | 14.46 | 34.99 | -20.52 | 6.141 | 7 | 11 | 4.727 | 86 |
| 3 vs. 24 | 14.46 | 12.98 | 1.485 | 5.704 | 7 | 17 | 0.3681 | 86 |
| 3 vs. 48 | 14.46 | 43.83 | -29.37 | 5.954 | 7 | 13 | 6.975 | 86 |
| 3 vs. 72 | 14.46 | 23.21 | -8.742 | 5.954 | 7 | 13 | 2.076 | 86 |
| 6 vs. 12 | 40.09 | 34.99 | 5.099 | 5.203 | 13 | 11 | 1.386 | 86 |
| 6 vs. 24 | 40.09 | 12.98 | 27.11 | 4.680 | 13 | 17 | 8.192 | 86 |
| 6 vs. 48 | 40.09 | 43.83 | -3.744 | 4.982 | 13 | 13 | 1.063 | 86 |
| 6 vs. 72 | 40.09 | 23.21 | 16.88 | 4.982 | 13 | 13 | 4.792 | 86 |
| 12 vs. 24 | 34.99 | 12.98 | 22.01 | 4.915 | 11 | 17 | 6.333 | 86 |
| 12 vs. 48 | 34.99 | 43.83 | -8.843 | 5.203 | 11 | 13 | 2.404 | 86 |
| 12 vs. 72 | 34.99 | 23.21 | 11.78 | 5.203 | 11 | 13 | 3.202 | 86 |
| 24 vs. 48 | 12.98 | 43.83 | -30.85 | 4.680 | 17 | 13 | 9.324 | 86 |
| 24 vs. 72 | 12.98 | 23.21 | -10.23 | 4.680 | 17 | 13 | 3.091 | 86 |
| 48 vs. 72 | 43.83 | 23.21 | 20.63 | 4.982 | 13 | 13 | 5.855 | 86 |

### Decay

| Table Analyzed | WT Decay Only | | | | |
|---|---|---|---|---|---|
| Data sets analyzed | A-G | | | | |
| **ANOVA summary** | | | | | |
| F | 2.533 | | | | |
| P value | 0.0263 | | | | |
| P value summary | * | | | | |
| Significant diff. among means (P < 0.05)? | Yes | | | | |
| R squared | 0.1502 | | | | |
| **Brown-Forsythe test** | | | | | |
| F (DFn, DFd) | 1.244 (6, 86) | | | | |
| P value | 0.2924 | | | | |
| P value summary | ns | | | | |
| Are SDs significantly different (P < 0.05)? | No | | | | |
| **Bartlett's test** | | | | | |
| Bartlett's statistic (corrected) | 22.27 | | | | |
| P value | 0.0011 | | | | |
| P value summary | ** | | | | |
| Are SDs significantly different (P < 0.05)? | Yes | | | | |
| **ANOVA table** | **SS** | **DF** | **MS** | **F (DFn, DFd)** | **P value** |
| Treatment (between columns) | 21.58 | 6 | 3.597 | F (6, 86) = 2.533 | P=0.0263 |
| Residual (within columns) | 122.2 | 86 | 1.420 | | |
| Total | 143.7 | 92 | | | |
| **Data summary** | | | | | |
| Number of treatments (columns) | 7 | | | | |
| Number of values (total) | 93 | | | | |

| Tukey's multiple comparisons test | Mean Diff. | 95.00% CI of diff. | Below threshol | Summary | Adjusted P Value | |
|---|---|---|---|---|---|---|
| 0 vs. 3 | -0.3042 | -1.895 to 1.287 | No | ns | 0.9973 | A-B |
| 0 vs. 6 | 0.8839 | -0.4113 to 2.179 | No | ns | 0.3854 | A-C |
| 0 vs. 12 | 0.6830 | -0.6803 to 2.046 | No | ns | 0.7366 | A-D |
| 0 vs. 24 | 0.8047 | -0.3966 to 2.006 | No | ns | 0.4084 | A-E |
| 0 vs. 48 | 1.292 | -0.002849 to 2.588 | No | ns | 0.0509 | A-F |
| 0 vs. 72 | 0.2991 | -0.9961 to 1.594 | No | ns | 0.9924 | A-G |
| 3 vs. 6 | 1.188 | -0.4989 to 2.875 | No | ns | 0.3470 | B-C |
| 3 vs. 12 | 0.9872 | -0.7527 to 2.727 | No | ns | 0.6093 | B-D |
| 3 vs. 24 | 1.109 | -0.5071 to 2.725 | No | ns | 0.3787 | B-E |
| 3 vs. 48 | 1.597 | -0.09044 to 3.284 | No | ns | 0.0758 | B-F |
| 3 vs. 72 | 0.6032 | -1.084 to 2.290 | No | ns | 0.9324 | B-G |
| 6 vs. 12 | -0.2009 | -1.675 to 1.273 | No | ns | 0.9996 | C-D |
| 6 vs. 24 | -0.07916 | -1.405 to 1.247 | No | ns | >0.9999 | C-E |
| 6 vs. 48 | 0.4085 | -1.003 to 1.820 | No | ns | 0.9755 | C-F |
| 6 vs. 72 | -0.5848 | -1.996 to 0.8266 | No | ns | 0.8718 | C-G |
| 12 vs. 24 | 0.1217 | -1.271 to 1.514 | No | ns | >0.9999 | D-E |
| 12 vs. 48 | 0.6094 | -0.8648 to 2.084 | No | ns | 0.8730 | D-F |
| 12 vs. 72 | -0.3839 | -1.858 to 1.090 | No | ns | 0.9857 | D-G |
| 24 vs. 48 | 0.4877 | -0.8381 to 1.813 | No | ns | 0.9233 | E-F |
| 24 vs. 72 | -0.5056 | -1.831 to 0.8202 | No | ns | 0.9099 | E-G |
| 48 vs. 72 | -0.9933 | -2.405 to 0.4181 | No | ns | 0.3478 | F-G |

| Test details | Mean 1 | Mean 2 | Mean Diff. | SE of diff. | n1 | n2 | q | DF |
|---|---|---|---|---|---|---|---|---|
| 0 vs. 3 | 5.353 | 5.657 | -0.3042 | 0.5270 | 19 | 7 | 0.8163 | 86 |
| 0 vs. 6 | 5.353 | 4.469 | 0.8839 | 0.4290 | 19 | 13 | 2.914 | 86 |
| 0 vs. 12 | 5.353 | 4.670 | 0.6830 | 0.4515 | 19 | 11 | 2.139 | 86 |
| 0 vs. 24 | 5.353 | 4.548 | 0.8047 | 0.3979 | 19 | 17 | 2.860 | 86 |
| 0 vs. 48 | 5.353 | 4.060 | 1.292 | 0.4290 | 19 | 13 | 4.260 | 86 |
| 0 vs. 72 | 5.353 | 5.054 | 0.2991 | 0.4290 | 19 | 13 | 0.9859 | 86 |
| 3 vs. 6 | 5.657 | 4.469 | 1.188 | 0.5587 | 7 | 13 | 3.007 | 86 |
| 3 vs. 12 | 5.657 | 4.670 | 0.9872 | 0.5762 | 7 | 11 | 2.423 | 86 |
| 3 vs. 24 | 5.657 | 4.548 | 1.109 | 0.5352 | 7 | 17 | 2.930 | 86 |
| 3 vs. 48 | 5.657 | 4.060 | 1.597 | 0.5587 | 7 | 13 | 4.041 | 86 |
| 3 vs. 72 | 5.657 | 5.054 | 0.6032 | 0.5587 | 7 | 13 | 1.527 | 86 |
| 6 vs. 12 | 4.469 | 4.670 | -0.2009 | 0.4883 | 13 | 11 | 0.5819 | 86 |
| 6 vs. 24 | 4.469 | 4.548 | -0.07916 | 0.4391 | 13 | 17 | 0.2550 | 86 |
| 6 vs. 48 | 4.469 | 4.060 | 0.4085 | 0.4675 | 13 | 13 | 1.236 | 86 |
| 6 vs. 72 | 4.469 | 5.054 | -0.5848 | 0.4675 | 13 | 13 | 1.769 | 86 |
| 12 vs. 24 | 4.670 | 4.548 | 0.1217 | 0.4612 | 11 | 17 | 0.3733 | 86 |
| 12 vs. 48 | 4.670 | 4.060 | 0.6094 | 0.4883 | 11 | 13 | 1.765 | 86 |
| 12 vs. 72 | 4.670 | 5.054 | -0.3839 | 0.4883 | 11 | 13 | 1.112 | 86 |
| 24 vs. 48 | 4.548 | 4.060 | 0.4877 | 0.4391 | 17 | 13 | 1.571 | 86 |
| 24 vs. 72 | 4.548 | 5.054 | -0.5056 | 0.4391 | 17 | 13 | 1.628 | 86 |
| 48 vs. 72 | 4.060 | 5.054 | -0.9933 | 0.4675 | 13 | 13 | 3.005 | 86 |

**Table S2: Statistics for Figure 1**

#### Amplitude

| | | | | | |
|---|---|---|---|---|---|
| Table Analyzed | Mean Amplitude | | | | |
| **Two-way ANOVA** | Ordinary | | | | |
| Alpha | 0.05 | | | | |
| **Source of Variation** | **% of total variation** | **P value** | **P value summary** | **Significant?** | |
| Interaction | 11.86 | 0.0001 | *** | Yes | |
| Hours TTX | 23.06 | <0.0001 | **** | Yes | |
| Genotype | 2.022 | 0.0257 | * | Yes | |
| **ANOVA table** | **SS (Type III)** | **DF** | **MS** | **F (DFn, DFd)** | **P value** |
| Interaction | 1803 | 6 | 300.5 | F (6, 156) = 4.959 | P=0.0001 |
| Hours TTX | 3505 | 6 | 584.1 | F (6, 156) = 9.640 | P<0.0001 |
| Genotype | 307.3 | 1 | 307.3 | F (1, 156) = 5.071 | P=0.0257 |
| Residual | 9453 | 156 | 60.60 | | |
| **Difference between column means** | | | | | |
| Predicted (LS) mean of wt | 25.50 | | | | |
| Predicted (LS) mean of ts | 28.37 | | | | |
| Difference between predicted means | -2.871 | | | | |
| SE of difference | 1.275 | | | | |
| 95% CI of difference | -5.389 to -0.3526 | | | | |
| **Data summary** | | | | | |
| Number of columns (Genotype) | 2 | | | | |
| Number of rows (Hours TTX) | 7 | | | | |
| Number of values | 170 | | | | |

| Test details | Predicted (LS) mean 1 | Predicted (LS) mea | Predicted (LS) mea | SE of diff. | N1 | N2 | q | DF |
|---|---|---|---|---|---|---|---|---|
| wt | | | | | | | | |
| Row 1 vs. Row 2 | 19.24 | 18.68 | 0.5597 | 3.442 | 19 | 7 | 0.2300 | 156.0 |
| Row 1 vs. Row 3 | 19.24 | 29.39 | -10.14 | 2.802 | 19 | 13 | 5.120 | 156.0 |
| Row 1 vs. Row 4 | 19.24 | 23.05 | -3.804 | 2.949 | 19 | 11 | 1.824 | 156.0 |
| Row 1 vs. Row 5 | 19.24 | 26.41 | -7.163 | 2.599 | 19 | 17 | 3.898 | 156.0 |
| Row 1 vs. Row 6 | 19.24 | 36.75 | -17.50 | 2.802 | 19 | 13 | 8.835 | 156.0 |
| Row 1 vs. Row 7 | 19.24 | 24.96 | -5.715 | 2.802 | 19 | 13 | 2.885 | 156.0 |
| Row 2 vs. Row 3 | 18.68 | 29.39 | -10.70 | 3.649 | 7 | 13 | 4.148 | 156.0 |
| Row 2 vs. Row 4 | 18.68 | 23.05 | -4.364 | 3.764 | 7 | 11 | 1.640 | 156.0 |
| Row 2 vs. Row 5 | 18.68 | 26.41 | -7.723 | 3.496 | 7 | 17 | 3.124 | 156.0 |
| Row 2 vs. Row 6 | 18.68 | 36.75 | -18.06 | 3.649 | 7 | 13 | 7.000 | 156.0 |
| Row 2 vs. Row 7 | 18.68 | 24.96 | -6.275 | 3.649 | 7 | 13 | 2.432 | 156.0 |
| Row 3 vs. Row 4 | 29.39 | 23.05 | 6.340 | 3.189 | 13 | 11 | 2.812 | 156.0 |
| Row 3 vs. Row 5 | 29.39 | 26.41 | 2.981 | 2.868 | 13 | 17 | 1.470 | 156.0 |
| Row 3 vs. Row 6 | 29.39 | 36.75 | -7.359 | 3.053 | 13 | 13 | 3.409 | 156.0 |
| Row 3 vs. Row 7 | 29.39 | 24.96 | 4.429 | 3.053 | 13 | 13 | 2.052 | 156.0 |
| Row 4 vs. Row 5 | 23.05 | 26.41 | -3.360 | 3.012 | 11 | 17 | 1.577 | 156.0 |
| Row 4 vs. Row 6 | 23.05 | 36.75 | -13.70 | 3.189 | 11 | 13 | 6.075 | 156.0 |
| Row 4 vs. Row 7 | 23.05 | 24.96 | -1.911 | 3.189 | 11 | 13 | 0.8475 | 156.0 |
| Row 5 vs. Row 6 | 26.41 | 36.75 | -10.34 | 2.868 | 17 | 13 | 5.099 | 156.0 |
| Row 5 vs. Row 7 | 26.41 | 24.96 | 1.448 | 2.868 | 17 | 13 | 0.7142 | 156.0 |
| Row 6 vs. Row 7 | 36.75 | 24.96 | 11.79 | 3.053 | 13 | 13 | 5.460 | 156.0 |
| ts | | | | | | | | |
| Row 1 vs. Row 2 | 20.92 | 21.04 | -0.1219 | 3.496 | 17 | 7 | 0.04932 | 156.0 |
| Row 1 vs. Row 3 | 20.92 | 30.61 | -9.698 | 3.960 | 17 | 5 | 3.463 | 156.0 |
| Row 1 vs. Row 4 | 20.92 | 34.47 | -13.56 | 3.012 | 17 | 11 | 6.365 | 156.0 |
| Row 1 vs. Row 5 | 20.92 | 34.82 | -13.90 | 2.809 | 17 | 14 | 6.997 | 156.0 |
| Row 1 vs. Row 6 | 20.92 | 26.07 | -5.155 | 3.012 | 17 | 11 | 2.420 | 156.0 |
| Row 1 vs. Row 7 | 20.92 | 30.65 | -9.732 | 2.935 | 17 | 12 | 4.689 | 156.0 |
| Row 2 vs. Row 3 | 21.04 | 30.61 | -9.576 | 4.558 | 7 | 5 | 2.971 | 156.0 |
| Row 2 vs. Row 4 | 21.04 | 34.47 | -13.43 | 3.764 | 7 | 11 | 5.048 | 156.0 |
| Row 2 vs. Row 5 | 21.04 | 34.82 | -13.78 | 3.603 | 7 | 14 | 5.407 | 156.0 |
| Row 2 vs. Row 6 | 21.04 | 26.07 | -5.033 | 3.764 | 7 | 11 | 1.891 | 156.0 |
| Row 2 vs. Row 7 | 21.04 | 30.65 | -9.610 | 3.702 | 7 | 12 | 3.671 | 156.0 |
| Row 3 vs. Row 4 | 30.61 | 34.47 | -3.858 | 4.199 | 5 | 11 | 1.299 | 156.0 |
| Row 3 vs. Row 5 | 30.61 | 34.82 | -4.202 | 4.056 | 5 | 14 | 1.465 | 156.0 |
| Row 3 vs. Row 6 | 30.61 | 26.07 | 4.543 | 4.199 | 5 | 11 | 1.530 | 156.0 |
| Row 3 vs. Row 7 | 30.61 | 30.65 | -0.03386 | 4.144 | 5 | 12 | 0.01156 | 156.0 |
| Row 4 vs. Row 5 | 34.47 | 34.82 | -0.3441 | 3.136 | 11 | 14 | 0.1551 | 156.0 |
| Row 4 vs. Row 6 | 34.47 | 26.07 | 8.401 | 3.319 | 11 | 11 | 3.580 | 156.0 |
| Row 4 vs. Row 7 | 34.47 | 30.65 | 3.824 | 3.249 | 11 | 12 | 1.664 | 156.0 |
| Row 5 vs. Row 6 | 34.82 | 26.07 | 8.745 | 3.136 | 14 | 11 | 3.943 | 156.0 |
| Row 5 vs. Row 7 | 34.82 | 30.65 | 4.168 | 3.062 | 14 | 12 | 1.925 | 156.0 |
| Row 6 vs. Row 7 | 26.07 | 30.65 | -4.577 | 3.249 | 11 | 12 | 1.992 | 156.0 |

| Tukey's multiple com | Predicted (LS) mean diff. | 95.00% CI of diff. | Below threshold? | Summary | Adjusted P Value |
|---|---|---|---|---|---|
| wt | | | | | |
| Row 1 vs. Row 2 | 0.5597 | -9.721 to 10.84 | No | ns | >0.9999 |
| Row 1 vs. Row 3 | -10.14 | -18.51 to -1.775 | Yes | ** | 0.0071 |
| Row 1 vs. Row 4 | -3.804 | -12.61 to 5.006 | No | ns | 0.8558 |
| Row 1 vs. Row 5 | -7.163 | -14.93 to 0.5997 | No | ns | 0.0915 |
| Row 1 vs. Row 6 | -17.50 | -25.87 to -9.134 | Yes | **** | <0.0001 |
| Row 1 vs. Row 7 | -5.715 | -14.08 to 2.655 | No | ns | 0.3941 |
| Row 2 vs. Row 3 | -10.70 | -21.61 to 0.1973 | No | ns | 0.0579 |
| Row 2 vs. Row 4 | -4.364 | -15.61 to 6.879 | No | ns | 0.9081 |
| Row 2 vs. Row 5 | -7.723 | -18.17 to 2.720 | No | ns | 0.2966 |
| Row 2 vs. Row 6 | -18.06 | -28.96 to -7.162 | Yes | **** | <0.0001 |
| Row 2 vs. Row 7 | -6.275 | -17.18 to 4.627 | No | ns | 0.6041 |
| Row 3 vs. Row 4 | 6.340 | -3.186 to 15.87 | No | ns | 0.4263 |
| Row 3 vs. Row 5 | 2.981 | -5.586 to 11.55 | No | ns | 0.9440 |
| Row 3 vs. Row 6 | -7.359 | -16.48 to 1.762 | No | ns | 0.2013 |
| Row 3 vs. Row 7 | 4.429 | -4.691 to 13.55 | No | ns | 0.7732 |
| Row 4 vs. Row 5 | -3.360 | -12.36 to 5.638 | No | ns | 0.9226 |
| Row 4 vs. Row 6 | -13.70 | -23.23 to -4.173 | Yes | *** | 0.0006 |
| Row 4 vs. Row 7 | -1.911 | -11.44 to 7.615 | No | ns | 0.9968 |
| Row 5 vs. Row 6 | -10.34 | -18.91 to -1.773 | Yes | ** | 0.0075 |
| Row 5 vs. Row 7 | 1.448 | -7.119 to 10.02 | No | ns | 0.9988 |
| Row 6 vs. Row 7 | 11.79 | 2.668 to 20.91 | Yes | ** | 0.0031 |
| ts | | | | | |
| Row 1 vs. Row 2 | -0.1219 | -10.56 to 10.32 | No | ns | >0.9999 |
| Row 1 vs. Row 3 | -9.698 | -21.53 to 2.132 | No | ns | 0.1857 |
| Row 1 vs. Row 4 | -13.56 | -22.55 to -4.558 | Yes | *** | 0.0003 |
| Row 1 vs. Row 5 | -13.90 | -22.29 to -5.508 | Yes | **** | <0.0001 |
| Row 1 vs. Row 6 | -5.155 | -14.15 to 3.843 | No | ns | 0.6095 |
| Row 1 vs. Row 7 | -9.732 | -18.50 to -0.9647 | Yes | * | 0.0191 |
| Row 2 vs. Row 3 | -9.576 | -23.19 to 4.039 | No | ns | 0.3573 |
| Row 2 vs. Row 4 | -13.43 | -24.68 to -2.191 | Yes | ** | 0.0085 |
| Row 2 vs. Row 5 | -13.78 | -24.54 to -3.014 | Yes | ** | 0.0035 |
| Row 2 vs. Row 6 | -5.033 | -16.28 to 6.210 | No | ns | 0.8333 |
| Row 2 vs. Row 7 | -9.610 | -20.67 to 1.449 | No | ns | 0.1344 |
| Row 3 vs. Row 4 | -3.858 | -16.40 to 8.684 | No | ns | 0.9691 |
| Row 3 vs. Row 5 | -4.202 | -16.32 to 7.913 | No | ns | 0.9448 |
| Row 3 vs. Row 6 | 4.543 | -7.998 to 17.09 | No | ns | 0.9325 |
| Row 3 vs. Row 7 | -0.03386 | -12.41 to 12.34 | No | ns | >0.9999 |
| Row 4 vs. Row 5 | -0.3441 | -9.713 to 9.025 | No | ns | >0.9999 |
| Row 4 vs. Row 6 | 8.401 | -1.514 to 18.32 | No | ns | 0.1555 |
| Row 4 vs. Row 7 | 3.824 | -5.882 to 13.53 | No | ns | 0.9019 |
| Row 5 vs. Row 6 | 8.745 | -0.6236 to 18.11 | No | ns | 0.0845 |
| Row 5 vs. Row 7 | 4.168 | -4.980 to 13.32 | No | ns | 0.8214 |
| Row 6 vs. Row 7 | -4.577 | -14.28 to 5.129 | No | ns | 0.7965 |

| Šidák's multiple comparisons test | Predicted (LS) mean diff. | 95.00% CI of diff. | Below threshold? | Summary | Adjusted P Value |
|---|---|---|---|---|---|
| wt - ts | | | | | |
| Row 1 | -1.672 | -8.737 to 5.393 | No | ns | 0.9942 |
| Row 2 | -2.353 | -13.66 to 8.958 | No | ns | 0.9974 |
| Row 3 | -1.226 | -12.36 to 9.910 | No | ns | >0.9999 |
| Row 4 | -11.42 | -20.45 to -2.401 | Yes | ** | 0.0052 |
| Row 5 | -8.409 | -16.05 to -0.7713 | Yes | * | 0.0223 |
| Row 6 | 10.68 | 2.008 to 19.35 | Yes | ** | 0.0071 |
| Row 7 | -5.689 | -14.16 to 2.783 | No | ns | 0.3975 |

| Test details | Predicted (LS) mean 1 | Predicted (LS) mea | Predicted (LS) mea | SE of diff. | N1 | N2 | t | DF |
|---|---|---|---|---|---|---|---|---|
| wt - ts | | | | | | | | |
| Row 1 | 19.24 | 20.92 | -1.672 | 2.599 | 19 | 17 | 0.6433 | 156.0 |
| Row 2 | 18.68 | 21.04 | -2.353 | 4.161 | 7 | 7 | 0.5656 | 156.0 |
| Row 3 | 29.39 | 30.61 | -1.226 | 4.096 | 13 | 5 | 0.2992 | 156.0 |
| Row 4 | 23.05 | 34.47 | -11.42 | 3.319 | 11 | 11 | 3.442 | 156.0 |
| Row 5 | 26.41 | 34.82 | -8.409 | 2.809 | 17 | 14 | 2.993 | 156.0 |
| Row 6 | 36.75 | 26.07 | 10.68 | 3.189 | 13 | 11 | 3.348 | 156.0 |
| Row 7 | 24.96 | 30.65 | -5.689 | 3.116 | 13 | 12 | 1.826 | 156.0 |

**Table S3A: Statistics for Figure 2**

#### Frequency

| Table Analyzed | Mean Frequency | | | | |
|---|---|---|---|---|---|
| **Two-way ANOVA** | Ordinary | | | | |
| Alpha | 0.05 | | | | |
| **Source of Variation** | **% of total variation** | **P value** | **P value summary** | **Significant?** | |
| Interaction | 2.787 | 0.3025 | ns | No | |
| Hours TTX | 34.91 | <0.0001 | **** | Yes | |
| Genotype | 0.1993 | 0.4717 | ns | No | |
| **ANOVA table** | **SS (Type III)** | **DF** | **MS** | **F (DFn, DFd)** | **P value** |
| Interaction | 1771 | 6 | 295.2 | F (6, 156) = 1.213 | P=0.3025 |
| Hours TTX | 22188 | 6 | 3698 | F (6, 156) = 15.19 | P<0.0001 |
| Genotype | 126.7 | 1 | 126.7 | F (1, 156) = 0.5204 | P=0.4717 |
| Residual | 37970 | 156 | 243.4 | | |
| **Difference between column means** | | | | | |
| Predicted (LS) mean of wt | 25.92 | | | | |
| Predicted (LS) mean of ts | 27.76 | | | | |
| Difference between predicted means | -1.843 | | | | |
| SE of difference | 2.555 | | | | |
| 95% CI of difference | -6.890 to 3.204 | | | | |
| **Data summary** | | | | | |
| Number of columns (Genotype) | 2 | | | | |
| Number of rows (Hours TTX) | 7 | | | | |
| Number of values | 170 | | | | |

| Test details | Predicted (LS) me | Predicted (LS) me | SE of diff | N1 | N2 | q | DF |
|---|---|---|---|---|---|---|---|
| wt | | | | | | | |
| Row 1 vs. Row 2 | 14.46 | -2.597 | 6.898 | 19 | 7 | 0.5323 | 156.0 |
| Row 1 vs. Row 3 | 40.09 | -28.22 | 5.615 | 19 | 13 | 7.107 | 156.0 |
| Row 1 vs. Row 4 | 34.99 | -23.12 | 5.911 | 19 | 11 | 5.532 | 156.0 |
| Row 1 vs. Row 5 | 12.98 | -1.112 | 5.208 | 19 | 17 | 0.3019 | 156.0 |
| Row 1 vs. Row 6 | 43.83 | -31.96 | 5.615 | 19 | 13 | 8.050 | 156.0 |
| Row 1 vs. Row 7 | 23.21 | -11.34 | 5.615 | 19 | 13 | 2.856 | 156.0 |
| Row 2 vs. Row 3 | 40.09 | -25.62 | 7.314 | 7 | 13 | 4.955 | 156.0 |
| Row 2 vs. Row 4 | 34.99 | -20.52 | 7.543 | 7 | 11 | 3.848 | 156.0 |
| Row 2 vs. Row 5 | 12.98 | 1.485 | 7.006 | 7 | 17 | 0.2997 | 156.0 |
| Row 2 vs. Row 6 | 43.83 | -29.37 | 7.314 | 7 | 13 | 5.678 | 156.0 |
| Row 2 vs. Row 7 | 23.21 | -8.742 | 7.314 | 7 | 13 | 1.690 | 156.0 |
| Row 3 vs. Row 4 | 34.99 | 5.099 | 6.391 | 13 | 11 | 1.128 | 156.0 |
| Row 3 vs. Row 5 | 12.98 | 27.11 | 5.748 | 13 | 17 | 6.670 | 156.0 |
| Row 3 vs. Row 6 | 43.83 | -3.744 | 6.119 | 13 | 13 | 0.8653 | 156.0 |
| Row 3 vs. Row 7 | 23.21 | 16.88 | 6.119 | 13 | 13 | 3.901 | 156.0 |
| Row 4 vs. Row 5 | 12.98 | 22.01 | 6.037 | 11 | 17 | 5.156 | 156.0 |
| Row 4 vs. Row 6 | 43.83 | -8.843 | 6.391 | 11 | 13 | 1.957 | 156.0 |
| Row 4 vs. Row 7 | 23.21 | 11.78 | 6.391 | 11 | 13 | 2.607 | 156.0 |
| Row 5 vs. Row 6 | 43.83 | -30.85 | 5.748 | 17 | 13 | 7.591 | 156.0 |
| Row 5 vs. Row 7 | 23.21 | -10.23 | 5.748 | 17 | 13 | 2.516 | 156.0 |
| Row 6 vs. Row 7 | 23.21 | 20.63 | 6.119 | 13 | 13 | 4.767 | 156.0 |
| ts | | | | | | | |
| Row 1 vs. Row 2 | 18.10 | -8.030 | 7.006 | 17 | 7 | 1.621 | 156.0 |
| Row 1 vs. Row 3 | 37.18 | -27.11 | 7.937 | 17 | 5 | 4.830 | 156.0 |
| Row 1 vs. Row 4 | 44.91 | -34.84 | 6.037 | 17 | 11 | 8.162 | 156.0 |
| Row 1 vs. Row 5 | 22.99 | -12.92 | 5.631 | 17 | 14 | 3.245 | 156.0 |
| Row 1 vs. Row 6 | 34.97 | -24.90 | 6.037 | 17 | 11 | 5.834 | 156.0 |
| Row 1 vs. Row 7 | 26.10 | -16.03 | 5.882 | 17 | 12 | 3.854 | 156.0 |
| Row 2 vs. Row 3 | 37.18 | -19.08 | 9.135 | 7 | 5 | 2.953 | 156.0 |
| Row 2 vs. Row 4 | 44.91 | -26.81 | 7.543 | 7 | 11 | 5.027 | 156.0 |
| Row 2 vs. Row 5 | 22.99 | -4.889 | 7.222 | 7 | 14 | 0.9574 | 156.0 |
| Row 2 vs. Row 6 | 34.97 | -16.87 | 7.543 | 7 | 11 | 3.164 | 156.0 |
| Row 2 vs. Row 7 | 26.10 | -8.000 | 7.420 | 7 | 12 | 1.525 | 156.0 |
| Row 3 vs. Row 4 | 44.91 | -7.738 | 8.415 | 5 | 11 | 1.300 | 156.0 |
| Row 3 vs. Row 5 | 22.99 | 14.19 | 8.128 | 5 | 14 | 2.468 | 156.0 |
| Row 3 vs. Row 6 | 34.97 | 2.201 | 8.415 | 5 | 11 | 0.3700 | 156.0 |
| Row 3 vs. Row 7 | 26.10 | 11.08 | 8.304 | 5 | 12 | 1.886 | 156.0 |
| Row 4 vs. Row 5 | 22.99 | 21.92 | 6.286 | 11 | 14 | 4.933 | 156.0 |
| Row 4 vs. Row 6 | 34.97 | 9.939 | 6.652 | 11 | 11 | 2.113 | 156.0 |
| Row 4 vs. Row 7 | 26.10 | 18.81 | 6.512 | 11 | 12 | 4.086 | 156.0 |
| Row 5 vs. Row 6 | 34.97 | -11.99 | 6.286 | 14 | 11 | 2.696 | 156.0 |
| Row 5 vs. Row 7 | 26.10 | -3.111 | 6.137 | 14 | 12 | 0.7168 | 156.0 |
| Row 6 vs. Row 7 | 26.10 | 8.874 | 6.512 | 11 | 12 | 1.927 | 156.0 |

| Tukey's multiple comparisons test | 95.00% CI of diff. | Below threshold? | Summary | Adjusted P Value |
|---|---|---|---|---|
| wt | | | | |
| Row 1 vs. Row 2 | -23.20 to 18.01 | No | ns | 0.9998 |
| Row 1 vs. Row 3 | -44.99 to -11.45 | Yes | **** | <0.0001 |
| Row 1 vs. Row 4 | -40.78 to -5.464 | Yes | ** | 0.0026 |
| Row 1 vs. Row 5 | -16.67 to 14.45 | No | ns | >0.9999 |
| Row 1 vs. Row 6 | -48.74 to -15.19 | Yes | **** | <0.0001 |
| Row 1 vs. Row 7 | -28.11 to 5.436 | No | ns | 0.4069 |
| Row 2 vs. Row 3 | -47.47 to -3.775 | Yes | * | 0.0105 |
| Row 2 vs. Row 4 | -43.06 to 2.008 | No | ns | 0.0999 |
| Row 2 vs. Row 5 | -19.44 to 22.41 | No | ns | >0.9999 |
| Row 2 vs. Row 6 | -51.22 to -7.519 | Yes | ** | 0.0018 |
| Row 2 vs. Row 7 | -30.59 to 13.11 | No | ns | 0.8951 |
| Row 3 vs. Row 4 | -13.99 to 24.19 | No | ns | 0.9849 |
| Row 3 vs. Row 5 | 9.938 to 44.28 | Yes | *** | 0.0001 |
| Row 3 vs. Row 6 | -22.02 to 14.54 | No | ns | 0.9964 |
| Row 3 vs. Row 7 | -1.398 to 35.16 | No | ns | 0.0910 |
| Row 4 vs. Row 5 | 3.976 to 40.04 | Yes | ** | 0.0065 |
| Row 4 vs. Row 6 | -27.94 to 10.25 | No | ns | 0.8098 |
| Row 4 vs. Row 7 | -7.310 to 30.87 | No | ns | 0.5209 |
| Row 5 vs. Row 6 | -48.02 to -13.68 | Yes | **** | <0.0001 |
| Row 5 vs. Row 7 | -27.40 to 6.944 | No | ns | 0.5641 |
| Row 6 vs. Row 7 | 2.346 to 38.91 | Yes | * | 0.0161 |
| ts | | | | |
| Row 1 vs. Row 2 | -28.96 to 12.90 | No | ns | 0.9126 |
| Row 1 vs. Row 3 | -50.82 to -3.396 | Yes | * | 0.0140 |
| Row 1 vs. Row 4 | -52.88 to -16.81 | Yes | **** | <0.0001 |
| Row 1 vs. Row 5 | -29.74 to 3.900 | No | ns | 0.2533 |
| Row 1 vs. Row 6 | -42.94 to -6.871 | Yes | ** | 0.0012 |
| Row 1 vs. Row 7 | -33.60 to 1.541 | No | ns | 0.0989 |
| Row 2 vs. Row 3 | -46.36 to 8.213 | No | ns | 0.3648 |
| Row 2 vs. Row 4 | -49.35 to -4.281 | Yes | ** | 0.0089 |
| Row 2 vs. Row 5 | -26.46 to 16.68 | No | ns | 0.9937 |
| Row 2 vs. Row 6 | -39.41 to 5.658 | No | ns | 0.2820 |
| Row 2 vs. Row 7 | -30.16 to 14.16 | No | ns | 0.9336 |
| Row 3 vs. Row 4 | -32.87 to 17.40 | No | ns | 0.9690 |
| Row 3 vs. Row 5 | -10.09 to 38.47 | No | ns | 0.5868 |
| Row 3 vs. Row 6 | -22.93 to 27.34 | No | ns | >0.9999 |
| Row 3 vs. Row 7 | -13.73 to 35.88 | No | ns | 0.8351 |
| Row 4 vs. Row 5 | 3.147 to 40.70 | Yes | * | 0.0111 |
| Row 4 vs. Row 6 | -9.933 to 29.81 | No | ns | 0.7480 |
| Row 4 vs. Row 7 | -0.6402 to 38.27 | No | ns | 0.0652 |
| Row 5 vs. Row 6 | -30.76 to 6.792 | No | ns | 0.4790 |
| Row 5 vs. Row 7 | -21.44 to 15.22 | No | ns | 0.9987 |
| Row 6 vs. Row 7 | -10.58 to 28.33 | No | ns | 0.8206 |

| Šidák's multiple comparisons test | Predicted (LS) mean diff. | 95.00% CI of diff. | Below threshold? | Summary | Adjusted P Value |
|---|---|---|---|---|---|
| wt - ts | | | | | |
| Row 1 | 1.797 | -12.36 to 15.96 | No | ns | 0.9999 |
| Row 2 | -3.636 | -26.31 to 19.03 | No | ns | 0.9995 |
| Row 3 | 2.911 | -19.41 to 25.23 | No | ns | 0.9999 |
| Row 4 | -9.926 | -28.01 to 8.159 | No | ns | 0.6455 |
| Row 5 | -10.01 | -25.32 to 5.296 | No | ns | 0.4309 |
| Row 6 | 8.857 | -8.518 to 26.23 | No | ns | 0.7236 |
| Row 7 | -2.895 | -19.87 to 14.08 | No | ns | 0.9993 |

| Test details | Predicted (LS) mean 1 | Predicted (LS) mean 2 | Predicted (LS) mean diff. | SE of diff. | N1 | N2 | t | DF |
|---|---|---|---|---|---|---|---|---|
| wt - ts | | | | | | | | |
| Row 1 | 11.87 | 10.07 | 1.797 | 5.208 | 19 | 17 | 0.3450 | 156.0 |
| Row 2 | 14.46 | 18.10 | -3.636 | 8.339 | 7 | 7 | 0.4361 | 156.0 |
| Row 3 | 40.09 | 37.18 | 2.911 | 8.210 | 13 | 5 | 0.3546 | 156.0 |
| Row 4 | 34.99 | 44.91 | -9.926 | 6.652 | 11 | 11 | 1.492 | 156.0 |
| Row 5 | 12.98 | 22.99 | -10.01 | 5.631 | 17 | 14 | 1.778 | 156.0 |
| Row 6 | 43.83 | 34.97 | 8.857 | 6.391 | 13 | 11 | 1.386 | 156.0 |
| Row 7 | 23.21 | 26.10 | -2.895 | 6.245 | 13 | 12 | 0.4635 | 156.0 |

**Table S3B: Statistics for Figure 2**

### Decay

| Table Analyzed | Mean Decay Tau | | | | |
|---|---|---|---|---|---|
| **Two-way ANOVA** | Ordinary | | | | |
| Alpha | 0.05 | | | | |
| **Source of Variation** | **% of total variation** | **P value** | **P value summary** | **Significant?** | |
| Interaction | 4.634 | 0.1860 | ns | No | |
| Hours TTX | 13.91 | 0.0003 | *** | Yes | |
| Genotype | 0.2847 | 0.4602 | ns | No | |
| **ANOVA table** | **SS (Type III)** | **DF** | **MS** | **F (DFn, DFd)** | **P value** |
| Interaction | 10.85 | 6 | 1.808 | F (6, 156) = 1.487 | P=0.1860 |
| Hours TTX | 32.56 | 6 | 5.426 | F (6, 156) = 4.461 | P=0.0003 |
| Genotype | 0.6667 | 1 | 0.6667 | F (1, 156) = 0.5481 | P=0.4602 |
| Residual | 189.7 | 156 | 1.216 | | |
| **Difference between column means** | | | | | |
| Predicted (LS) mean of wt | 4.830 | | | | |
| Predicted (LS) mean of ts | 4.696 | | | | |
| Difference between predicted means | 0.1337 | | | | |
| SE of difference | 0.1806 | | | | |
| 95% CI of difference | -0.2231 to 0.4905 | | | | |
| **Data summary** | | | | | |
| Number of columns (Genotype) | 2 | | | | |
| Number of rows (Hours TTX) | 7 | | | | |
| Number of values | 170 | | | | |

| Tukey's multiple compari | Predicted (LS) r | 95.00% CI of diff. | Below threshold? | Summary | Adjusted P Value | Test details | Predicted (LS) r | Predicted (LS) mea | Predicted (LS) me | SE of diff. | N1 | N2 | q | DF |
|---|---|---|---|---|---|---|---|---|---|---|---|---|---|---|
| wt | | | | | | wt | | | | | | | | |
| Row 1 vs. Row 2 | -0.3042 | -1.761 to 1.152 | No | ns | 0.9960 | Row 1 vs. Row 2 | 5.353 | 5.657 | -0.3042 | 0.4876 | 19 | 7 | 0.8822 | 156.0 |
| Row 1 vs. Row 3 | 0.8839 | -0.3019 to 2.070 | No | ns | 0.2875 | Row 1 vs. Row 3 | 5.353 | 4.469 | 0.8839 | 0.3970 | 19 | 13 | 3.149 | 156.0 |
| Row 1 vs. Row 4 | 0.6830 | -0.5652 to 1.931 | No | ns | 0.6602 | Row 1 vs. Row 4 | 5.353 | 4.670 | 0.6830 | 0.4178 | 19 | 11 | 2.312 | 156.0 |
| Row 1 vs. Row 5 | 0.8047 | -0.2951 to 1.905 | No | ns | 0.3093 | Row 1 vs. Row 5 | 5.353 | 4.548 | 0.8047 | 0.3682 | 19 | 17 | 3.091 | 156.0 |
| Row 1 vs. Row 6 | 1.292 | 0.1066 to 2.478 | Yes | * | 0.0230 | Row 1 vs. Row 6 | 5.353 | 4.060 | 1.292 | 0.3970 | 19 | 13 | 4.604 | 156.0 |
| Row 1 vs. Row 7 | 0.2991 | -0.8867 to 1.485 | No | ns | 0.9888 | Row 1 vs. Row 7 | 5.353 | 5.054 | 0.2991 | 0.3970 | 19 | 13 | 1.065 | 156.0 |
| Row 2 vs. Row 3 | 1.188 | -0.3564 to 2.733 | No | ns | 0.2517 | Row 2 vs. Row 3 | 5.657 | 4.469 | 1.188 | 0.5170 | 7 | 13 | 3.250 | 156.0 |
| Row 2 vs. Row 4 | 0.9872 | -0.6057 to 2.580 | No | ns | 0.5158 | Row 2 vs. Row 4 | 5.657 | 4.670 | 0.9872 | 0.5332 | 7 | 11 | 2.618 | 156.0 |
| Row 2 vs. Row 5 | 1.109 | -0.3706 to 2.588 | No | ns | 0.2811 | Row 2 vs. Row 5 | 5.657 | 4.548 | 1.109 | 0.4953 | 7 | 17 | 3.166 | 156.0 |
| Row 2 vs. Row 6 | 1.597 | 0.05206 to 3.141 | Yes | * | 0.0377 | Row 2 vs. Row 6 | 5.657 | 4.060 | 1.597 | 0.5170 | 7 | 13 | 4.367 | 156.0 |
| Row 2 vs. Row 7 | 0.6032 | -0.9412 to 2.148 | No | ns | 0.9055 | Row 2 vs. Row 7 | 5.657 | 5.054 | 0.6032 | 0.5170 | 7 | 13 | 1.650 | 156.0 |
| Row 3 vs. Row 4 | -0.2009 | -1.551 to 1.149 | No | ns | 0.9994 | Row 3 vs. Row 4 | 4.469 | 4.670 | -0.2009 | 0.4518 | 13 | 11 | 0.6288 | 156.0 |
| Row 3 vs. Row 5 | -0.07916 | -1.293 to 1.135 | No | ns | >0.9999 | Row 3 vs. Row 5 | 4.469 | 4.548 | -0.07916 | 0.4063 | 13 | 17 | 0.2755 | 156.0 |
| Row 3 vs. Row 6 | 0.4085 | -0.8837 to 1.701 | No | ns | 0.9646 | Row 3 vs. Row 6 | 4.469 | 4.060 | 0.4085 | 0.4326 | 13 | 13 | 1.335 | 156.0 |
| Row 3 vs. Row 7 | -0.5848 | -1.877 to 0.7074 | No | ns | 0.8261 | Row 3 vs. Row 7 | 4.469 | 5.054 | -0.5848 | 0.4326 | 13 | 13 | 1.912 | 156.0 |
| Row 4 vs. Row 5 | 0.1217 | -1.153 to 1.397 | No | ns | >0.9999 | Row 4 vs. Row 5 | 4.670 | 4.548 | 0.1217 | 0.4268 | 11 | 17 | 0.4034 | 156.0 |
| Row 4 vs. Row 6 | 0.6094 | -0.7403 to 1.959 | No | ns | 0.8276 | Row 4 vs. Row 6 | 4.670 | 4.060 | 0.6094 | 0.4518 | 11 | 13 | 1.907 | 156.0 |
| Row 4 vs. Row 7 | -0.3839 | -1.734 to 0.9657 | No | ns | 0.9791 | Row 4 vs. Row 7 | 4.670 | 5.054 | -0.3839 | 0.4518 | 11 | 13 | 1.202 | 156.0 |
| Row 5 vs. Row 6 | 0.4877 | -0.7262 to 1.701 | No | ns | 0.8932 | Row 5 vs. Row 6 | 4.548 | 4.060 | 0.4877 | 0.4063 | 17 | 13 | 1.697 | 156.0 |
| Row 5 vs. Row 7 | -0.5056 | -1.719 to 0.7082 | No | ns | 0.8755 | Row 5 vs. Row 7 | 4.548 | 5.054 | -0.5056 | 0.4063 | 17 | 13 | 1.760 | 156.0 |
| Row 6 vs. Row 7 | -0.9933 | -2.285 to 0.2989 | No | ns | 0.2525 | Row 6 vs. Row 7 | 4.060 | 5.054 | -0.9933 | 0.4326 | 13 | 13 | 3.247 | 156.0 |
| ts | | | | | | ts | | | | | | | | |
| Row 1 vs. Row 2 | 0.07820 | -1.401 to 1.558 | No | ns | >0.9999 | Row 1 vs. Row 2 | 5.467 | 5.389 | 0.07820 | 0.4953 | 17 | 7 | 0.2233 | 156.0 |
| Row 1 vs. Row 3 | 0.9962 | -0.6798 to 2.672 | No | ns | 0.5665 | Row 1 vs. Row 3 | 5.467 | 4.471 | 0.9962 | 0.5611 | 17 | 5 | 2.511 | 156.0 |
| Row 1 vs. Row 4 | 1.123 | -0.1514 to 2.398 | No | ns | 0.1234 | Row 1 vs. Row 4 | 5.467 | 4.343 | 1.123 | 0.4268 | 17 | 11 | 3.723 | 156.0 |
| Row 1 vs. Row 5 | 1.144 | -0.04492 to 2.333 | No | ns | 0.0677 | Row 1 vs. Row 5 | 5.467 | 4.323 | 1.144 | 0.3980 | 17 | 14 | 4.065 | 156.0 |
| Row 1 vs. Row 6 | 0.6218 | -0.6530 to 1.897 | No | ns | 0.7695 | Row 1 vs. Row 6 | 5.467 | 4.845 | 0.6218 | 0.4268 | 17 | 11 | 2.061 | 156.0 |
| Row 1 vs. Row 7 | 1.431 | 0.1891 to 2.673 | Yes | * | 0.0128 | Row 1 vs. Row 7 | 5.467 | 4.036 | 1.431 | 0.4158 | 17 | 12 | 4.868 | 156.0 |
| Row 2 vs. Row 3 | 0.9180 | -1.011 to 2.847 | No | ns | 0.7894 | Row 2 vs. Row 3 | 5.389 | 4.471 | 0.9180 | 0.6458 | 7 | 5 | 2.010 | 156.0 |
| Row 2 vs. Row 4 | 1.045 | -0.5476 to 2.638 | No | ns | 0.4442 | Row 2 vs. Row 4 | 5.389 | 4.343 | 1.045 | 0.5332 | 7 | 11 | 2.772 | 156.0 |
| Row 2 vs. Row 5 | 1.066 | -0.4592 to 2.591 | No | ns | 0.3651 | Row 2 vs. Row 5 | 5.389 | 4.323 | 1.066 | 0.5105 | 7 | 14 | 2.953 | 156.0 |
| Row 2 vs. Row 6 | 0.5436 | -1.049 to 2.136 | No | ns | 0.9489 | Row 2 vs. Row 6 | 5.389 | 4.845 | 0.5436 | 0.5332 | 7 | 11 | 1.442 | 156.0 |
| Row 2 vs. Row 7 | 1.353 | -0.2138 to 2.920 | No | ns | 0.1395 | Row 2 vs. Row 7 | 5.389 | 4.036 | 1.353 | 0.5245 | 7 | 12 | 3.648 | 156.0 |
| Row 3 vs. Row 4 | 0.1272 | -1.650 to 1.904 | No | ns | >0.9999 | Row 3 vs. Row 4 | 4.471 | 4.343 | 0.1272 | 0.5948 | 5 | 11 | 0.3024 | 156.0 |
| Row 3 vs. Row 5 | 0.1478 | -1.569 to 1.864 | No | ns | >0.9999 | Row 3 vs. Row 5 | 4.471 | 4.323 | 0.1478 | 0.5746 | 5 | 14 | 0.3638 | 156.0 |
| Row 3 vs. Row 6 | -0.3744 | -2.151 to 1.402 | No | ns | 0.9958 | Row 3 vs. Row 6 | 4.471 | 4.845 | -0.3744 | 0.5948 | 5 | 11 | 0.8902 | 156.0 |
| Row 3 vs. Row 7 | 0.4350 | -1.319 to 2.189 | No | ns | 0.9898 | Row 3 vs. Row 7 | 4.471 | 4.036 | 0.4350 | 0.5870 | 5 | 12 | 1.048 | 156.0 |
| Row 4 vs. Row 5 | 0.02062 | -1.307 to 1.348 | No | ns | >0.9999 | Row 4 vs. Row 5 | 4.343 | 4.323 | 0.02062 | 0.4444 | 11 | 14 | 0.06564 | 156.0 |
| Row 4 vs. Row 6 | -0.5016 | -1.906 to 0.9032 | No | ns | 0.9368 | Row 4 vs. Row 6 | 4.343 | 4.845 | -0.5016 | 0.4703 | 11 | 11 | 1.508 | 156.0 |
| Row 4 vs. Row 7 | 0.3078 | -1.067 to 1.683 | No | ns | 0.9941 | Row 4 vs. Row 7 | 4.343 | 4.036 | 0.3078 | 0.4604 | 11 | 12 | 0.9455 | 156.0 |
| Row 5 vs. Row 6 | -0.5222 | -1.850 to 0.8051 | No | ns | 0.9024 | Row 5 vs. Row 6 | 4.323 | 4.845 | -0.5222 | 0.4444 | 14 | 11 | 1.662 | 156.0 |
| Row 5 vs. Row 7 | 0.2872 | -1.009 to 1.583 | No | ns | 0.9944 | Row 5 vs. Row 7 | 4.323 | 4.036 | 0.2872 | 0.4339 | 14 | 12 | 0.9360 | 156.0 |
| Row 6 vs. Row 7 | 0.8094 | -0.5658 to 2.185 | No | ns | 0.5782 | Row 6 vs. Row 7 | 4.845 | 4.036 | 0.8094 | 0.4604 | 11 | 12 | 2.486 | 156.0 |

| Šidák's multiple comparisons test | Predicted (LS) mean d | 95.00% CI of diff. | Below threshold? | Summary | Adjusted P Value | Test details | Predicted (LS) mean 1 | Predicted (LS) mean 2 | Predicted (LS) mean diff. | SE of diff. | N1 | N2 | t | DF |
|---|---|---|---|---|---|---|---|---|---|---|---|---|---|---|
| wt - ts | | | | | | wt - ts | | | | | | | | |
| Row 1 | -0.1143 | -1.115 to 0.8866 | No | ns | >0.9999 | Row 1 | 5.353 | 5.467 | -0.1143 | 0.3682 | 19 | 17 | 0.3105 | 156.0 |
| Row 2 | 0.2681 | -1.334 to 1.871 | No | ns | 0.9994 | Row 2 | 5.657 | 5.389 | 0.2681 | 0.5895 | 7 | 7 | 0.4547 | 156.0 |
| Row 3 | -0.001935 | -1.580 to 1.576 | No | ns | >0.9999 | Row 3 | 4.469 | 4.471 | -0.001935 | 0.5804 | 13 | 5 | 0.00333 | 156.0 |
| Row 4 | 0.3262 | -0.9522 to 1.605 | No | ns | 0.9909 | Row 4 | 4.670 | 4.343 | 0.3262 | 0.4703 | 11 | 11 | 0.6936 | 156.0 |
| Row 5 | 0.2251 | -0.8570 to 1.307 | No | ns | 0.9974 | Row 5 | 4.548 | 4.323 | 0.2251 | 0.3980 | 17 | 14 | 0.5654 | 156.0 |
| Row 6 | -0.7848 | -2.013 to 0.4434 | No | ns | 0.4603 | Row 6 | 4.060 | 4.845 | -0.7848 | 0.4518 | 13 | 11 | 1.737 | 156.0 |
| Row 7 | 1.018 | -0.1823 to 2.218 | No | ns | 0.1470 | Row 7 | 5.054 | 4.036 | 1.018 | 0.4415 | 13 | 12 | 2.305 | 156.0 |

**Table S3C: Statistics for Figure 2**

| Two-way RM ANOVA | Matching: Across row | | | | |
|---|---|---|---|---|---|
| Assume sphericity? | Yes | | | | |
| Alpha | 0.05 | | | | |
| **Source of Variation** | **% of total variation** | **P value** | **P value summary** | **Significant?** | |
| Genotype x PhTx | 2.626 | 0.1640 | ns | No | |
| Genotype | 17.68 | 0.0372 | * | Yes | |
| PhTx | 6.917 | 0.0313 | * | Yes | |
| Subject | 50.73 | 0.0291 | * | Yes | |
| **ANOVA table** | **SS** | **DF** | **MS** | **F (DFn, DFd)** | **P value** |
| Genotype x PhTx | 85.48 | 1 | 85.48 | F (1, 15) = 2.141 | P=0.1640 |
| Genotype | 575.5 | 1 | 575.5 | F (1, 15) = 5.227 | P=0.0372 |
| PhTx | 225.2 | 1 | 225.2 | F (1, 15) = 5.640 | P=0.0313 |
| Subject | 1652 | 15 | 110.1 | F (15, 15) = 2.758 | P=0.0291 |
| Residual | 598.8 | 15 | 39.92 | | |
| **Difference between row means** | | | | | |
| Mean of wt 24h | 27.64 | | | | |
| Mean of ts 24h | 36.25 | | | | |
| Difference between means | -8.609 | | | | |
| SE of difference | 3.766 | | | | |
| 95% CI of difference | -16.64 to -0.5831 | | | | |
| **Difference between column means** | | | | | |
| Mean of -phtx | 34.64 | | | | |
| Mean of +phtx | 29.26 | | | | |
| Difference between means | 5.385 | | | | |
| SE of difference | 2.268 | | | | |
| 95% CI of difference | 0.5520 to 10.22 | | | | |
| **Interaction CI** | | | | | |
| Mean diff, A1 - B1 | 2.067 | | | | |
| Mean diff, A2 - B2 | 8.703 | | | | |
| (A1 -B1) - (A2 - B2) | -6.636 | | | | |
| 95% CI of difference | -16.30 to 3.030 | | | | |
| (B1 - A1) - (B2 - A2) | 6.636 | | | | |
| 95% CI of difference | -3.030 to 16.30 | | | | |
| **Data summary** | | | | | |
| Number of columns (PhTx) | 2 | | | | |
| Number of rows (Genotype) | 2 | | | | |
| Number of subjects (Subject) | 17 | | | | |
| Number of missing values | 0 | | | | |

| Šídák's multiple comparisons test | 95.00% CI of diff. | Below threshold? | Summary | Adjusted P Value | | | |
|---|---|---|---|---|---|---|---|
| -phtx - +phtx | | | | | | | |
| wt 24h | -6.992 to 11.13 | No | ns | 0.8230 | | | |
| ts 24h | 2.012 to 15.39 | Yes | * | 0.0112 | | | |
| **Test details** | **Predicted (LS) mean 2** | **Predicted (LS) mean diff.** | **SE of diff.** | **N1** | **N2** | **t** | **DF** |
| -phtx - +phtx | | | | | | | |
| wt 24h | 26.61 | 2.067 | 3.648 | 6 | 6 | 0.5667 | 15.00 |
| ts 24h | 31.90 | 8.703 | 2.694 | 11 | 11 | 3.230 | 15.00 |

**Table S4A: Statistics for Figure 3 – Amplitude 24 h TTX+PhTx**

| Two-way RM ANOVA | Matching: Across row | | | | |
|---|---|---|---|---|---|
| Assume sphericity? | Yes | | | | |
| Alpha | 0.05 | | | | |
| **Source of Variation** | **% of total variation** | **P value** | **P value summary** | **Significant?** | |
| Genotype x PhTx | 0.6154 | 0.1603 | ns | No | |
| Genotype | 3.857 | 0.3950 | ns | No | |
| PhTx | 3.973 | 0.0017 | ** | Yes | |
| Subject | 86.08 | <0.0001 | **** | Yes | |
| **ANOVA table** | **SS** | **DF** | **MS** | **F (DFn, DFd)** | **P value** |
| Genotype x PhTx | 22.71 | 1 | 22.71 | F (1, 17) = 2.156 | P=0.1603 |
| Genotype | 142.3 | 1 | 142.3 | F (1, 17) = 0.7616 | P=0.3950 |
| PhTx | 146.6 | 1 | 146.6 | F (1, 17) = 13.91 | P=0.0017 |
| Subject | 3176 | 17 | 186.8 | F (17, 17) = 17.73 | P<0.0001 |
| Residual | 179.1 | 17 | 10.53 | | |
| **Difference between row means** | | | | | |
| Mean of wt 48h | 38.51 | | | | |
| Mean of ts 48h | 34.59 | | | | |
| Difference between means | 3.919 | | | | |
| SE of difference | 4.491 | | | | |
| 95% CI of difference | -5.556 to 13.39 | | | | |
| **Difference between column means** | | | | | |
| Mean of -phtx | 38.54 | | | | |
| Mean of +phtx | 34.56 | | | | |
| Difference between means | 3.978 | | | | |
| SE of difference | 1.066 | | | | |
| 95% CI of difference | 1.728 to 6.228 | | | | |
| **Interaction CI** | | | | | |
| Mean diff, A1 - B1 | 5.544 | | | | |
| Mean diff, A2 - B2 | 2.412 | | | | |
| (A1 -B1) - (A2 - B2) | 3.131 | | | | |
| 95% CI of difference | -1.369 to 7.631 | | | | |
| (B1 - A1) - (B2 - A2) | -3.131 | | | | |
| 95% CI of difference | -7.631 to 1.369 | | | | |
| **Data summary** | | | | | |
| Number of columns (PhTx) | 2 | | | | |
| Number of rows (Genotype) | 2 | | | | |
| Number of subjects (Subject) | 19 | | | | |
| Number of missing values | 0 | | | | |

| Šidák's multiple comparisons test | Predicted (LS) mean diff. | 95.00% CI of diff. | Below threshold? | Summary | Adjusted P Value | | | |
|---|---|---|---|---|---|---|---|---|
| -phtx - +phtx | | | | | | | | |
| wt 48h | 5.544 | 2.150 to 8.937 | Yes | ** | 0.0018 | | | |
| ts 48h | 2.412 | -1.567 to 6.391 | No | ns | 0.2868 | | | |
| **Test details** | **Predicted (LS) mean 1** | **Predicted (LS) mea** | **Predicted (LS) mea** | **SE of diff.** | **N1** | **N2** | **t** | **DF** |
| -phtx - +phtx | | | | | | | | |
| wt 48h | 41.28 | 35.74 | 5.544 | 1.384 | 11 | 11 | 4.006 | 17.00 |
| ts 48h | 35.80 | 33.39 | 2.412 | 1.623 | 8 | 8 | 1.486 | 17.00 |

**Table S4B: Statistics for Figure 3 – Amplitude 48 h TTX+PhTx**

| Two-way RM ANOVA | Matching: Across row | | | | |
|---|---|---|---|---|---|
| Assume sphericity? | Yes | | | | |
| Alpha | 0.05 | | | | |
| **Source of Variation** | **% of total variation** | **P value** | **P value summary** | **Significant?** | |
| TTX Time x Drug | 3.760 | 0.0105 | * | Yes | |
| TTX Time | 1.032 | 0.6412 | ns | No | |
| Drug | 4.665 | 0.0052 | ** | Yes | |
| Genotype | 77.93 | <0.0001 | **** | Yes | |
| **ANOVA table** | **SS** | **DF** | **MS** | **F (DFn, DFd)** | **P value** |
| TTX Time x Drug | 2.386 | 1 | 2.386 | F (1, 17) = 8.278 | P=0.0105 |
| TTX Time | 0.6548 | 1 | 0.6548 | F (1, 17) = 0.2251 | P=0.6412 |
| Drug | 2.961 | 1 | 2.961 | F (1, 17) = 10.27 | P=0.0052 |
| Genotype | 49.46 | 17 | 2.909 | F (17, 17) = 10.09 | P<0.0001 |
| Residual | 4.900 | 17 | 0.2882 | | |
| **Difference between row means** | | | | | |
| Mean of WT | 4.850 | | | | |
| Mean of TiS | 4.568 | | | | |
| Difference between means | 0.2824 | | | | |
| SE of difference | 0.5952 | | | | |
| 95% CI of difference | -0.9735 to 1.538 | | | | |
| **Difference between column means** | | | | | |
| Mean of -phtx | 4.408 | | | | |
| Mean of +phtx | 5.009 | | | | |
| Difference between means | -0.6005 | | | | |
| SE of difference | 0.1874 | | | | |
| 95% CI of difference | -0.9958 to -0.2052 | | | | |
| **Interaction CI** | | | | | |
| Mean diff, A1 - B1 | -0.06143 | | | | |
| Mean diff, A2 - B2 | -1.140 | | | | |
| (A1 -B1) - (A2 - B2) | 1.078 | | | | |
| 95% CI of difference | 0.2875 to 1.869 | | | | |
| (B1 - A1) - (B2 - A2) | -1.078 | | | | |
| 95% CI of difference | -1.869 to -0.2875 | | | | |
| **Data summary** | | | | | |
| Number of columns (Drug) | 2 | | | | |
| Number of rows (TTX Time) | 2 | | | | |
| Number of subjects (Genotype) | 19 | | | | |
| Number of missing values | 0 | | | | |

| Šídák's multiple comparisons test | Predicted (LS) mean diff. | 95.00% CI of diff. | Below threshold? | Summary | Adjusted P Value | | | |
|---|---|---|---|---|---|---|---|---|
| -phtx - +phtx | | | | | | | | |
| WT | -0.06143 | -0.8214 to 0.6985 | No | ns | 0.9761 | | | |
| TiS | -1.140 | -1.656 to -0.6233 | Yes | **** | <0.0001 | | | |
| **Test details** | **Predicted (LS) mean 1** | **Predicted (LS) mea** | **Predicted (LS) mea** | **SE of diff.** | **N1** | **N2** | **t** | **DF** |
| -phtx - +phtx | | | | | | | | |
| WT | 4.819 | 4.881 | -0.06143 | 0.3100 | 6 | 6 | 0.1982 | 17.00 |
| TiS | 3.998 | 5.137 | -1.140 | 0.2106 | 13 | 13 | 5.412 | 17.00 |

**Table S4C: Statistics for Figure 3 – Decay Tau 24 h TTX+PhTx**

| Two-way RM ANOVA | Matching: Across row | | | | |
|---|---|---|---|---|---|
| Assume sphericity? | Yes | | | | |
| Alpha | 0.05 | | | | |
| **Source of Variation** | **% of total variation** | **P value** | **P value summary** | **Significant?** | |
| TTX Time x Drug | 4.306 | 0.0029 | ** | Yes | |
| TTX Time | 0.2954 | 0.8106 | ns | No | |
| Drug | 3.177 | 0.0084 | ** | Yes | |
| Genotype | 84.75 | <0.0001 | **** | Yes | |
| **ANOVA table** | **SS** | **DF** | **MS** | **F (DFn, DFd)** | **P value** |
| TTX Time x Drug | 0.7674 | 1 | 0.7674 | F (1, 17) = 12.04 | P=0.0029 |
| TTX Time | 0.05264 | 1 | 0.05264 | F (1, 17) = 0.05926 | P=0.8106 |
| Drug | 0.5662 | 1 | 0.5662 | F (1, 17) = 8.883 | P=0.0084 |
| Genotype | 15.10 | 17 | 0.8884 | F (17, 17) = 13.94 | P<0.0001 |
| Residual | 1.084 | 17 | 0.06374 | | |
| **Difference between row means** | | | | | |
| Mean of WT | 3.984 | | | | |
| Mean of TiS | 3.909 | | | | |
| Difference between means | 0.07539 | | | | |
| SE of difference | 0.3097 | | | | |
| 95% CI of difference | -0.5780 to 0.7288 | | | | |
| **Difference between column means** | | | | | |
| Mean of -phtx | 3.823 | | | | |
| Mean of +phtx | 4.070 | | | | |
| Difference between means | -0.2472 | | | | |
| SE of difference | 0.08295 | | | | |
| 95% CI of difference | -0.4222 to -0.07222 | | | | |
| **Interaction CI** | | | | | |
| Mean diff, A1 - B1 | -0.5351 | | | | |
| Mean diff, A2 - B2 | 0.04060 | | | | |
| (A1 -B1) - (A2 - B2) | -0.5757 | | | | |
| 95% CI of difference | -0.9257 to -0.2256 | | | | |
| (B1 - A1) - (B2 - A2) | 0.5757 | | | | |
| 95% CI of difference | 0.2256 to 0.9257 | | | | |
| **Data summary** | | | | | |
| Number of columns (Drug) | 2 | | | | |
| Number of rows (TTX Time) | 2 | | | | |
| Number of subjects (Genotype) | 19 | | | | |
| Number of missing values | 0 | | | | |

| Šidák's multiple comparisons test | Predicted (LS) mean diff. | 95.00% CI of diff. | Below threshold? | Summary | Adjusted P Value | | | |
|---|---|---|---|---|---|---|---|---|
| -phtx - +phtx | | | | | | | | |
| WT | -0.5351 | -0.7990 to -0.2711 | Yes | *** | 0.0002 | | | |
| TiS | 0.04060 | -0.2689 to 0.3501 | No | ns | 0.9383 | | | |
| **Test details** | **Predicted (LS) mean 1** | **Predicted (LS) mea** | **Predicted (LS) mea** | **SE of diff.** | **N1** | **N2** | **t** | **DF** |
| -phtx - +phtx | | | | | | | | |
| WT | 3.717 | 4.252 | -0.5351 | 0.1077 | 11 | 11 | 4.970 | 17.00 |
| TiS | 3.929 | 3.888 | 0.04060 | 0.1262 | 8 | 8 | 0.3216 | 17.00 |

**Table S4D: Statistics for Figure 3 – Decay Tau 48 h TTX+PhTx**

| Two-way ANOVA | Ordinary | | | | |
|---|---|---|---|---|---|
| Alpha | 0.05 | | | | |
| **Source of Variation** | **% of total variation** | **P value** | **P value summary** | **Significant?** | |
| Interaction | 1.877 | 0.0002 | *** | Yes | |
| Hours TTX | 12.89 | <0.0001 | **** | Yes | |
| Genotype | 0.6064 | 0.0033 | ** | Yes | |
| **ANOVA table** | **SS (Type III)** | **DF** | **MS** | **F (DFn, DFd)** | **P value** |
| Interaction | 545270884 | 6 | 90878481 | F (6, 1206) = 4 | P=0.0002 |
| Hours TTX | 3744214808 | 6 | 624035801 | F (6, 1206) = 3 | P<0.0001 |
| Genotype | 176167494 | 1 | 176167494 | F (1, 1206) = 8 | P=0.0033 |
| Residual | 24440846490 | 1206 | 20266042 | | |
| **Difference between column means** | | | | | |
| Predicted (LS) mean of wt | 10042 | | | | |
| Predicted (LS) mean of tis | 10817 | | | | |
| Difference between predicted means | -775.0 | | | | |
| SE of difference | 262.8 | | | | |
| 95% CI of difference | -1291 to -259.3 | | | | |
| **Data summary** | | | | | |
| Number of columns (Genotype) | 2 | | | | |
| Number of rows (Hours TTX) | 7 | | | | |
| Number of values | 1220 | | | | |

| Šidák's multiple comparisons test | Predicted (LS) mean diff. | 95.00% CI of diff. | Below threshold? | Summary | Adjusted P Value | | | |
|---|---|---|---|---|---|---|---|---|
| wt - tis | | | | | | | | |
| ttx00 | 558.0 | -1278 to 2394 | No | ns | 0.9764 | | | |
| ttx03 | -2223 | -4117 to -328.4 | Yes | * | 0.0115 | | | |
| ttx06 | -108.1 | -1971 to 1755 | No | ns | >0.9999 | | | |
| ttx12 | -2303 | -3819 to -787.0 | Yes | *** | 0.0003 | | | |
| ttx24 | -1691 | -3667 to 285.9 | No | ns | 0.1423 | | | |
| ttx48 | 1392 | -540.6 to 3324 | No | ns | 0.3177 | | | |
| ttx72 | -1050 | -3070 to 969.1 | No | ns | 0.7109 | | | |
| **Test details** | **Predicted (LS) mean 1** | **Predicted (LS) mean 2** | **Predicted (LS) mean diff.** | **SE of diff.** | **N1** | **N2** | **t** | **DF** |
| wt - tis | | | | | | | | |
| ttx00 | 8919 | 8361 | 558.0 | 683.4 | 103 | 75 | 0.8165 | 1206 |
| ttx03 | 6559 | 8782 | -2223 | 704.9 | 88 | 76 | 3.153 | 1206 |
| ttx06 | 10822 | 10930 | -108.1 | 693.2 | 88 | 81 | 0.1559 | 1206 |
| ttx12 | 11060 | 13363 | -2303 | 564.0 | 131 | 124 | 4.083 | 1206 |
| ttx24 | 9213 | 10903 | -1691 | 735.5 | 67 | 85 | 2.299 | 1206 |
| ttx48 | 13772 | 12380 | 1392 | 719.1 | 86 | 72 | 1.936 | 1206 |
| ttx72 | 9948 | 10998 | -1050 | 751.5 | 68 | 76 | 1.398 | 1206 |

| Tukey's multiple c | Predicted | 95.00% CI of diff | Below th | Summar | Adjusted P Value |
|---|---|---|---|---|---|
| wt | | | | | |
| ttx00 vs. ttx03 | 2360 | 429.7 to 4290 | Yes | ** | 0.0059 |
| ttx00 vs. ttx06 | -1903 | -3833 to 26.85 | No | ns | 0.0562 |
| ttx00 vs. ttx12 | -2141 | -3892 to -390.3 | Yes | ** | 0.0058 |
| ttx00 vs. ttx24 | -293.6 | -2380 to 1793 | No | ns | 0.9996 |
| ttx00 vs. ttx48 | -4853 | -6795 to -2911 | Yes | **** | <0.0001 |
| ttx00 vs. ttx72 | -1029 | -3106 to 1049 | No | ns | 0.7670 |
| ttx03 vs. ttx06 | -4263 | -6267 to -2258 | Yes | **** | <0.0001 |
| ttx03 vs. ttx12 | -4501 | -6333 to -2668 | Yes | **** | <0.0001 |
| ttx03 vs. ttx24 | -2653 | -4809 to -497.5 | Yes | ** | 0.0054 |
| ttx03 vs. ttx48 | -7213 | -9229 to -5197 | Yes | **** | <0.0001 |
| ttx03 vs. ttx72 | -3388 | -5535 to -1242 | Yes | **** | <0.0001 |
| ttx06 vs. ttx12 | -238.1 | -2071 to 1594 | No | ns | 0.9998 |
| ttx06 vs. ttx24 | 1610 | -546.1 to 3765 | No | ns | 0.2934 |
| ttx06 vs. ttx48 | -2950 | -4966 to -933.8 | Yes | *** | 0.0003 |
| ttx06 vs. ttx72 | 874.3 | -1272 to 3021 | No | ns | 0.8932 |
| ttx12 vs. ttx24 | 1848 | -149.3 to 3845 | No | ns | 0.0912 |
| ttx12 vs. ttx48 | -2712 | -4557 to -866.5 | Yes | *** | 0.0003 |
| ttx12 vs. ttx72 | 1112 | -874.8 to 3100 | No | ns | 0.6475 |
| ttx24 vs. ttx48 | -4559 | -6726 to -2393 | Yes | **** | <0.0001 |
| ttx24 vs. ttx72 | -735.2 | -3024 to 1553 | No | ns | 0.9644 |
| ttx48 vs. ttx72 | 3824 | 1667 to 5982 | Yes | **** | <0.0001 |
| tis | | | | | |
| ttx00 vs. ttx03 | -421.2 | -2585 to 1743 | No | ns | 0.9975 |
| ttx00 vs. ttx06 | -2569 | -4700 to -438.6 | Yes | ** | 0.0070 |
| ttx00 vs. ttx12 | -5002 | -6947 to -3057 | Yes | **** | <0.0001 |
| ttx00 vs. ttx24 | -2542 | -4648 to -435.8 | Yes | ** | 0.0069 |
| ttx00 vs. ttx48 | -4019 | -6213 to -1825 | Yes | **** | <0.0001 |
| ttx00 vs. ttx72 | -2637 | -4801 to -473.2 | Yes | ** | 0.0061 |
| ttx03 vs. ttx06 | -2148 | -4271 to -24.77 | Yes | * | 0.0453 |
| ttx03 vs. ttx12 | -4581 | -6518 to -2644 | Yes | **** | <0.0001 |
| ttx03 vs. ttx24 | -2121 | -4220 to -22.05 | Yes | * | 0.0457 |
| ttx03 vs. ttx48 | -3598 | -5784 to -1411 | Yes | **** | <0.0001 |
| ttx03 vs. ttx72 | -2216 | -4373 to -59.18 | Yes | * | 0.0395 |
| ttx06 vs. ttx12 | -2433 | -4332 to -533.3 | Yes | ** | 0.0031 |
| ttx06 vs. ttx24 | 27.04 | -2037 to 2091 | No | ns | >0.9999 |
| ttx06 vs. ttx48 | -1450 | -3603 to 703.6 | No | ns | 0.4226 |
| ttx06 vs. ttx72 | -67.96 | -2191 to 2055 | No | ns | >0.9999 |
| ttx12 vs. ttx24 | 2460 | 587.6 to 4332 | Yes | ** | 0.0021 |
| ttx12 vs. ttx48 | 982.9 | -987.0 to 2953 | No | ns | 0.7606 |
| ttx12 vs. ttx72 | 2365 | 427.9 to 4302 | Yes | ** | 0.0060 |
| ttx24 vs. ttx48 | -1477 | -3606 to 652.6 | No | ns | 0.3847 |
| ttx24 vs. ttx72 | -95.00 | -2194 to 2004 | No | ns | >0.9999 |
| ttx48 vs. ttx72 | 1382 | -804.6 to 3568 | No | ns | 0.5031 |

| Test details | Predicted | Predicted (LS) m | Predicted | SE of dif | N1 | N2 | q | DF |
|---|---|---|---|---|---|---|---|---|
| wt | | | | | | | | |
| ttx00 vs. ttx03 | 8919 | 6559 | 2360 | 653.5 | 103 | 88 | 5.106 | 1206 |
| ttx00 vs. ttx06 | 8919 | 10822 | -1903 | 653.5 | 103 | 88 | 4.119 | 1206 |
| ttx00 vs. ttx12 | 8919 | 11060 | -2141 | 592.8 | 103 | 131 | 5.108 | 1206 |
| ttx00 vs. ttx24 | 8919 | 9213 | -293.6 | 706.6 | 103 | 67 | 0.5876 | 1206 |
| ttx00 vs. ttx48 | 8919 | 13772 | -4853 | 657.6 | 103 | 86 | 10.44 | 1206 |
| ttx00 vs. ttx72 | 8919 | 9948 | -1029 | 703.4 | 103 | 68 | 2.068 | 1206 |
| ttx03 vs. ttx06 | 6559 | 10822 | -4263 | 678.7 | 88 | 88 | 8.883 | 1206 |
| ttx03 vs. ttx12 | 6559 | 11060 | -4501 | 620.5 | 88 | 131 | 10.26 | 1206 |
| ttx03 vs. ttx24 | 6559 | 9213 | -2653 | 729.9 | 88 | 67 | 5.141 | 1206 |
| ttx03 vs. ttx48 | 6559 | 13772 | -7213 | 682.6 | 88 | 86 | 14.94 | 1206 |
| ttx03 vs. ttx72 | 6559 | 9948 | -3388 | 726.9 | 88 | 68 | 6.593 | 1206 |
| ttx06 vs. ttx12 | 10822 | 11060 | -238.1 | 620.5 | 88 | 131 | 0.5426 | 1206 |
| ttx06 vs. ttx24 | 10822 | 9213 | 1610 | 729.9 | 88 | 67 | 3.119 | 1206 |
| ttx06 vs. ttx48 | 10822 | 13772 | -2950 | 682.6 | 88 | 86 | 6.111 | 1206 |
| ttx06 vs. ttx72 | 10822 | 9948 | 874.3 | 726.9 | 88 | 68 | 1.701 | 1206 |
| ttx12 vs. ttx24 | 11060 | 9213 | 1848 | 676.2 | 131 | 67 | 3.864 | 1206 |
| ttx12 vs. ttx48 | 11060 | 13772 | -2712 | 624.8 | 131 | 86 | 6.138 | 1206 |
| ttx12 vs. ttx72 | 11060 | 9948 | 1112 | 672.9 | 131 | 68 | 2.338 | 1206 |
| ttx24 vs. ttx48 | 9213 | 13772 | -4559 | 733.6 | 67 | 86 | 8.790 | 1206 |
| ttx24 vs. ttx72 | 9213 | 9948 | -735.2 | 774.9 | 67 | 68 | 1.342 | 1206 |
| ttx48 vs. ttx72 | 13772 | 9948 | 3824 | 730.5 | 86 | 68 | 7.403 | 1206 |
| tis | | | | | | | | |
| ttx00 vs. ttx03 | 8361 | 8782 | -421.2 | 732.7 | 75 | 76 | 0.8129 | 1206 |
| ttx00 vs. ttx06 | 8361 | 10930 | -2569 | 721.4 | 75 | 81 | 5.037 | 1206 |
| ttx00 vs. ttx12 | 8361 | 13363 | -5002 | 658.5 | 75 | 124 | 10.74 | 1206 |
| ttx00 vs. ttx24 | 8361 | 10903 | -2542 | 713.2 | 75 | 85 | 5.041 | 1206 |
| ttx00 vs. ttx48 | 8361 | 12380 | -4019 | 742.8 | 75 | 72 | 7.652 | 1206 |
| ttx00 vs. ttx72 | 8361 | 10998 | -2637 | 732.7 | 75 | 76 | 5.090 | 1206 |
| ttx03 vs. ttx06 | 8782 | 10930 | -2148 | 718.9 | 76 | 81 | 4.225 | 1206 |
| ttx03 vs. ttx12 | 8782 | 13363 | -4581 | 655.8 | 76 | 124 | 9.878 | 1206 |
| ttx03 vs. ttx24 | 8782 | 10903 | -2121 | 710.7 | 76 | 85 | 4.221 | 1206 |
| ttx03 vs. ttx48 | 8782 | 12380 | -3598 | 740.4 | 76 | 72 | 6.873 | 1206 |
| ttx03 vs. ttx72 | 8782 | 10998 | -2216 | 730.3 | 76 | 76 | 4.291 | 1206 |
| ttx06 vs. ttx12 | 10930 | 13363 | -2433 | 643.1 | 81 | 124 | 5.349 | 1206 |
| ttx06 vs. ttx24 | 10930 | 10903 | 27.04 | 699.0 | 81 | 85 | 0.05471 | 1206 |
| ttx06 vs. ttx48 | 10930 | 12380 | -1450 | 729.2 | 81 | 72 | 2.812 | 1206 |
| ttx06 vs. ttx72 | 10930 | 10998 | -67.96 | 718.9 | 81 | 76 | 0.1337 | 1206 |
| ttx12 vs. ttx24 | 13363 | 10903 | 2460 | 633.9 | 124 | 85 | 5.487 | 1206 |
| ttx12 vs. ttx48 | 13363 | 12380 | 982.9 | 667.0 | 124 | 72 | 2.084 | 1206 |
| ttx12 vs. ttx72 | 13363 | 10998 | 2365 | 655.8 | 124 | 76 | 5.099 | 1206 |
| ttx24 vs. ttx48 | 10903 | 12380 | -1477 | 721.0 | 85 | 72 | 2.897 | 1206 |
| ttx24 vs. ttx72 | 10903 | 10998 | -95.00 | 710.7 | 85 | 76 | 0.1890 | 1206 |
| ttx48 vs. ttx72 | 12380 | 10998 | 1382 | 740.4 | 72 | 76 | 2.640 | 1206 |

**Table S5A: Statistics for Figure 3 – sGluA1 synaptically localized**

| Source of Variation | % of total variation | P value | P value summary | Significant? | |
|---|---|---|---|---|---|
| Interaction | 2.115 | <0.0001 | **** | Yes | |
| Hours TTX | 15.94 | <0.0001 | **** | Yes | |
| Genotype | 0.3264 | 0.0289 | * | Yes | |
| **ANOVA table** | **SS (Type III)** | **DF** | **MS** | **F (DFn, DFd)** | **P value** |
| Interaction | 256389421 | 6 | 42731570 | F (6, 1188) = 5.168 | P<0.0001 |
| Hours TTX | 1932895459 | 6 | 322149243 | F (6, 1188) = 38.96 | P<0.0001 |
| Genotype | 39576509 | 1 | 39576509 | F (1, 1188) = 4.786 | P=0.0289 |
| Residual | 9823030227 | 1188 | 8268544 | | |
| **Difference between column means** | | | | | |
| Predicted (LS) mean of wt | 6892 | | | | |
| Predicted (LS) mean of tis | 7264 | | | | |
| Difference between predicted means | -371.2 | | | | |
| SE of difference | 169.6 | | | | |
| 95% CI of difference | -704.0 to -38.31 | | | | |
| **Data summary** | | | | | |
| Number of columns (Genotype) | 2 | | | | |
| Number of rows (Hours TTX) | 7 | | | | |
| Number of values | 1202 | | | | |

| Šidák's multiple comparisons test | Predicted (LS) mean diff. | 95.00% CI of diff. | Below threshold? | Summary | Adjusted P Value | | | |
|---|---|---|---|---|---|---|---|---|
| wt - tis | | | | | | | | |
| ttx00 | -123.8 | -1299 to 1052 | No | ns | >0.9999 | | | |
| ttx03 | -680.2 | -1890 to 529.9 | No | ns | 0.6262 | | | |
| ttx06 | -122.8 | -1313 to 1067 | No | ns | >0.9999 | | | |
| ttx12 | -339.0 | -1307 to 629.2 | No | ns | 0.9493 | | | |
| ttx24 | -1707 | -3048 to -365.9 | Yes | ** | 0.0045 | | | |
| ttx48 | 1646 | 411.3 to 2880 | Yes | ** | 0.0025 | | | |
| ttx72 | -1271 | -2561 to 18.73 | No | ns | 0.0560 | | | |
| **Test details** | **Predicted (LS) mean 1** | **Predicted (LS) mean 2** | **Predicted (LS) mean diff.** | **SE of diff.** | **N1** | **N2** | **t** | **DF** |
| wt - tis | | | | | | | | |
| ttx00 | 5541 | 5665 | -123.8 | 437.4 | 102 | 75 | 0.2831 | 1188 |
| ttx03 | 4694 | 5374 | -680.2 | 450.3 | 88 | 76 | 1.511 | 1188 |
| ttx06 | 7222 | 7345 | -122.8 | 442.8 | 88 | 81 | 0.2773 | 1188 |
| ttx12 | 7712 | 8051 | -339.0 | 360.3 | 131 | 124 | 0.9410 | 1188 |
| ttx24 | 5829 | 7536 | -1707 | 498.9 | 59 | 76 | 3.421 | 1188 |
| ttx48 | 9704 | 8059 | 1646 | 459.3 | 86 | 72 | 3.583 | 1188 |
| ttx72 | 7544 | 8815 | -1271 | 480.0 | 68 | 76 | 2.648 | 1188 |

| Tukey's multiple comparisons test | Predicted (LS | 95.00% CI of diff. | Below threshold? | Summary | Adjusted P Value |
|---|---|---|---|---|---|
| wt | | | | | |
| ttx00 vs. ttx03 | 846.7 | -388.9 to 2082 | No | ns | 0.4000 |
| ttx00 vs. ttx06 | -1681 | -2917 to -445.7 | Yes | ** | 0.0012 |
| ttx00 vs. ttx12 | -2172 | -3293 to -1050 | Yes | **** | <0.0001 |
| ttx00 vs. ttx24 | -288.1 | -1677 to 1101 | No | ns | 0.9964 |
| ttx00 vs. ttx48 | -4164 | -5407 to -2920 | Yes | **** | <0.0001 |
| ttx00 vs. ttx72 | -2003 | -3332 to -673.3 | Yes | *** | 0.0002 |
| ttx03 vs. ttx06 | -2528 | -3808 to -1248 | Yes | **** | <0.0001 |
| ttx03 vs. ttx12 | -3018 | -4189 to -1848 | Yes | **** | <0.0001 |
| ttx03 vs. ttx24 | -1135 | -2564 to 294.2 | No | ns | 0.2234 |
| ttx03 vs. ttx48 | -5010 | -6298 to -3723 | Yes | **** | <0.0001 |
| ttx03 vs. ttx72 | -2850 | -4221 to -1478 | Yes | **** | <0.0001 |
| ttx06 vs. ttx12 | -490.3 | -1661 to 680.2 | No | ns | 0.8796 |
| ttx06 vs. ttx24 | 1393 | -35.80 to 2822 | No | ns | 0.0616 |
| ttx06 vs. ttx48 | -2482 | -3770 to -1195 | Yes | **** | <0.0001 |
| ttx06 vs. ttx72 | -321.5 | -1693 to 1050 | No | ns | 0.9930 |
| ttx12 vs. ttx24 | 1883 | 551.9 to 3215 | Yes | *** | 0.0006 |
| ttx12 vs. ttx48 | -1992 | -3171 to -813.4 | Yes | **** | <0.0001 |
| ttx12 vs. ttx72 | 168.7 | -1101 to 1438 | No | ns | 0.9997 |
| ttx24 vs. ttx48 | -3875 | -5311 to -2440 | Yes | **** | <0.0001 |
| ttx24 vs. ttx72 | -1715 | -3226 to -203.8 | Yes | * | 0.0145 |
| ttx48 vs. ttx72 | 2161 | 782.6 to 3539 | Yes | **** | <0.0001 |
| tis | | | | | |
| ttx00 vs. ttx03 | 290.3 | -1092 to 1673 | No | ns | 0.9962 |
| ttx00 vs. ttx06 | -1680 | -3041 to -319.3 | Yes | ** | 0.0052 |
| ttx00 vs. ttx12 | -2387 | -3629 to -1144 | Yes | **** | <0.0001 |
| ttx00 vs. ttx24 | -1871 | -3253 to -488.8 | Yes | ** | 0.0013 |
| ttx00 vs. ttx48 | -2394 | -3795 to -992.8 | Yes | **** | <0.0001 |
| ttx00 vs. ttx72 | -3150 | -4532 to -1768 | Yes | **** | <0.0001 |
| ttx03 vs. ttx06 | -1971 | -3327 to -614.3 | Yes | *** | 0.0004 |
| ttx03 vs. ttx12 | -2677 | -3914 to -1440 | Yes | **** | <0.0001 |
| ttx03 vs. ttx24 | -2161 | -3539 to -783.7 | Yes | **** | <0.0001 |
| ttx03 vs. ttx48 | -2684 | -4081 to -1288 | Yes | **** | <0.0001 |
| ttx03 vs. ttx72 | -3441 | -4818 to -2063 | Yes | **** | <0.0001 |
| ttx06 vs. ttx12 | -706.5 | -1920 to 506.8 | No | ns | 0.6028 |
| ttx06 vs. ttx24 | -190.8 | -1547 to 1165 | No | ns | 0.9996 |
| ttx06 vs. ttx48 | -713.7 | -2089 to 661.8 | No | ns | 0.7252 |
| ttx06 vs. ttx72 | -1470 | -2826 to -113.7 | Yes | * | 0.0237 |
| ttx12 vs. ttx24 | 515.7 | -721.5 to 1753 | No | ns | 0.8820 |
| ttx12 vs. ttx48 | -7.229 | -1266 to 1251 | No | ns | >0.9999 |
| ttx12 vs. ttx72 | -763.5 | -2001 to 473.7 | No | ns | 0.5329 |
| ttx24 vs. ttx48 | -522.9 | -1920 to 873.7 | No | ns | 0.9265 |
| ttx24 vs. ttx72 | -1279 | -2657 to 98.49 | No | ns | 0.0889 |
| ttx48 vs. ttx72 | -756.2 | -2153 to 640.4 | No | ns | 0.6830 |

| Test details | Predicted (LS | Predicted (LS) me | Predicted (LS) mear | SE of diff. | N1 | N2 | q | DF |
|---|---|---|---|---|---|---|---|---|
| wt | | | | | | | | |
| ttx00 vs. ttx03 | 5541 | 4694 | 846.7 | 418.4 | 102 | 88 | 2.862 | 1188 |
| ttx00 vs. ttx06 | 5541 | 7222 | -1681 | 418.4 | 102 | 88 | 5.683 | 1188 |
| ttx00 vs. ttx12 | 5541 | 7712 | -2172 | 379.7 | 102 | 131 | 8.088 | 1188 |
| ttx00 vs. ttx24 | 5541 | 5829 | -288.1 | 470.3 | 102 | 59 | 0.8663 | 1188 |
| ttx00 vs. ttx48 | 5541 | 9704 | -4164 | 421.0 | 102 | 86 | 13.99 | 1188 |
| ttx00 vs. ttx72 | 5541 | 7544 | -2003 | 450.2 | 102 | 68 | 6.292 | 1188 |
| ttx03 vs. ttx06 | 4694 | 7222 | -2528 | 433.5 | 88 | 88 | 8.247 | 1188 |
| ttx03 vs. ttx12 | 4694 | 7712 | -3018 | 396.3 | 88 | 131 | 10.77 | 1188 |
| ttx03 vs. ttx24 | 4694 | 5829 | -1135 | 483.8 | 88 | 59 | 3.317 | 1188 |
| ttx03 vs. ttx48 | 4694 | 9704 | -5010 | 436.0 | 88 | 86 | 16.25 | 1188 |
| ttx03 vs. ttx72 | 4694 | 7544 | -2850 | 464.3 | 88 | 68 | 8.680 | 1188 |
| ttx06 vs. ttx12 | 7222 | 7712 | -490.3 | 396.3 | 88 | 131 | 1.749 | 1188 |
| ttx06 vs. ttx24 | 7222 | 5829 | 1393 | 483.8 | 88 | 59 | 4.072 | 1188 |
| ttx06 vs. ttx48 | 7222 | 9704 | -2482 | 436.0 | 88 | 86 | 8.051 | 1188 |
| ttx06 vs. ttx72 | 7222 | 7544 | -321.5 | 464.3 | 88 | 68 | 0.9794 | 1188 |
| ttx12 vs. ttx24 | 7712 | 5829 | 1883 | 450.8 | 131 | 59 | 5.908 | 1188 |
| ttx12 vs. ttx48 | 7712 | 9704 | -1992 | 399.1 | 131 | 86 | 7.059 | 1188 |
| ttx12 vs. ttx72 | 7712 | 7544 | 168.7 | 429.8 | 131 | 68 | 0.5552 | 1188 |
| ttx24 vs. ttx48 | 5829 | 9704 | -3875 | 486.1 | 59 | 86 | 11.27 | 1188 |
| ttx24 vs. ttx72 | 5829 | 7544 | -1715 | 511.6 | 59 | 68 | 4.740 | 1188 |
| ttx48 vs. ttx72 | 9704 | 7544 | 2161 | 466.6 | 86 | 68 | 6.549 | 1188 |
| tis | | | | | | | | |
| ttx00 vs. ttx03 | 5665 | 5374 | 290.3 | 468.0 | 75 | 76 | 0.8773 | 1188 |
| ttx00 vs. ttx06 | 5665 | 7345 | -1680 | 460.8 | 75 | 81 | 5.157 | 1188 |
| ttx00 vs. ttx12 | 5665 | 8051 | -2387 | 420.6 | 75 | 124 | 8.025 | 1188 |
| ttx00 vs. ttx24 | 5665 | 7536 | -1871 | 468.0 | 75 | 76 | 5.654 | 1188 |
| ttx00 vs. ttx48 | 5665 | 8059 | -2394 | 474.4 | 75 | 72 | 7.136 | 1188 |
| ttx00 vs. ttx72 | 5665 | 8815 | -3150 | 468.0 | 75 | 76 | 9.519 | 1188 |
| ttx03 vs. ttx06 | 5374 | 7345 | -1971 | 459.2 | 76 | 81 | 6.069 | 1188 |
| ttx03 vs. ttx12 | 5374 | 8051 | -2677 | 418.9 | 76 | 124 | 9.038 | 1188 |
| ttx03 vs. ttx24 | 5374 | 7536 | -2161 | 466.5 | 76 | 76 | 6.553 | 1188 |
| ttx03 vs. ttx48 | 5374 | 8059 | -2684 | 472.9 | 76 | 72 | 8.027 | 1188 |
| ttx03 vs. ttx72 | 5374 | 8815 | -3441 | 466.5 | 76 | 76 | 10.43 | 1188 |
| ttx06 vs. ttx12 | 7345 | 8051 | -706.5 | 410.8 | 81 | 124 | 2.432 | 1188 |
| ttx06 vs. ttx24 | 7345 | 7536 | -190.8 | 459.2 | 81 | 76 | 0.5876 | 1188 |
| ttx06 vs. ttx48 | 7345 | 8059 | -713.7 | 465.7 | 81 | 72 | 2.167 | 1188 |
| ttx06 vs. ttx72 | 7345 | 8815 | -1470 | 459.2 | 81 | 76 | 4.527 | 1188 |
| ttx12 vs. ttx24 | 8051 | 7536 | 515.7 | 418.9 | 124 | 76 | 1.741 | 1188 |
| ttx12 vs. ttx48 | 8051 | 8059 | -7.229 | 426.1 | 124 | 72 | 0.02400 | 1188 |
| ttx12 vs. ttx72 | 8051 | 8815 | -763.5 | 418.9 | 124 | 76 | 2.577 | 1188 |
| ttx24 vs. ttx48 | 7536 | 8059 | -522.9 | 472.9 | 76 | 72 | 1.564 | 1188 |
| ttx24 vs. ttx72 | 7536 | 8815 | -1279 | 466.5 | 76 | 76 | 3.878 | 1188 |
| ttx48 vs. ttx72 | 8059 | 8815 | -756.2 | 472.9 | 72 | 76 | 2.262 | 1188 |

**Table S5B: Statistics for Figure 3 – sGluA1 dendritic shaft**

#### Amplitude

| Two-way ANOVA | Ordinary | | | | |
|---|---|---|---|---|---|
| Alpha | 0.05 | | | | |
| **Source of Variation** | **% of total variation** | **P value** | **P value summary** | **Significant?** | |
| Interaction | 2.001 | 0.6339 | ns | No | |
| Hours FK506 | 30.15 | <0.0001 | **** | Yes | |
| Genotype | 8.867 | 0.0012 | ** | Yes | |
| **ANOVA table** | **SS (Type III)** | **DF** | **MS** | **F (DFn, DFd)** | **P value** |
| Interaction | 146.0 | 4 | 36.49 | F (4, 76) = 0.6424 | P=0.6339 |
| Hours FK506 | 2199 | 4 | 549.8 | F (4, 76) = 9.678 | P<0.0001 |
| Genotype | 646.7 | 1 | 646.7 | F (1, 76) = 11.38 | P=0.0012 |
| Residual | 4317 | 76 | 56.81 | | |
| **Difference between column means** | | | | | |
| Predicted (LS) mean of wt | 21.21 | | | | |
| Predicted (LS) mean of ts | 26.80 | | | | |
| Difference between predicted means | -5.595 | | | | |
| SE of difference | 1.658 | | | | |
| 95% CI of difference | -8.897 to -2.292 | | | | |
| **Data summary** | | | | | |
| Number of columns (Genotype) | 2 | | | | |
| Number of rows (Hours FK506) | 5 | | | | |
| Number of values | 86 | | | | |

| Šidák's multiple comparisons test | Predicted (LS) mean diff. | 95.00% CI of diff. | Below threshold? | Summary | Adjusted P Value |
|---|---|---|---|---|---|
| wt - ts | | | | | |
| FK506 0h | -1.169 | -10.23 to 7.894 | No | ns | 0.9987 |
| FK506 3h | -5.086 | -14.45 to 4.275 | No | ns | 0.5728 |
| FK506 6h | -5.060 | -13.98 to 3.865 | No | ns | 0.5280 |
| FK506 12h | -8.066 | -18.34 to 2.211 | No | ns | 0.1934 |
| FK506 24h | -8.594 | -19.64 to 2.453 | No | ns | 0.2009 |

| Test details | Predicted (LS) mean 1 | Predicted (LS) mean 2 | Predicted (LS) mean diff. | SE of diff. | N1 | N2 | t | DF |
|---|---|---|---|---|---|---|---|---|
| wt - ts | | | | | | | | |
| FK506 0h | 15.75 | 16.92 | -1.169 | 3.440 | 12 | 8 | 0.3398 | 76.00 |
| FK506 3h | 17.34 | 22.43 | -5.086 | 3.553 | 9 | 9 | 1.431 | 76.00 |
| FK506 6h | 22.87 | 27.93 | -5.060 | 3.388 | 11 | 9 | 1.494 | 76.00 |
| FK506 12h | 24.47 | 32.53 | -8.066 | 3.901 | 7 | 8 | 2.068 | 76.00 |
| FK506 24h | 25.63 | 34.22 | -8.594 | 4.193 | 7 | 6 | 2.049 | 76.00 |

| Tukey's multiple comparisons test | Predicted (LS) mean diff. | 95.00% CI of diff. | Below threshold? | Summary | Adjusted P Value |
|---|---|---|---|---|---|
| wt | | | | | |
| FK506 0h vs. FK506 3h | -1.592 | -10.88 to 7.696 | No | ns | 0.9891 |
| FK506 0h vs. FK506 6h | -7.118 | -15.91 to 1.674 | No | ns | 0.1686 |
| FK506 0h vs. FK506 12h | -8.718 | -18.73 to 1.299 | No | ns | 0.1179 |
| FK506 0h vs. FK506 24h | -9.881 | -19.90 to 0.1361 | No | ns | 0.0550 |
| FK506 3h vs. FK506 6h | -5.526 | -14.99 to 3.940 | No | ns | 0.4824 |
| FK506 3h vs. FK506 12h | -7.126 | -17.74 to 3.487 | No | ns | 0.3389 |
| FK506 3h vs. FK506 24h | -8.289 | -18.90 to 2.325 | No | ns | 0.1976 |
| FK506 6h vs. FK506 12h | -1.600 | -11.78 to 8.583 | No | ns | 0.9921 |
| FK506 6h vs. FK506 24h | -2.763 | -12.95 to 7.420 | No | ns | 0.9417 |
| FK506 12h vs. FK506 24h | -1.163 | -12.42 to 10.10 | No | ns | 0.9985 |
| ts | | | | | |
| FK506 0h vs. FK506 3h | -5.508 | -15.74 to 4.726 | No | ns | 0.5631 |
| FK506 0h vs. FK506 6h | -11.01 | -21.24 to -0.7748 | Yes | * | 0.0287 |
| FK506 0h vs. FK506 12h | -15.61 | -26.15 to -5.084 | Yes | *** | 0.0008 |
| FK506 0h vs. FK506 24h | -17.31 | -28.68 to -5.931 | Yes | *** | 0.0006 |
| FK506 3h vs. FK506 6h | -5.501 | -15.43 to 4.428 | No | ns | 0.5349 |
| FK506 3h vs. FK506 12h | -10.11 | -20.34 to 0.1274 | No | ns | 0.0546 |
| FK506 3h vs. FK506 24h | -11.80 | -22.90 to -0.6970 | Yes | * | 0.0316 |
| FK506 6h vs. FK506 12h | -4.606 | -14.84 to 5.628 | No | ns | 0.7176 |
| FK506 6h vs. FK506 24h | -6.297 | -17.40 to 4.804 | No | ns | 0.5115 |
| FK506 12h vs. FK506 24h | -1.691 | -13.07 to 9.684 | No | ns | 0.9936 |

| Test details | Predicted (LS) mean 1 | Predicted (LS) mean 2 | Predicted (LS) mean diff. | SE of diff. | N1 | N2 | q | DF |
|---|---|---|---|---|---|---|---|---|
| wt | | | | | | | | |
| FK506 0h vs. FK506 3h | 15.75 | 17.34 | -1.592 | 3.324 | 12 | 9 | 0.6772 | 76.00 |
| FK506 0h vs. FK506 6h | 15.75 | 22.87 | -7.118 | 3.146 | 12 | 11 | 3.199 | 76.00 |
| FK506 0h vs. FK506 12h | 15.75 | 24.47 | -8.718 | 3.585 | 12 | 7 | 3.439 | 76.00 |
| FK506 0h vs. FK506 24h | 15.75 | 25.63 | -9.881 | 3.585 | 12 | 7 | 3.898 | 76.00 |
| FK506 3h vs. FK506 6h | 17.34 | 22.87 | -5.526 | 3.388 | 9 | 11 | 2.307 | 76.00 |
| FK506 3h vs. FK506 12h | 17.34 | 24.47 | -7.126 | 3.798 | 9 | 7 | 2.653 | 76.00 |
| FK506 3h vs. FK506 24h | 17.34 | 25.63 | -8.289 | 3.798 | 9 | 7 | 3.086 | 76.00 |
| FK506 6h vs. FK506 12h | 22.87 | 24.47 | -1.600 | 3.644 | 11 | 7 | 0.6210 | 76.00 |
| FK506 6h vs. FK506 24h | 22.87 | 25.63 | -2.763 | 3.644 | 11 | 7 | 1.072 | 76.00 |
| FK506 12h vs. FK506 24h | 24.47 | 25.63 | -1.163 | 4.029 | 7 | 7 | 0.4081 | 76.00 |
| ts | | | | | | | | |
| FK506 0h vs. FK506 3h | 16.92 | 22.43 | -5.508 | 3.662 | 8 | 9 | 2.127 | 76.00 |
| FK506 0h vs. FK506 6h | 16.92 | 27.93 | -11.01 | 3.662 | 8 | 9 | 4.251 | 76.00 |
| FK506 0h vs. FK506 12h | 16.92 | 32.53 | -15.61 | 3.769 | 8 | 8 | 5.860 | 76.00 |
| FK506 0h vs. FK506 24h | 16.92 | 34.22 | -17.31 | 4.070 | 8 | 6 | 6.012 | 76.00 |
| FK506 3h vs. FK506 6h | 22.43 | 27.93 | -5.501 | 3.553 | 9 | 9 | 2.189 | 76.00 |
| FK506 3h vs. FK506 12h | 22.43 | 32.53 | -10.11 | 3.662 | 9 | 8 | 3.903 | 76.00 |
| FK506 3h vs. FK506 24h | 22.43 | 34.22 | -11.80 | 3.972 | 9 | 6 | 4.200 | 76.00 |
| FK506 6h vs. FK506 12h | 27.93 | 32.53 | -4.606 | 3.662 | 9 | 8 | 1.779 | 76.00 |
| FK506 6h vs. FK506 24h | 27.93 | 34.22 | -6.297 | 3.972 | 9 | 6 | 2.242 | 76.00 |
| FK506 12h vs. FK506 24h | 32.53 | 34.22 | -1.691 | 4.070 | 8 | 6 | 0.5874 | 76.00 |

**Table S6A: Statistics for Figure 4**

### Decay

| Source of Variation | % of total variation | P value | P value summary | Significant? | |
|---|---|---|---|---|---|
| Interaction | 2.273 | 0.6411 | ns | No | |
| FK506 | 21.43 | 0.0003 | *** | Yes | |
| Genotype | 8.782 | 0.0025 | ** | Yes | |
| **ANOVA table** | **SS (Type III)** | **DF** | **MS** | **F (DFn, DFd)** | **P value** |
| Interaction | 1.733 | 4 | 0.4332 | F (4, 75) = 0.6321 | P=0.6411 |
| FK506 | 16.34 | 4 | 4.085 | F (4, 75) = 5.961 | P=0.0003 |
| Genotype | 6.695 | 1 | 6.695 | F (1, 75) = 9.769 | P=0.0025 |
| Residual | 51.40 | 75 | 0.6853 | | |
| **Difference between column means** | | | | | |
| Predicted (LS) mean of wt | 4.903 | | | | |
| Predicted (LS) mean of ts | 4.332 | | | | |
| Difference between predicted means | 0.5710 | | | | |
| SE of difference | 0.1827 | | | | |
| 95% CI of difference | 0.2071 to 0.9350 | | | | |
| **Data summary** | | | | | |
| Number of columns (Genotype) | 2 | | | | |
| Number of rows (FK506) | 5 | | | | |
| Number of values | 85 | | | | |

| Tukey's multiple comparisor | Predicted (LS) mean diff. | 95.00% CI of diff. | Below threshold? | Summary | Adjusted P Value | Test details | Predicted (LS) mean 1 | Predicted (LS) mea | Predicted (LS) mea | SE of diff. | N1 | N2 | q | DF |
|---|---|---|---|---|---|---|---|---|---|---|---|---|---|---|
| wt | | | | | | wt | | | | | | | | |
| FK506 0h vs. FK506 3h | -0.007325 | -1.047 to 1.033 | No | ns | >0.9999 | FK506 0h vs. FK506 3h | 5.290 | 5.297 | -0.007325 | 0.3721 | 11 | 9 | 0.02784 | 75.00 |
| FK506 0h vs. FK506 6h | 0.5043 | -0.4824 to 1.491 | No | ns | 0.6113 | FK506 0h vs. FK506 6h | 5.290 | 4.785 | 0.5043 | 0.3530 | 11 | 11 | 2.021 | 75.00 |
| FK506 0h vs. FK506 12h | 0.5843 | -0.5345 to 1.703 | No | ns | 0.5914 | FK506 0h vs. FK506 12h | 5.290 | 4.705 | 0.5843 | 0.4003 | 11 | 7 | 2.065 | 75.00 |
| FK506 0h vs. FK506 24h | 0.8509 | -0.2679 to 1.970 | No | ns | 0.2201 | FK506 0h vs. FK506 24h | 5.290 | 4.439 | 0.8509 | 0.4003 | 11 | 7 | 3.007 | 75.00 |
| FK506 3h vs. FK506 6h | 0.5116 | -0.5284 to 1.552 | No | ns | 0.6453 | FK506 3h vs. FK506 6h | 5.297 | 4.785 | 0.5116 | 0.3721 | 9 | 11 | 1.945 | 75.00 |
| FK506 3h vs. FK506 12h | 0.5916 | -0.5745 to 1.758 | No | ns | 0.6181 | FK506 3h vs. FK506 12h | 5.297 | 4.705 | 0.5916 | 0.4172 | 9 | 7 | 2.006 | 75.00 |
| FK506 3h vs. FK506 24h | 0.8583 | -0.3079 to 2.024 | No | ns | 0.2496 | FK506 3h vs. FK506 24h | 5.297 | 4.439 | 0.8583 | 0.4172 | 9 | 7 | 2.909 | 75.00 |
| FK506 6h vs. FK506 12h | 0.07999 | -1.039 to 1.199 | No | ns | 0.9996 | FK506 6h vs. FK506 12h | 4.785 | 4.705 | 0.07999 | 0.4003 | 11 | 7 | 0.2826 | 75.00 |
| FK506 6h vs. FK506 24h | 0.3466 | -0.7722 to 1.465 | No | ns | 0.9084 | FK506 6h vs. FK506 24h | 4.785 | 4.439 | 0.3466 | 0.4003 | 11 | 7 | 1.225 | 75.00 |
| FK506 12h vs. FK506 24h | 0.2666 | -0.9703 to 1.504 | No | ns | 0.9743 | FK506 12h vs. FK506 24h | 4.705 | 4.439 | 0.2666 | 0.4425 | 7 | 7 | 0.8521 | 75.00 |
| ts | | | | | | ts | | | | | | | | |
| FK506 0h vs. FK506 3h | 0.06465 | -1.060 to 1.189 | No | ns | 0.9998 | FK506 0h vs. FK506 3h | 5.006 | 4.942 | 0.06465 | 0.4023 | 8 | 9 | 0.2273 | 75.00 |
| FK506 0h vs. FK506 6h | 0.6410 | -0.4835 to 1.765 | No | ns | 0.5063 | FK506 0h vs. FK506 6h | 5.006 | 4.365 | 0.6410 | 0.4023 | 8 | 9 | 2.253 | 75.00 |
| FK506 0h vs. FK506 12h | 1.386 | 0.2286 to 2.543 | Yes | * | 0.0109 | FK506 0h vs. FK506 12h | 5.006 | 3.621 | 1.386 | 0.4139 | 8 | 8 | 4.734 | 75.00 |
| FK506 0h vs. FK506 24h | 1.280 | 0.02991 to 2.529 | Yes | * | 0.0421 | FK506 0h vs. FK506 24h | 5.006 | 3.727 | 1.280 | 0.4471 | 8 | 6 | 4.048 | 75.00 |
| FK506 3h vs. FK506 6h | 0.5763 | -0.5145 to 1.667 | No | ns | 0.5806 | FK506 3h vs. FK506 6h | 4.942 | 4.365 | 0.5763 | 0.3902 | 9 | 9 | 2.088 | 75.00 |
| FK506 3h vs. FK506 12h | 1.321 | 0.1965 to 2.445 | Yes | * | 0.0131 | FK506 3h vs. FK506 12h | 4.942 | 3.621 | 1.321 | 0.4023 | 9 | 8 | 4.644 | 75.00 |
| FK506 3h vs. FK506 24h | 1.215 | -0.004619 to 2.435 | No | ns | 0.0514 | FK506 3h vs. FK506 24h | 4.942 | 3.727 | 1.215 | 0.4363 | 9 | 6 | 3.938 | 75.00 |
| FK506 6h vs. FK506 12h | 0.7447 | -0.3798 to 1.869 | No | ns | 0.3526 | FK506 6h vs. FK506 12h | 4.365 | 3.621 | 0.7447 | 0.4023 | 9 | 8 | 2.618 | 75.00 |
| FK506 6h vs. FK506 24h | 0.6387 | -0.5809 to 1.858 | No | ns | 0.5889 | FK506 6h vs. FK506 24h | 4.365 | 3.727 | 0.6387 | 0.4363 | 9 | 6 | 2.070 | 75.00 |
| FK506 12h vs. FK506 24h | -0.1060 | -1.356 to 1.144 | No | ns | 0.9993 | FK506 12h vs. FK506 24h | 3.621 | 3.727 | -0.1060 | 0.4471 | 8 | 6 | 0.3352 | 75.00 |

#### Table S7A: Statistics for Figure 5 - pCaMKII synaptic

| Source of Variation | % of total variation | P value | P value | Significant? | |
|---|---|---|---|---|---|
| Interaction | 2.541 | <0.0001 | **** | Yes | |
| Hours TTX | 54.98 | <0.0001 | **** | Yes | |
| Genotype | 2.403 | <0.0001 | **** | Yes | |
| **ANOVA table** | **SS (Type III)** | **DF** | **MS** | **F (DFn, DFd)** | **P value** |
| Interaction | 2545 | 6 | 424.2 | F (6, 690) = 7.905 | P<0.0001 |
| Hours TTX | 55085 | 6 | 9181 | F (6, 690) = 171.1 | P<0.0001 |
| Genotype | 2408 | 1 | 2408 | F (1, 690) = 44.86 | P<0.0001 |
| Residual | 37032 | 690 | 53.67 | | |
| **Difference between column means** | | | | | |
| Predicted (LS) mean of wt | 14.53 | | | | |
| Predicted (LS) mean of ts | 18.30 | | | | |
| Difference between predicted means | -3.773 | | | | |
| SE of difference | 0.5633 | | | | |
| 95% CI of difference | -4.879 to -2.667 | | | | |
| **Data summary** | | | | | |
| Number of columns (Genotype) | 2 | | | | |
| Number of rows (Hours TTX) | 7 | | | | |
| Number of values | 704 | | | | |

| Šidák's multiple comparisons test | Predicted (LS) mean diff. | 95.00% CI of diff. | Below threshold? | Summary | Adjusted P | | | |
|---|---|---|---|---|---|---|---|---|
| wt - ts | | | | | | | | |
| ttx00 | 0.8417 | -2.908 to 4.591 | No | ns | 0.9960 | | | |
| ttx03 | -1.142 | -4.568 to 2.285 | No | ns | 0.9607 | | | |
| ttx06 | -10.72 | -14.69 to -6.749 | Yes | **** | <0.0001 | | | |
| ttx12 | -3.761 | -8.331 to 0.8083 | No | ns | 0.1749 | | | |
| ttx24 | -7.798 | -11.84 to -3.753 | Yes | **** | <0.0001 | | | |
| ttx48 | -1.715 | -5.850 to 2.420 | No | ns | 0.8840 | | | |
| ttx72 | -2.117 | -6.201 to 1.966 | No | ns | 0.7132 | | | |
| **Test details** | **Predicted (LS) mean 1** | **Predicted (LS) mean 2** | **Predicted (LS) mean diff.** | **SE of diff.** | **N1** | **N2** | **t** | **DF** |
| wt - ts | | | | | | | | |
| ttx00 | 12.18 | 11.34 | 0.8417 | 1.393 | 59 | 52 | 0.6041 | 690.0 |
| ttx03 | 5.908 | 7.050 | -1.142 | 1.273 | 71 | 62 | 0.8966 | 690.0 |
| ttx06 | 30.69 | 41.41 | -10.72 | 1.476 | 44 | 56 | 7.264 | 690.0 |
| ttx12 | 14.41 | 18.17 | -3.761 | 1.698 | 49 | 30 | 2.215 | 690.0 |
| ttx24 | 9.352 | 17.15 | -7.798 | 1.503 | 47 | 48 | 5.187 | 690.0 |
| ttx48 | 12.66 | 14.38 | -1.715 | 1.537 | 47 | 44 | 1.116 | 690.0 |
| ttx72 | 16.48 | 18.60 | -2.117 | 1.518 | 54 | 41 | 1.395 | 690.0 |

| Tukey's multiple comparisons test | Predicted (LS) mea | 95.00% CI of diff. | Below threshold? | Summary | Adjusted P Value | Predicted (LS) mea | Predicted (LS) mea | Predicted (LS) mea | SE of diff. | N1 | N2 | q | DF |
|---|---|---|---|---|---|---|---|---|---|---|---|---|---|
| wt | | | | | | | | | | | | | |
| ttx00 vs. ttx03 | 6.274 | 2.458 to 10.09 | Yes | **** | <0.0001 | 12.18 | 5.908 | 6.274 | 1.291 | 59 | 71 | 6.875 | 690.0 |
| ttx00 vs. ttx06 | -18.50 | -22.82 to -14.19 | Yes | **** | <0.0001 | 12.18 | 30.69 | -18.50 | 1.459 | 59 | 44 | 17.93 | 690.0 |
| ttx00 vs. ttx12 | -2.224 | -6.412 to 1.963 | No | ns | 0.7010 | 12.18 | 14.41 | -2.224 | 1.416 | 59 | 49 | 2.222 | 690.0 |
| ttx00 vs. ttx24 | 2.831 | -1.405 to 7.066 | No | ns | 0.4308 | 12.18 | 9.352 | 2.831 | 1.432 | 59 | 47 | 2.795 | 690.0 |
| ttx00 vs. ttx48 | -0.4797 | -4.715 to 3.756 | No | ns | 0.9999 | 12.18 | 12.66 | -0.4797 | 1.432 | 59 | 47 | 0.4736 | 690.0 |
| ttx00 vs. ttx72 | -4.296 | -8.376 to -0.2165 | Yes | * | 0.0315 | 12.18 | 16.48 | -4.296 | 1.380 | 59 | 54 | 4.404 | 690.0 |
| ttx03 vs. ttx06 | -24.78 | -28.93 to -20.62 | Yes | **** | <0.0001 | 5.908 | 30.69 | -24.78 | 1.406 | 71 | 44 | 24.93 | 690.0 |
| ttx03 vs. ttx12 | -8.499 | -12.52 to -4.475 | Yes | **** | <0.0001 | 5.908 | 14.41 | -8.499 | 1.361 | 71 | 49 | 8.834 | 690.0 |
| ttx03 vs. ttx24 | -3.444 | -7.517 to 0.6300 | No | ns | 0.1609 | 5.908 | 9.352 | -3.444 | 1.378 | 71 | 47 | 3.535 | 690.0 |
| ttx03 vs. ttx48 | -6.754 | -10.83 to -2.680 | Yes | **** | <0.0001 | 5.908 | 12.66 | -6.754 | 1.378 | 71 | 47 | 6.933 | 690.0 |
| ttx03 vs. ttx72 | -10.57 | -14.48 to -6.659 | Yes | **** | <0.0001 | 5.908 | 16.48 | -10.57 | 1.323 | 71 | 54 | 11.30 | 690.0 |
| ttx06 vs. ttx12 | 16.28 | 11.78 to 20.78 | Yes | **** | <0.0001 | 30.69 | 14.41 | 16.28 | 1.522 | 44 | 49 | 15.13 | 690.0 |
| ttx06 vs. ttx24 | 21.33 | 16.79 to 25.88 | Yes | **** | <0.0001 | 30.69 | 9.352 | 21.33 | 1.537 | 44 | 47 | 19.63 | 690.0 |
| ttx06 vs. ttx48 | 18.02 | 13.48 to 22.57 | Yes | **** | <0.0001 | 30.69 | 12.66 | 18.02 | 1.537 | 44 | 47 | 16.59 | 690.0 |
| ttx06 vs. ttx72 | 14.21 | 9.807 to 18.61 | Yes | **** | <0.0001 | 30.69 | 16.48 | 14.21 | 1.488 | 44 | 54 | 13.50 | 690.0 |
| ttx12 vs. ttx24 | 5.055 | 0.6322 to 9.478 | Yes | * | 0.0135 | 14.41 | 9.352 | 5.055 | 1.496 | 49 | 47 | 4.780 | 690.0 |
| ttx12 vs. ttx48 | 1.745 | -2.678 to 6.168 | No | ns | 0.9065 | 14.41 | 12.66 | 1.745 | 1.496 | 49 | 47 | 1.650 | 690.0 |
| ttx12 vs. ttx72 | -2.072 | -6.346 to 2.202 | No | ns | 0.7836 | 14.41 | 16.48 | -2.072 | 1.445 | 49 | 54 | 2.027 | 690.0 |
| ttx24 vs. ttx48 | -3.310 | -7.779 to 1.158 | No | ns | 0.3019 | 9.352 | 12.66 | -3.310 | 1.511 | 47 | 47 | 3.098 | 690.0 |
| ttx24 vs. ttx72 | -7.127 | -11.45 to -2.805 | Yes | **** | <0.0001 | 9.352 | 16.48 | -7.127 | 1.461 | 47 | 54 | 6.897 | 690.0 |
| ttx48 vs. ttx72 | -3.817 | -8.138 to 0.5050 | No | ns | 0.1240 | 12.66 | 16.48 | -3.817 | 1.461 | 47 | 54 | 3.693 | 690.0 |
| ts | | | | | | | | | | | | | |
| ttx00 vs. ttx03 | 4.291 | 0.2172 to 8.364 | Yes | * | 0.0314 | 11.34 | 7.050 | 4.291 | 1.378 | 52 | 62 | 4.405 | 690.0 |
| ttx00 vs. ttx06 | -30.06 | -34.24 to -25.89 | Yes | **** | <0.0001 | 11.34 | 41.41 | -30.06 | 1.411 | 52 | 56 | 30.14 | 690.0 |
| ttx00 vs. ttx12 | -6.828 | -11.79 to -1.861 | Yes | ** | 0.0010 | 11.34 | 18.17 | -6.828 | 1.680 | 52 | 30 | 5.749 | 690.0 |
| ttx00 vs. ttx24 | -5.809 | -10.15 to -1.473 | Yes | ** | 0.0016 | 11.34 | 17.15 | -5.809 | 1.466 | 52 | 48 | 5.603 | 690.0 |
| ttx00 vs. ttx48 | -3.036 | -7.474 to 1.401 | No | ns | 0.4007 | 11.34 | 14.38 | -3.036 | 1.501 | 52 | 44 | 2.861 | 690.0 |
| ttx00 vs. ttx72 | -7.255 | -11.78 to -2.731 | Yes | **** | <0.0001 | 11.34 | 18.60 | -7.255 | 1.530 | 52 | 41 | 6.706 | 690.0 |
| ttx03 vs. ttx06 | -34.36 | -38.35 to -30.36 | Yes | **** | <0.0001 | 7.050 | 41.41 | -34.36 | 1.351 | 62 | 56 | 35.97 | 690.0 |
| ttx03 vs. ttx12 | -11.12 | -15.94 to -6.301 | Yes | **** | <0.0001 | 7.050 | 18.17 | -11.12 | 1.629 | 62 | 30 | 9.651 | 690.0 |
| ttx03 vs. ttx24 | -10.10 | -14.26 to -5.935 | Yes | **** | <0.0001 | 7.050 | 17.15 | -10.10 | 1.408 | 62 | 48 | 10.14 | 690.0 |
| ttx03 vs. ttx48 | -7.327 | -11.60 to -3.057 | Yes | **** | <0.0001 | 7.050 | 14.38 | -7.327 | 1.444 | 62 | 44 | 7.175 | 690.0 |
| ttx03 vs. ttx72 | -11.55 | -15.91 to -7.185 | Yes | **** | <0.0001 | 7.050 | 18.60 | -11.55 | 1.475 | 62 | 41 | 11.07 | 690.0 |
| ttx06 vs. ttx12 | 23.24 | 18.34 to 28.14 | Yes | **** | <0.0001 | 41.41 | 18.17 | 23.24 | 1.658 | 56 | 30 | 19.83 | 690.0 |
| ttx06 vs. ttx24 | 24.26 | 19.99 to 28.52 | Yes | **** | <0.0001 | 41.41 | 17.15 | 24.26 | 1.441 | 56 | 48 | 23.80 | 690.0 |
| ttx06 vs. ttx48 | 27.03 | 22.66 to 31.39 | Yes | **** | <0.0001 | 41.41 | 14.38 | 27.03 | 1.476 | 56 | 44 | 25.90 | 690.0 |
| ttx06 vs. ttx72 | 22.81 | 18.36 to 27.26 | Yes | **** | <0.0001 | 41.41 | 18.60 | 22.81 | 1.506 | 56 | 41 | 21.42 | 690.0 |
| ttx12 vs. ttx24 | 1.019 | -4.023 to 6.060 | No | ns | 0.9969 | 18.17 | 17.15 | 1.019 | 1.705 | 30 | 48 | 0.8449 | 690.0 |
| ttx12 vs. ttx48 | 3.791 | -1.338 to 8.921 | No | ns | 0.3045 | 18.17 | 14.38 | 3.791 | 1.735 | 30 | 44 | 3.091 | 690.0 |
| ttx12 vs. ttx72 | -0.4276 | -5.632 to 4.777 | No | ns | >0.9999 | 18.17 | 18.60 | -0.4276 | 1.760 | 30 | 41 | 0.3436 | 690.0 |
| ttx24 vs. ttx48 | 2.773 | -1.749 to 7.294 | No | ns | 0.5393 | 17.15 | 14.38 | 2.773 | 1.529 | 48 | 44 | 2.565 | 690.0 |
| ttx24 vs. ttx72 | -1.446 | -6.053 to 3.161 | No | ns | 0.9680 | 17.15 | 18.60 | -1.446 | 1.558 | 48 | 41 | 1.313 | 690.0 |
| ttx48 vs. ttx72 | -4.219 | -8.921 to 0.4833 | No | ns | 0.1121 | 14.38 | 18.60 | -4.219 | 1.590 | 44 | 41 | 3.752 | 690.0 |

#### Table S7B: Statistics for Figure 5 - pCaMKII dendritic shaft

| Source of Variation | % of total variation | P value | P value su | Significant? | |
|---|---|---|---|---|---|
| Interaction | 1.162 | 0.0013 | ** | Yes | |
| Hours TTX | 58.55 | <0.0001 | **** | Yes | |
| Genotype | 1.404 | <0.0001 | **** | Yes | |
| **ANOVA table** | **SS (Type III)** | **DF** | **MS** | **F (DFn, DFd)** | **P value** |
| Interaction | 586.3 | 6 | 97.72 | F (6, 690) = 3.688 | P=0.0013 |
| Hours TTX | 29551 | 6 | 4925 | F (6, 690) = 185.9 | P<0.0001 |
| Genotype | 708.5 | 1 | 708.5 | F (1, 690) = 26.73 | P<0.0001 |
| Residual | 18286 | 690 | 26.50 | | |
| **Difference between column means** | | | | | |
| Predicted (LS) mean of wt | 11.25 | | | | |
| Predicted (LS) mean of ts | 13.30 | | | | |
| Difference between predicted means | -2.047 | | | | |
| SE of difference | 0.3959 | | | | |
| 95% CI of difference | -2.824 to -1.270 | | | | |
| **Data summary** | | | | | |
| Number of columns (Genotype) | 2 | | | | |
| Number of rows (Hours TTX) | 7 | | | | |
| Number of values | 704 | | | | |

| Šidák's multiple comparisons test | Predicted (LS) mean diff. | 95.00% CI of diff. | Below thre | Summary | Adjusted F | | | |
|---|---|---|---|---|---|---|---|---|
| wt - ts | | | | | | | | |
| ttx00 | -1.223 | -3.857 to 1.412 | No | ns | 0.8117 | | | |
| ttx03 | -1.060 | -3.468 to 1.348 | No | ns | 0.8489 | | | |
| ttx06 | -4.793 | -7.584 to -2.003 | Yes | **** | <0.0001 | | | |
| ttx12 | 0.2110 | -3.000 to 3.422 | No | ns | >0.9999 | | | |
| ttx24 | -4.792 | -7.634 to -1.949 | Yes | **** | <0.0001 | | | |
| ttx48 | -0.2591 | -3.165 to 2.647 | No | ns | >0.9999 | | | |
| ttx72 | -2.411 | -5.281 to 0.4579 | No | ns | 0.1566 | | | |
| **Test details** | **Predicted (LS) mean 1** | **Predicted (LS) mea** | **Predicted** | **SE of diff.** | **N1** | **N2** | **t** | **DF** |
| wt - ts | | | | | | | | |
| ttx00 | 7.716 | 8.939 | -1.223 | 0.9792 | 59 | 52 | 1.249 | 690.0 |
| ttx03 | 5.296 | 6.356 | -1.060 | 0.8948 | 71 | 62 | 1.185 | 690.0 |
| ttx06 | 24.70 | 29.49 | -4.793 | 1.037 | 44 | 56 | 4.622 | 690.0 |
| ttx12 | 11.70 | 11.48 | 0.2110 | 1.193 | 49 | 30 | 0.1768 | 690.0 |
| ttx24 | 7.953 | 12.74 | -4.792 | 1.056 | 47 | 48 | 4.536 | 690.0 |
| ttx48 | 10.17 | 10.43 | -0.2591 | 1.080 | 47 | 44 | 0.2399 | 690.0 |
| ttx72 | 11.24 | 13.65 | -2.411 | 1.066 | 54 | 41 | 2.261 | 690.0 |

| Tukey's multiple cc | Predicted ( | 95.00% CI of diff. | Below thre | Summary | Adjusted F |
|---|---|---|---|---|---|
| wt | | | | | |
| ttx00 vs. ttx03 | 2.420 | -0.2618 to 5.102 | No | ns | 0.1080 |
| ttx00 vs. ttx06 | -16.98 | -20.02 to -13.95 | Yes | **** | <0.0001 |
| ttx00 vs. ttx12 | -3.979 | -6.921 to -1.037 | Yes | ** | 0.0014 |
| ttx00 vs. ttx24 | -0.2372 | -3.213 to 2.739 | No | ns | >0.9999 |
| ttx00 vs. ttx48 | -2.454 | -5.431 to 0.5219 | No | ns | 0.1842 |
| ttx00 vs. ttx72 | -3.522 | -6.389 to -0.6550 | Yes | ** | 0.0056 |
| ttx03 vs. ttx06 | -19.40 | -22.32 to -16.48 | Yes | **** | <0.0001 |
| ttx03 vs. ttx12 | -6.399 | -9.226 to -3.572 | Yes | **** | <0.0001 |
| ttx03 vs. ttx24 | -2.657 | -5.520 to 0.2055 | No | ns | 0.0890 |
| ttx03 vs. ttx48 | -4.874 | -7.737 to -2.012 | Yes | **** | <0.0001 |
| ttx03 vs. ttx72 | -5.942 | -8.690 to -3.193 | Yes | **** | <0.0001 |
| ttx06 vs. ttx12 | 13.00 | 9.843 to 16.17 | Yes | **** | <0.0001 |
| ttx06 vs. ttx24 | 16.75 | 13.55 to 19.94 | Yes | **** | <0.0001 |
| ttx06 vs. ttx48 | 14.53 | 11.34 to 17.72 | Yes | **** | <0.0001 |
| ttx06 vs. ttx72 | 13.46 | 10.37 to 16.55 | Yes | **** | <0.0001 |
| ttx12 vs. ttx24 | 3.742 | 0.6340 to 6.850 | Yes | ** | 0.0072 |
| ttx12 vs. ttx48 | 1.525 | -1.583 to 4.633 | No | ns | 0.7737 |
| ttx12 vs. ttx72 | 0.4573 | -2.546 to 3.461 | No | ns | 0.9994 |
| ttx24 vs. ttx48 | -2.217 | -5.357 to 0.9231 | No | ns | 0.3610 |
| ttx24 vs. ttx72 | -3.285 | -6.321 to -0.2480 | Yes | * | 0.0242 |
| ttx48 vs. ttx72 | -1.068 | -4.104 to 1.969 | No | ns | 0.9447 |
| ts | | | | | |
| ttx00 vs. ttx03 | 2.583 | -0.2798 to 5.445 | No | ns | 0.1080 |
| ttx00 vs. ttx06 | -20.55 | -23.49 to -17.62 | Yes | **** | <0.0001 |
| ttx00 vs. ttx12 | -2.545 | -6.036 to 0.9446 | No | ns | 0.3208 |
| ttx00 vs. ttx24 | -3.806 | -6.853 to -0.7592 | Yes | ** | 0.0044 |
| ttx00 vs. ttx48 | -1.491 | -4.609 to 1.628 | No | ns | 0.7945 |
| ttx00 vs. ttx72 | -4.711 | -7.890 to -1.531 | Yes | *** | 0.0003 |
| ttx03 vs. ttx06 | -23.14 | -25.94 to -20.33 | Yes | **** | <0.0001 |
| ttx03 vs. ttx12 | -5.128 | -8.514 to -1.743 | Yes | *** | 0.0002 |
| ttx03 vs. ttx24 | -6.389 | -9.316 to -3.462 | Yes | **** | <0.0001 |
| ttx03 vs. ttx48 | -4.073 | -7.074 to -1.073 | Yes | ** | 0.0013 |
| ttx03 vs. ttx72 | -7.293 | -10.36 to -4.229 | Yes | **** | <0.0001 |
| ttx06 vs. ttx12 | 18.01 | 14.57 to 21.45 | Yes | **** | <0.0001 |
| ttx06 vs. ttx24 | 16.75 | 13.75 to 19.74 | Yes | **** | <0.0001 |
| ttx06 vs. ttx48 | 19.06 | 16.00 to 22.13 | Yes | **** | <0.0001 |
| ttx06 vs. ttx72 | 15.84 | 12.72 to 18.97 | Yes | **** | <0.0001 |
| ttx12 vs. ttx24 | -1.261 | -4.804 to 2.282 | No | ns | 0.9414 |
| ttx12 vs. ttx48 | 1.055 | -2.549 to 4.659 | No | ns | 0.9775 |
| ttx12 vs. ttx72 | -2.165 | -5.823 to 1.492 | No | ns | 0.5821 |
| ttx24 vs. ttx48 | 2.316 | -0.8616 to 5.493 | No | ns | 0.3217 |
| ttx24 vs. ttx72 | -0.9045 | -4.142 to 2.333 | No | ns | 0.9822 |
| ttx48 vs. ttx72 | -3.220 | -6.524 to 0.08430 | No | ns | 0.0618 |

| Test details | Predicted ( | Predicted (LS) mea | Predicted | SE of diff. | N1 | N2 | q | DF |
|---|---|---|---|---|---|---|---|---|
| wt | | | | | | | | |
| ttx00 vs. ttx03 | 7.716 | 5.296 | 2.420 | 0.9069 | 59 | 71 | 3.774 | 690.0 |
| ttx00 vs. ttx06 | 7.716 | 24.70 | -16.98 | 1.025 | 59 | 44 | 23.42 | 690.0 |
| ttx00 vs. ttx12 | 7.716 | 11.70 | -3.979 | 0.9950 | 59 | 49 | 5.656 | 690.0 |
| ttx00 vs. ttx24 | 7.716 | 7.953 | -0.2372 | 1.006 | 59 | 47 | 0.3333 | 690.0 |
| ttx00 vs. ttx48 | 7.716 | 10.17 | -2.454 | 1.006 | 59 | 47 | 3.449 | 690.0 |
| ttx00 vs. ttx72 | 7.716 | 11.24 | -3.522 | 0.9695 | 59 | 54 | 5.137 | 690.0 |
| ttx03 vs. ttx06 | 5.296 | 24.70 | -19.40 | 0.9877 | 71 | 44 | 27.78 | 690.0 |
| ttx03 vs. ttx12 | 5.296 | 11.70 | -6.399 | 0.9561 | 71 | 49 | 9.465 | 690.0 |
| ttx03 vs. ttx24 | 5.296 | 7.953 | -2.657 | 0.9680 | 71 | 47 | 3.882 | 690.0 |
| ttx03 vs. ttx48 | 5.296 | 10.17 | -4.874 | 0.9680 | 71 | 47 | 7.121 | 690.0 |
| ttx03 vs. ttx72 | 5.296 | 11.24 | -5.942 | 0.9295 | 71 | 54 | 9.040 | 690.0 |
| ttx06 vs. ttx12 | 24.70 | 11.70 | 13.00 | 1.069 | 44 | 49 | 17.20 | 690.0 |
| ttx06 vs. ttx24 | 24.70 | 7.953 | 16.75 | 1.080 | 44 | 47 | 21.93 | 690.0 |
| ttx06 vs. ttx48 | 24.70 | 10.17 | 14.53 | 1.080 | 44 | 47 | 19.03 | 690.0 |
| ttx06 vs. ttx72 | 24.70 | 11.24 | 13.46 | 1.045 | 44 | 54 | 18.21 | 690.0 |
| ttx12 vs. ttx24 | 11.70 | 7.953 | 3.742 | 1.051 | 49 | 47 | 5.035 | 690.0 |
| ttx12 vs. ttx48 | 11.70 | 10.17 | 1.525 | 1.051 | 49 | 47 | 2.052 | 690.0 |
| ttx12 vs. ttx72 | 11.70 | 11.24 | 0.4573 | 1.016 | 49 | 54 | 0.6367 | 690.0 |
| ttx24 vs. ttx48 | 7.953 | 10.17 | -2.217 | 1.062 | 47 | 47 | 2.953 | 690.0 |
| ttx24 vs. ttx72 | 7.953 | 11.24 | -3.285 | 1.027 | 47 | 54 | 4.523 | 690.0 |
| ttx48 vs. ttx72 | 10.17 | 11.24 | -1.068 | 1.027 | 47 | 54 | 1.470 | 690.0 |
| ts | | | | | | | | |
| ttx00 vs. ttx03 | 8.939 | 6.356 | 2.583 | 0.9680 | 52 | 62 | 3.773 | 690.0 |
| ttx00 vs. ttx06 | 8.939 | 29.49 | -20.55 | 0.9914 | 52 | 56 | 29.32 | 690.0 |
| ttx00 vs. ttx12 | 8.939 | 11.48 | -2.545 | 1.180 | 52 | 30 | 3.050 | 690.0 |
| ttx00 vs. ttx24 | 8.939 | 12.74 | -3.806 | 1.030 | 52 | 48 | 5.224 | 690.0 |
| ttx00 vs. ttx48 | 8.939 | 10.43 | -1.491 | 1.054 | 52 | 44 | 1.999 | 690.0 |
| ttx00 vs. ttx72 | 8.939 | 13.65 | -4.711 | 1.075 | 52 | 41 | 6.196 | 690.0 |
| ttx03 vs. ttx06 | 6.356 | 29.49 | -23.14 | 0.9490 | 62 | 56 | 34.48 | 690.0 |
| ttx03 vs. ttx12 | 6.356 | 11.48 | -5.128 | 1.145 | 62 | 30 | 6.334 | 690.0 |
| ttx03 vs. ttx24 | 6.356 | 12.74 | -6.389 | 0.9897 | 62 | 48 | 9.129 | 690.0 |
| ttx03 vs. ttx48 | 6.356 | 10.43 | -4.073 | 1.015 | 62 | 44 | 5.677 | 690.0 |
| ttx03 vs. ttx72 | 6.356 | 13.65 | -7.293 | 1.036 | 62 | 41 | 9.954 | 690.0 |
| ttx06 vs. ttx12 | 29.49 | 11.48 | 18.01 | 1.165 | 56 | 30 | 21.87 | 690.0 |
| ttx06 vs. ttx24 | 29.49 | 12.74 | 16.75 | 1.013 | 56 | 48 | 23.39 | 690.0 |
| ttx06 vs. ttx48 | 29.49 | 10.43 | 19.06 | 1.037 | 56 | 44 | 26.00 | 690.0 |
| ttx06 vs. ttx72 | 29.49 | 13.65 | 15.84 | 1.058 | 56 | 41 | 21.18 | 690.0 |
| ttx12 vs. ttx24 | 11.48 | 12.74 | -1.261 | 1.198 | 30 | 48 | 1.488 | 690.0 |
| ttx12 vs. ttx48 | 11.48 | 10.43 | 1.055 | 1.219 | 30 | 44 | 1.224 | 690.0 |
| ttx12 vs. ttx72 | 11.48 | 13.65 | -2.165 | 1.237 | 30 | 41 | 2.476 | 690.0 |
| ttx24 vs. ttx48 | 12.74 | 10.43 | 2.316 | 1.074 | 48 | 44 | 3.048 | 690.0 |
| ttx24 vs. ttx72 | 12.74 | 13.65 | -0.9045 | 1.095 | 48 | 41 | 1.168 | 690.0 |
| ttx48 vs. ttx72 | 10.43 | 13.65 | -3.220 | 1.117 | 44 | 41 | 4.075 | 690.0 |

#### Table S7C: Statistics for Figure 5 - βCaMKII synaptic

| Source of Variation | % of total variation | P value | P value sumn | Significant? | |
|---|---|---|---|---|---|
| Interaction | 5.524 | <0.0001 | **** | Yes | |
| Hours TTX | 11.59 | <0.0001 | **** | Yes | |
| Genotype | 0.4460 | 0.0404 | * | Yes | |
| **ANOVA table** | **SS (Type III)** | **DF** | **MS** | **F (DFn, DFd)** | **P value** |
| Interaction | 1089680138 | 6 | 181613356 | F (6, 761) = 8.703 | P<0.0001 |
| Hours TTX | 2286854152 | 6 | 381142359 | F (6, 761) = 18.27 | P<0.0001 |
| Genotype | 87978242 | 1 | 87978242 | F (1, 761) = 4.216 | P=0.0404 |
| Residual | 15879897695 | 761 | 20867145 | | |
| **Difference between column means** | | | | | |
| Predicted (LS) mean of wt | 11303 | | | | |
| Predicted (LS) mean of ts | 11993 | | | | |
| Difference between predicted means | -689.9 | | | | |
| SE of difference | 336.0 | | | | |
| 95% CI of difference | -1349 to -30.32 | | | | |
| **Data summary** | | | | | |
| Number of columns (Genotype) | 2 | | | | |
| Number of rows (Hours TTX) | 7 | | | | |
| Number of values | 775 | | | | |

| Šidák's multiple c | Predicted (LS | 95.00% CI of diff. | Below thresl | Summary | Adjusted F | | | |
|---|---|---|---|---|---|---|---|---|
| wt - ts | | | | | | | | |
| ttx00 | 564.2 | -1934 to 3063 | No | ns | 0.9959 | | | |
| ttx03 | 751.5 | -1688 to 3190 | No | ns | 0.9744 | | | |
| ttx06 | -3430 | -6180 to -678.9 | Yes | ** | 0.0058 | | | |
| ttx12 | -2037 | -4309 to 236.3 | No | ns | 0.1079 | | | |
| ttx24 | -4206 | -6379 to -2032 | Yes | **** | <0.0001 | | | |
| ttx48 | 2553 | 343.2 to 4764 | Yes | * | 0.0136 | | | |
| ttx72 | 973.5 | -1371 to 3318 | No | ns | 0.8833 | | | |
| **Test details** | **Predicted (LS** | **Predicted (LS) mea** | **Predicted (L** | **SE of diff.** | **N1** | **N2** | **t** | **DF** |
| wt - ts | | | | | | | | |
| ttx00 | 10530 | 9966 | 564.2 | 928.9 | 46 | 51 | 0.6074 | 761.0 |
| ttx03 | 9520 | 8768 | 751.5 | 906.7 | 60 | 44 | 0.8288 | 761.0 |
| ttx06 | 12243 | 15673 | -3430 | 1023 | 29 | 64 | 3.354 | 761.0 |
| ttx12 | 10334 | 12371 | -2037 | 844.9 | 57 | 60 | 2.410 | 761.0 |
| ttx24 | 11022 | 15227 | -4206 | 807.9 | 66 | 62 | 5.205 | 761.0 |
| ttx48 | 14895 | 12342 | 2553 | 821.6 | 69 | 56 | 3.108 | 761.0 |
| ttx72 | 10579 | 9605 | 973.5 | 871.5 | 61 | 50 | 1.117 | 761.0 |

| Test details | Predicted ( | Predicted (LS) mea | Predicted | SE of diff. | N1 | N2 | q | DF | Predicted ( | 95.00% CI of diff. | Below thre | Summary | Adjusted F |
|---|---|---|---|---|---|---|---|---|---|---|---|---|---|
| wt | | | | | | | | | | | | | |
| ttx00 vs. ttx03 | 10530 | 9520 | 1010 | 895.2 | 46 | 60 | 1.596 | 761.0 | 1010 | -1636 to 3657 | No | ns | 0.9193 |
| ttx00 vs. ttx06 | 10530 | 12243 | -1713 | 1083 | 46 | 29 | 2.237 | 761.0 | -1713 | -4915 to 1489 | No | ns | 0.6943 |
| ttx00 vs. ttx12 | 10530 | 10334 | 195.8 | 905.4 | 46 | 57 | 0.3058 | 761.0 | 195.8 | -2481 to 2872 | No | ns | >0.9999 |
| ttx00 vs. ttx24 | 10530 | 11022 | -491.9 | 877.4 | 46 | 66 | 0.7928 | 761.0 | -491.9 | -3086 to 2102 | No | ns | 0.9978 |
| ttx00 vs. ttx48 | 10530 | 14895 | -4365 | 869.5 | 46 | 69 | 7.100 | 761.0 | -4365 | -6936 to -1795 | Yes | **** | <0.0001 |
| ttx00 vs. ttx72 | 10530 | 10579 | -48.87 | 892.0 | 46 | 61 | 0.07747 | 761.0 | -48.87 | -2686 to 2588 | No | ns | >0.9999 |
| ttx03 vs. ttx06 | 9520 | 12243 | -2723 | 1033 | 60 | 29 | 3.728 | 761.0 | -2723 | -5777 to 331.0 | No | ns | 0.1168 |
| ttx03 vs. ttx12 | 9520 | 10334 | -814.4 | 844.9 | 60 | 57 | 1.363 | 761.0 | -814.4 | -3312 to 1683 | No | ns | 0.9615 |
| ttx03 vs. ttx24 | 9520 | 11022 | -1502 | 814.8 | 60 | 66 | 2.607 | 761.0 | -1502 | -3911 to 906.8 | No | ns | 0.5190 |
| ttx03 vs. ttx48 | 9520 | 14895 | -5376 | 806.4 | 60 | 69 | 9.428 | 761.0 | -5376 | -7759 to -2992 | Yes | **** | <0.0001 |
| ttx03 vs. ttx72 | 9520 | 10579 | -1059 | 830.6 | 60 | 61 | 1.803 | 761.0 | -1059 | -3514 to 1396 | No | ns | 0.8633 |
| ttx06 vs. ttx12 | 12243 | 10334 | 1909 | 1042 | 29 | 57 | 2.591 | 761.0 | 1909 | -1171 to 4989 | No | ns | 0.5267 |
| ttx06 vs. ttx24 | 12243 | 11022 | 1221 | 1018 | 29 | 66 | 1.697 | 761.0 | 1221 | -1787 to 4230 | No | ns | 0.8942 |
| ttx06 vs. ttx48 | 12243 | 14895 | -2652 | 1011 | 29 | 69 | 3.710 | 761.0 | -2652 | -5641 to 336.2 | No | ns | 0.1203 |
| ttx06 vs. ttx72 | 12243 | 10579 | 1664 | 1030 | 29 | 61 | 2.284 | 761.0 | 1664 | -1382 to 4710 | No | ns | 0.6726 |
| ttx12 vs. ttx24 | 10334 | 11022 | -687.6 | 826.0 | 57 | 66 | 1.177 | 761.0 | -687.6 | -3129 to 1754 | No | ns | 0.9815 |
| ttx12 vs. ttx48 | 10334 | 14895 | -4561 | 817.6 | 57 | 69 | 7.889 | 761.0 | -4561 | -6978 to -2144 | Yes | **** | <0.0001 |
| ttx12 vs. ttx72 | 10334 | 10579 | -244.6 | 841.5 | 57 | 61 | 0.4111 | 761.0 | -244.6 | -2732 to 2243 | No | ns | >0.9999 |
| ttx24 vs. ttx48 | 11022 | 14895 | -3873 | 786.5 | 66 | 69 | 6.965 | 761.0 | -3873 | -6199 to -1548 | Yes | **** | <0.0001 |
| ttx24 vs. ttx72 | 11022 | 10579 | 443.0 | 811.3 | 66 | 61 | 0.7722 | 761.0 | 443.0 | -1955 to 2841 | No | ns | 0.9981 |
| ttx48 vs. ttx72 | 14895 | 10579 | 4317 | 802.8 | 69 | 61 | 7.604 | 761.0 | 4317 | 1943 to 6690 | Yes | **** | <0.0001 |
| ts | | | | | | | | | | | | | |
| ttx00 vs. ttx03 | 9966 | 8768 | 1197 | 939.9 | 51 | 44 | 1.802 | 761.0 | 1197 | -1581 to 3976 | No | ns | 0.8638 |
| ttx00 vs. ttx06 | 9966 | 15673 | -5707 | 857.4 | 51 | 64 | 9.413 | 761.0 | -5707 | -8242 to -3172 | Yes | **** | <0.0001 |
| ttx00 vs. ttx12 | 9966 | 12371 | -2405 | 870.0 | 51 | 60 | 3.909 | 761.0 | -2405 | -4977 to 167.0 | No | ns | 0.0844 |
| ttx00 vs. ttx24 | 9966 | 15227 | -5262 | 863.6 | 51 | 62 | 8.617 | 761.0 | -5262 | -7815 to -2709 | Yes | **** | <0.0001 |
| ttx00 vs. ttx48 | 9966 | 12342 | -2376 | 884.2 | 51 | 56 | 3.801 | 761.0 | -2376 | -4990 to 237.6 | No | ns | 0.1028 |
| ttx00 vs. ttx72 | 9966 | 9605 | 360.4 | 909.1 | 51 | 50 | 0.5607 | 761.0 | 360.4 | -2327 to 3048 | No | ns | 0.9997 |
| ttx03 vs. ttx06 | 8768 | 15673 | -6904 | 894.6 | 44 | 64 | 10.91 | 761.0 | -6904 | -9549 to -4260 | Yes | **** | <0.0001 |
| ttx03 vs. ttx12 | 8768 | 12371 | -3602 | 906.7 | 44 | 60 | 5.619 | 761.0 | -3602 | -6283 to -922.1 | Yes | ** | 0.0015 |
| ttx03 vs. ttx24 | 8768 | 15227 | -6459 | 900.5 | 44 | 62 | 10.14 | 761.0 | -6459 | -9121 to -3797 | Yes | **** | <0.0001 |
| ttx03 vs. ttx48 | 8768 | 12342 | -3574 | 920.3 | 44 | 56 | 5.492 | 761.0 | -3574 | -6294 to -853.1 | Yes | ** | 0.0021 |
| ttx03 vs. ttx72 | 8768 | 9605 | -837.0 | 944.2 | 44 | 50 | 1.254 | 761.0 | -837.0 | -3628 to 1954 | No | ns | 0.9746 |
| ttx06 vs. ttx12 | 15673 | 12371 | 3302 | 820.9 | 64 | 60 | 5.688 | 761.0 | 3302 | 875.1 to 5729 | Yes | ** | 0.0012 |
| ttx06 vs. ttx24 | 15673 | 15227 | 445.2 | 814.0 | 64 | 62 | 0.7734 | 761.0 | 445.2 | -1961 to 2852 | No | ns | 0.9981 |
| ttx06 vs. ttx48 | 15673 | 12342 | 3331 | 835.9 | 64 | 56 | 5.635 | 761.0 | 3331 | 859.6 to 5802 | Yes | ** | 0.0014 |
| ttx06 vs. ttx72 | 15673 | 9605 | 6067 | 862.2 | 64 | 50 | 9.952 | 761.0 | 6067 | 3518 to 8616 | Yes | **** | <0.0001 |
| ttx12 vs. ttx24 | 12371 | 15227 | -2857 | 827.3 | 60 | 62 | 4.884 | 761.0 | -2857 | -5302 to -411.1 | Yes | * | 0.0104 |
| ttx12 vs. ttx48 | 12371 | 12342 | 28.77 | 848.8 | 60 | 56 | 0.04793 | 761.0 | 28.77 | -2480 to 2538 | No | ns | >0.9999 |
| ttx12 vs. ttx72 | 12371 | 9605 | 2765 | 874.7 | 60 | 50 | 4.471 | 761.0 | 2765 | 179.6 to 5351 | Yes | * | 0.0271 |
| ttx24 vs. ttx48 | 15227 | 12342 | 2885 | 842.1 | 62 | 56 | 4.846 | 761.0 | 2885 | 395.9 to 5375 | Yes | * | 0.0115 |
| ttx24 vs. ttx72 | 15227 | 9605 | 5622 | 868.3 | 62 | 50 | 9.157 | 761.0 | 5622 | 3055 to 8189 | Yes | **** | <0.0001 |
| ttx48 vs. ttx72 | 12342 | 9605 | 2737 | 888.8 | 56 | 50 | 4.354 | 761.0 | 2737 | 109.2 to 5364 | Yes | * | 0.0349 |

### Table S7D: Statistics for Figure 5 - βCaMKII dendritic shaft

| Source of Variation | % of total variation | P value | P value sum | Significant? | |
|---|---|---|---|---|---|
| Interaction | 7.149 | <0.0001 | **** | Yes | |
| Hours TTX | 17.00 | <0.0001 | **** | Yes | |
| Genotype | 0.4346 | 0.0332 | * | Yes | |
| **ANOVA table** | **SS (Type III)** | **DF** | **MS** | **F (DFn, DFd)** | **P value** |
| Interaction | 463470794 | 6 | 77245132 | F (6, 784) = 12.47 | P<0.0001 |
| Hours TTX | 1101995476 | 6 | 183665913 | F (6, 784) = 29.66 | P<0.0001 |
| Genotype | 28174597 | 1 | 28174597 | F (1, 784) = 4.549 | P=0.0332 |
| Residual | 4855541355 | 784 | 6193293 | | |
| **Difference between column means** | | | | | |
| Predicted (LS) mean of wt | 8476 | | | | |
| Predicted (LS) mean of ts | 8097 | | | | |
| Difference between predicted means | 379.0 | | | | |
| SE of difference | 177.7 | | | | |
| 95% CI of difference | 30.19 to 727.8 | | | | |
| **Data summary** | | | | | |
| Number of columns (Genotype) | 2 | | | | |
| Number of rows (Hours TTX) | 7 | | | | |
| Number of values | 798 | | | | |

| Šídák's multiple | Predicted (LS) | 95.00% CI of dif | Below thre | Summary | Adjusted P | | | |
|---|---|---|---|---|---|---|---|---|
| wt - ts | | | | | | | | |
| ttx00 | 639.8 | -721.4 to 2001 | No | ns | 0.8019 | | | |
| ttx03 | 1717 | 388.9 to 3046 | Yes | ** | 0.0037 | | | |
| ttx06 | -433.7 | -1683 to 816.1 | No | ns | 0.9514 | | | |
| ttx12 | 801.5 | -436.7 to 2040 | No | ns | 0.4508 | | | |
| ttx24 | -2516 | -3700 to -1332 | Yes | **** | <0.0001 | | | |
| ttx48 | 2368 | 1164 to 3572 | Yes | **** | <0.0001 | | | |
| ttx72 | 76.30 | -1201 to 1353 | No | ns | >0.9999 | | | |
| **Test details** | **Predicted (LS)** | **Predicted (LS) i** | **Predicted** | **SE of diff.** | **N1** | **N2** | **t** | **DF** |
| wt - ts | | | | | | | | |
| ttx00 | 7217 | 6578 | 639.8 | 506.0 | 46 | 51 | 1.264 | 784.0 |
| ttx03 | 8033 | 6316 | 1717 | 493.9 | 60 | 44 | 3.477 | 784.0 |
| ttx06 | 9568 | 10001 | -433.7 | 464.6 | 52 | 64 | 0.9334 | 784.0 |
| ttx12 | 8212 | 7410 | 801.5 | 460.3 | 57 | 60 | 1.741 | 784.0 |
| ttx24 | 7641 | 10157 | -2516 | 440.1 | 66 | 62 | 5.716 | 784.0 |
| ttx48 | 11218 | 8850 | 2368 | 447.6 | 69 | 56 | 5.290 | 784.0 |
| ttx72 | 7441 | 7365 | 76.30 | 474.8 | 61 | 50 | 0.1607 | 784.0 |

| Šídák's multiple co | Predicted | 95.00% CI of diff. | Below thre | Summary | Adjusted P | Predicted | Predicted (LS) me | Predicted | SE of diff. | N1 | N2 | t | DF |
|---|---|---|---|---|---|---|---|---|---|---|---|---|---|
| wt | | | | | | | | | | | | | |
| ttx00 vs. ttx03 | -815.8 | -2299 to 667.1 | No | ns | 0.8764 | 7217 | 8033 | -815.8 | 487.7 | 46 | 60 | 1.673 | 784.0 |
| ttx00 vs. ttx06 | -2350 | -3882 to -818.5 | Yes | **** | <0.0001 | 7217 | 9568 | -2350 | 503.7 | 46 | 52 | 4.666 | 784.0 |
| ttx00 vs. ttx12 | -994.1 | -2494 to 505.6 | No | ns | 0.6130 | 7217 | 8212 | -994.1 | 493.2 | 46 | 57 | 2.015 | 784.0 |
| ttx00 vs. ttx24 | -423.9 | -1877 to 1029 | No | ns | >0.9999 | 7217 | 7641 | -423.9 | 478.0 | 46 | 66 | 0.8868 | 784.0 |
| ttx00 vs. ttx48 | -4000 | -5441 to -2560 | Yes | **** | <0.0001 | 7217 | 11218 | -4000 | 473.7 | 46 | 69 | 8.445 | 784.0 |
| ttx00 vs. ttx72 | -223.7 | -1701 to 1254 | No | ns | >0.9999 | 7217 | 7441 | -223.7 | 486.0 | 46 | 61 | 0.4602 | 784.0 |
| ttx03 vs. ttx06 | -1534 | -2968 to -100.6 | Yes | * | 0.0246 | 8033 | 9568 | -1534 | 471.5 | 60 | 52 | 3.254 | 784.0 |
| ttx03 vs. ttx12 | -178.3 | -1578 to 1221 | No | ns | >0.9999 | 8033 | 8212 | -178.3 | 460.3 | 60 | 57 | 0.3873 | 784.0 |
| ttx03 vs. ttx24 | 392.0 | -957.8 to 1742 | No | ns | >0.9999 | 8033 | 7641 | 392.0 | 443.9 | 60 | 66 | 0.8829 | 784.0 |
| ttx03 vs. ttx48 | -3185 | -4520 to -1849 | Yes | **** | <0.0001 | 8033 | 11218 | -3185 | 439.3 | 60 | 69 | 7.249 | 784.0 |
| ttx03 vs. ttx72 | 592.2 | -783.7 to 1968 | No | ns | 0.9883 | 8033 | 7441 | 592.2 | 452.5 | 60 | 61 | 1.309 | 784.0 |
| ttx06 vs. ttx12 | 1356 | -95.04 to 2807 | No | ns | 0.0924 | 9568 | 8212 | 1356 | 477.2 | 52 | 57 | 2.841 | 784.0 |
| ttx06 vs. ttx24 | 1926 | 523.2 to 3329 | Yes | *** | 0.0007 | 9568 | 7641 | 1926 | 461.5 | 52 | 66 | 4.174 | 784.0 |
| ttx06 vs. ttx48 | -1650 | -3040 to -260.6 | Yes | ** | 0.0068 | 9568 | 11218 | -1650 | 457.0 | 52 | 69 | 3.611 | 784.0 |
| ttx06 vs. ttx72 | 2127 | 698.3 to 3555 | Yes | *** | 0.0001 | 9568 | 7441 | 2127 | 469.7 | 52 | 61 | 4.527 | 784.0 |
| ttx12 vs. ttx24 | 570.2 | -798.0 to 1938 | No | ns | 0.9920 | 8212 | 7641 | 570.2 | 450.0 | 57 | 66 | 1.267 | 784.0 |
| ttx12 vs. ttx48 | -3006 | -4361 to -1652 | Yes | **** | <0.0001 | 8212 | 11218 | -3006 | 445.4 | 57 | 69 | 6.749 | 784.0 |
| ttx12 vs. ttx72 | 770.5 | -623.5 to 2164 | No | ns | 0.8720 | 8212 | 7441 | 770.5 | 458.5 | 57 | 61 | 1.681 | 784.0 |
| ttx24 vs. ttx48 | -3577 | -4879 to -2274 | Yes | **** | <0.0001 | 7641 | 11218 | -3577 | 428.5 | 66 | 69 | 8.347 | 784.0 |
| ttx24 vs. ttx72 | 200.2 | -1144 to 1544 | No | ns | >0.9999 | 7641 | 7441 | 200.2 | 442.0 | 66 | 61 | 0.4530 | 784.0 |
| ttx48 vs. ttx72 | 3777 | 2447 to 5107 | Yes | **** | <0.0001 | 11218 | 7441 | 3777 | 437.4 | 69 | 61 | 8.635 | 784.0 |
| ts | | | | | | | | | | | | | |
| ttx00 vs. ttx03 | 261.9 | -1295 to 1819 | No | ns | >0.9999 | 6578 | 6316 | 261.9 | 512.0 | 51 | 44 | 0.5114 | 784.0 |
| ttx00 vs. ttx06 | -3424 | -4844 to -2003 | Yes | **** | <0.0001 | 6578 | 10001 | -3424 | 467.1 | 51 | 64 | 7.329 | 784.0 |
| ttx00 vs. ttx12 | -832.4 | -2274 to 608.8 | No | ns | 0.8242 | 6578 | 7410 | -832.4 | 474.0 | 51 | 60 | 1.756 | 784.0 |
| ttx00 vs. ttx24 | -3580 | -5010 to -2149 | Yes | **** | <0.0001 | 6578 | 10157 | -3580 | 470.5 | 51 | 62 | 7.609 | 784.0 |
| ttx00 vs. ttx48 | -2272 | -3737 to -807.8 | Yes | **** | <0.0001 | 6578 | 8850 | -2272 | 481.7 | 51 | 56 | 4.718 | 784.0 |
| ttx00 vs. ttx72 | -787.1 | -2293 to 718.8 | No | ns | 0.9183 | 6578 | 7365 | -787.1 | 495.3 | 51 | 50 | 1.589 | 784.0 |
| ttx03 vs. ttx06 | -3685 | -5167 to -2204 | Yes | **** | <0.0001 | 6316 | 10001 | -3685 | 487.4 | 44 | 64 | 7.562 | 784.0 |
| ttx03 vs. ttx12 | -1094 | -2596 to 407.6 | No | ns | 0.4374 | 6316 | 7410 | -1094 | 493.9 | 44 | 60 | 2.215 | 784.0 |
| ttx03 vs. ttx24 | -3841 | -5333 to -2350 | Yes | **** | <0.0001 | 6316 | 10157 | -3841 | 490.6 | 44 | 62 | 7.831 | 784.0 |
| ttx03 vs. ttx48 | -2534 | -4059 to -1010 | Yes | **** | <0.0001 | 6316 | 8850 | -2534 | 501.3 | 44 | 56 | 5.055 | 784.0 |
| ttx03 vs. ttx72 | -1049 | -2613 to 515.2 | No | ns | 0.5918 | 6316 | 7365 | -1049 | 514.4 | 44 | 50 | 2.039 | 784.0 |
| ttx06 vs. ttx12 | 2591 | 1231 to 3951 | Yes | **** | <0.0001 | 10001 | 7410 | 2591 | 447.2 | 64 | 60 | 5.794 | 784.0 |
| ttx06 vs. ttx24 | -156.0 | -1504 to 1192 | No | ns | >0.9999 | 10001 | 10157 | -156.0 | 443.5 | 64 | 62 | 0.3517 | 784.0 |
| ttx06 vs. ttx48 | 1151 | -233.4 to 2536 | No | ns | 0.2184 | 10001 | 8850 | 1151 | 455.4 | 64 | 56 | 2.528 | 784.0 |
| ttx06 vs. ttx72 | 2637 | 1208 to 4065 | Yes | **** | <0.0001 | 10001 | 7365 | 2637 | 469.7 | 64 | 50 | 5.613 | 784.0 |
| ttx12 vs. ttx24 | -2747 | -4118 to -1377 | Yes | **** | <0.0001 | 7410 | 10157 | -2747 | 450.7 | 60 | 62 | 6.096 | 784.0 |
| ttx12 vs. ttx48 | -1440 | -2846 to -34.06 | Yes | * | 0.0394 | 7410 | 8850 | -1440 | 462.4 | 60 | 56 | 3.114 | 784.0 |
| ttx12 vs. ttx72 | 45.30 | -1404 to 1494 | No | ns | >0.9999 | 7410 | 7365 | 45.30 | 476.5 | 60 | 50 | 0.09505 | 784.0 |
| ttx24 vs. ttx48 | 1307 | -87.85 to 2702 | No | ns | 0.0903 | 10157 | 8850 | 1307 | 458.8 | 62 | 56 | 2.849 | 784.0 |
| ttx24 vs. ttx72 | 2792 | 1354 to 4231 | Yes | **** | <0.0001 | 10157 | 7365 | 2792 | 473.0 | 62 | 50 | 5.903 | 784.0 |
| ttx48 vs. ttx72 | 1485 | 13.05 to 2958 | Yes | * | 0.0458 | 8850 | 7365 | 1485 | 484.2 | 56 | 50 | 3.068 | 784.0 |

### Table S7E: Statistics for Figure 5 - αCaMKII synaptic

| Source of Variation | % of total variation | P value | P value summ | Significant? | |
|---|---|---|---|---|---|
| Interaction | 16.64 | <0.0001 | **** | Yes | |
| Hours TTX | 18.26 | <0.0001 | **** | Yes | |
| Genotype | 1.606 | 0.0002 | *** | Yes | |
| **ANOVA table** | **SS (Type III)** | **DF** | **MS** | **F (DFn, DFd)** | **P value** |
| Interaction | 3880549404 | 6 | 646758234 | F (6, 553) = 24.09 | P<0.0001 |
| Hours TTX | 4258407086 | 6 | 709734514 | F (6, 553) = 26.43 | P<0.0001 |
| Genotype | 374596303 | 1 | 374596303 | F (1, 553) = 13.95 | P=0.0002 |
| Residual | 14847781752 | 553 | 26849515 | | |
| **Difference between column means** | | | | | |
| Predicted (LS) mean of wt | 10281 | | | | |
| Predicted (LS) mean of tis | 8647 | | | | |
| Difference between predicted means | 1634 | | | | |
| SE of difference | 437.4 | | | | |
| 95% CI of difference | 774.7 to 2493 | | | | |
| **Data summary** | | | | | |
| Number of columns (Genotype) | 2 | | | | |
| Number of rows (Hours TTX) | 7 | | | | |
| Number of values | 567 | | | | |

| Šidák's multiple | Predicted | 95.00% CI | Below thre | Summary | Adjusted F | | | |
|---|---|---|---|---|---|---|---|---|
| wt - tis | | | | | | | | |
| ttx00 | 401.0 | -2480 to 32 | No | ns | 0.9998 | | | |
| ttx03 | 2546 | -482.7 to 5 | No | ns | 0.1563 | | | |
| ttx06 | -8319 | -11461 to - | Yes | **** | <0.0001 | | | |
| ttx12 | 4073 | 687.8 to 74 | Yes | ** | 0.0088 | | | |
| ttx24 | 4576 | 1528 to 76 | Yes | *** | 0.0004 | | | |
| ttx48 | -1914 | -4997 to 11 | No | ns | 0.5031 | | | |
| ttx72 | 10074 | 6851 to 13 | Yes | **** | <0.0001 | | | |
| **Test details** | **Predicted** | **Predicted** | **Predicted** | **SE of diff.** | **N1** | **N2** | **t** | **DF** |
| wt - tis | | | | | | | | |
| ttx00 | 13856 | 13455 | 401.0 | 1070 | 49 | 45 | 0.3748 | 553.0 |
| ttx03 | 7453 | 4907 | 2546 | 1125 | 41 | 44 | 2.264 | 553.0 |
| ttx06 | 4520 | 12839 | -8319 | 1167 | 41 | 38 | 7.130 | 553.0 |
| ttx12 | 10488 | 6415 | 4073 | 1257 | 33 | 35 | 3.240 | 553.0 |
| ttx24 | 9114 | 4538 | 4576 | 1132 | 44 | 40 | 4.043 | 553.0 |
| ttx48 | 11996 | 13910 | -1914 | 1145 | 42 | 40 | 1.672 | 553.0 |
| ttx72 | 14539 | 4465 | 10074 | 1197 | 38 | 37 | 8.417 | 553.0 |

| Tukey's multiple c | Predicted | 95.00% CI of di | Below thre | Summary | Adjusted F | Predicted | Predicted (LS) | Predicted | SE of diff. | N1 | N2 | q | DF |
|---|---|---|---|---|---|---|---|---|---|---|---|---|---|
| wt | | | | | | | | | | | | | |
| ttx00 vs. ttx03 | 6403 | 3157 to 9648 | Yes | **** | <0.0001 | 13856 | 7453 | 6403 | 1097 | 49 | 41 | 8.256 | 553.0 |
| ttx00 vs. ttx06 | 9336 | 6090 to 12581 | Yes | **** | <0.0001 | 13856 | 4520 | 9336 | 1097 | 49 | 41 | 12.04 | 553.0 |
| ttx00 vs. ttx12 | 3367 | -85.61 to 6820 | No | ns | 0.0614 | 13856 | 10488 | 3367 | 1167 | 49 | 33 | 4.081 | 553.0 |
| ttx00 vs. ttx24 | 4741 | 1557 to 7926 | Yes | *** | 0.0003 | 13856 | 9114 | 4741 | 1076 | 49 | 44 | 6.231 | 553.0 |
| ttx00 vs. ttx48 | 1860 | -1365 to 5084 | No | ns | 0.6118 | 13856 | 11996 | 1860 | 1090 | 49 | 42 | 2.414 | 553.0 |
| ttx00 vs. ttx72 | -682.7 | -3997 to 2632 | No | ns | 0.9965 | 13856 | 14539 | -682.7 | 1120 | 49 | 38 | 0.8620 | 553.0 |
| ttx03 vs. ttx06 | 2933 | -453.6 to 6320 | No | ns | 0.1396 | 7453 | 4520 | 2933 | 1144 | 41 | 41 | 3.625 | 553.0 |
| ttx03 vs. ttx12 | -3035 | -6622 to 550.6 | No | ns | 0.1594 | 7453 | 10488 | -3035 | 1212 | 41 | 33 | 3.542 | 553.0 |
| ttx03 vs. ttx24 | -1661 | -4990 to 1667 | No | ns | 0.7584 | 7453 | 9114 | -1661 | 1125 | 41 | 44 | 2.089 | 553.0 |
| ttx03 vs. ttx48 | -4543 | -7910 to -1177 | Yes | ** | 0.0014 | 7453 | 11996 | -4543 | 1138 | 41 | 42 | 5.648 | 553.0 |
| ttx03 vs. ttx72 | -7086 | -10538 to -3633 | Yes | **** | <0.0001 | 7453 | 14539 | -7086 | 1167 | 41 | 38 | 8.588 | 553.0 |
| ttx06 vs. ttx12 | -5969 | -9555 to -2383 | Yes | **** | <0.0001 | 4520 | 10488 | -5969 | 1212 | 41 | 33 | 6.965 | 553.0 |
| ttx06 vs. ttx24 | -4595 | -7923 to -1266 | Yes | *** | 0.0010 | 4520 | 9114 | -4595 | 1125 | 41 | 44 | 5.777 | 553.0 |
| ttx06 vs. ttx48 | -7476 | -10843 to -4110 | Yes | **** | <0.0001 | 4520 | 11996 | -7476 | 1138 | 41 | 42 | 9.294 | 553.0 |
| ttx06 vs. ttx72 | -10019 | -13471 to -6566 | Yes | **** | <0.0001 | 4520 | 14539 | -10019 | 1167 | 41 | 38 | 12.14 | 553.0 |
| ttx12 vs. ttx24 | 1374 | -2157 to 4905 | No | ns | 0.9116 | 10488 | 9114 | 1374 | 1193 | 33 | 44 | 1.628 | 553.0 |
| ttx12 vs. ttx48 | -1508 | -5075 to 2059 | No | ns | 0.8737 | 10488 | 11996 | -1508 | 1205 | 33 | 42 | 1.769 | 553.0 |
| ttx12 vs. ttx72 | -4050 | -7699 to -401.5 | Yes | * | 0.0186 | 10488 | 14539 | -4050 | 1233 | 33 | 38 | 4.645 | 553.0 |
| ttx24 vs. ttx48 | -2882 | -6190 to 426.1 | No | ns | 0.1347 | 9114 | 11996 | -2882 | 1118 | 44 | 42 | 3.646 | 553.0 |
| ttx24 vs. ttx72 | -5424 | -8820 to -2028 | Yes | **** | <0.0001 | 9114 | 14539 | -5424 | 1148 | 44 | 38 | 6.685 | 553.0 |
| ttx48 vs. ttx72 | -2542 | -5975 to 890.6 | No | ns | 0.3017 | 11996 | 14539 | -2542 | 1160 | 42 | 38 | 3.099 | 553.0 |
| tis | | | | | | | | | | | | | |
| ttx00 vs. ttx03 | 8548 | 5297 to 11799 | Yes | **** | <0.0001 | 13455 | 4907 | 8548 | 1099 | 45 | 44 | 11.00 | 553.0 |
| ttx00 vs. ttx06 | 615.9 | -2762 to 3994 | No | ns | 0.9982 | 13455 | 12839 | 615.9 | 1142 | 45 | 38 | 0.7630 | 553.0 |
| ttx00 vs. ttx12 | 7040 | 3584 to 10496 | Yes | **** | <0.0001 | 13455 | 6415 | 7040 | 1168 | 45 | 35 | 8.525 | 553.0 |
| ttx00 vs. ttx24 | 8917 | 5585 to 12249 | Yes | **** | <0.0001 | 13455 | 4538 | 8917 | 1126 | 45 | 40 | 11.20 | 553.0 |
| ttx00 vs. ttx48 | -455.4 | -3788 to 2877 | No | ns | 0.9997 | 13455 | 13910 | -455.4 | 1126 | 45 | 40 | 0.5720 | 553.0 |
| ttx00 vs. ttx72 | 8990 | 5587 to 12393 | Yes | **** | <0.0001 | 13455 | 4465 | 8990 | 1150 | 45 | 37 | 11.06 | 553.0 |
| ttx03 vs. ttx06 | -7932 | -11328 to -4536 | Yes | **** | <0.0001 | 4907 | 12839 | -7932 | 1148 | 44 | 38 | 9.776 | 553.0 |
| ttx03 vs. ttx12 | -1508 | -4981 to 1965 | No | ns | 0.8587 | 4907 | 6415 | -1508 | 1174 | 44 | 35 | 1.817 | 553.0 |
| ttx03 vs. ttx24 | 368.9 | -2981 to 3719 | No | ns | >0.9999 | 4907 | 4538 | 368.9 | 1132 | 44 | 40 | 0.4609 | 553.0 |
| ttx03 vs. ttx48 | -9003 | -12353 to -5653 | Yes | **** | <0.0001 | 4907 | 13910 | -9003 | 1132 | 44 | 40 | 11.25 | 553.0 |
| ttx03 vs. ttx72 | 441.9 | -2978 to 3862 | No | ns | 0.9998 | 4907 | 4465 | 441.9 | 1156 | 44 | 37 | 0.5408 | 553.0 |
| ttx06 vs. ttx12 | 6424 | 2831 to 10016 | Yes | **** | <0.0001 | 12839 | 6415 | 6424 | 1214 | 38 | 35 | 7.484 | 553.0 |
| ttx06 vs. ttx24 | 8301 | 4827 to 11774 | Yes | **** | <0.0001 | 12839 | 4538 | 8301 | 1174 | 38 | 40 | 10.00 | 553.0 |
| ttx06 vs. ttx48 | -1071 | -4545 to 2402 | No | ns | 0.9705 | 12839 | 13910 | -1071 | 1174 | 38 | 40 | 1.291 | 553.0 |
| ttx06 vs. ttx72 | 8374 | 4832 to 11915 | Yes | **** | <0.0001 | 12839 | 4465 | 8374 | 1197 | 38 | 37 | 9.896 | 553.0 |
| ttx12 vs. ttx24 | 1877 | -1672 to 5426 | No | ns | 0.7047 | 6415 | 4538 | 1877 | 1199 | 35 | 40 | 2.213 | 553.0 |
| ttx12 vs. ttx48 | -7495 | -11044 to -3946 | Yes | **** | <0.0001 | 6415 | 13910 | -7495 | 1199 | 35 | 40 | 8.838 | 553.0 |
| ttx12 vs. ttx72 | 1950 | -1665 to 5566 | No | ns | 0.6850 | 6415 | 4465 | 1950 | 1222 | 35 | 37 | 2.257 | 553.0 |
| ttx24 vs. ttx48 | -9372 | -12801 to -5944 | Yes | **** | <0.0001 | 4538 | 13910 | -9372 | 1159 | 40 | 40 | 11.44 | 553.0 |
| ttx24 vs. ttx72 | 73.02 | -3425 to 3571 | No | ns | >0.9999 | 4538 | 4465 | 73.02 | 1182 | 40 | 37 | 0.08738 | 553.0 |
| ttx48 vs. ttx72 | 9445 | 5948 to 12943 | Yes | **** | <0.0001 | 13910 | 4465 | 9445 | 1182 | 40 | 37 | 11.30 | 553.0 |
